# MetENP/MetENPWeb: An R package and web application for metabolomics enrichment and pathway analysis in Metabolomics Workbench

**DOI:** 10.1101/2020.11.20.391912

**Authors:** Kumari Sonal Choudhary, Eoin Fahy, Kevin Coakley, Manish Sud, Mano R Maurya, Shankar Subramaniam

**Affiliations:** Departments of Bioengineering, Cellular & Molecular Medicine and Computer Science & Engineering, University of California San Diego, La Jolla, CA 92093 USA; San Diego Supercomputer Center, University of California, San Diego, 9500 Gilman Drive, La Jolla, CA 92037, USA

## Abstract

With the advent of high throughput mass spectrometric methods, metabolomics has emerged as an essential area of research in biomedicine with the potential to provide deep biological insights into normal and diseased functions in physiology. However, to achieve the potential offered by metabolomics measures, there is a need for biologist-friendly integrative analysis tools that can transform data into mechanisms that relate to phenotypes. Here, we describe MetENP, an R package, and a user-friendly web application deployed at the Metabolomics Workbench site extending the metabolomics enrichment analysis to include species-specific pathway analysis, pathway enrichment scores, gene-enzyme information, and enzymatic activities of the significantly altered metabolites. MetENP provides a highly customizable workflow through various user-specified options and includes support for all metabolite species with available KEGG pathways. MetENPweb is a web application for calculating metabolite and pathway enrichment analysis.

**Availability and Implementation:** The MetENP package is freely available from Metabolomics Workbench GitHub: (https://github.com/metabolomicsworkbench/MetENP), the web application, is freely available at (https://www.metabolomicsworkbench.org/data/analyze.php)

## INTRODUCTION

Metabolomics has evolved as a significant field within multi-omics systems biology to decipher mechanisms and predict cells, tissues, and whole organisms’ phenotypes. Metabolomics has broad applications for disease diagnosis, biomarker discovery, and precision medicine (1, 2). The number of publicly available metabolomics data sets is growing rapidly. Hence, there is a need to develop customizable tools and workflows that are biologist-friendly and capable of providing mechanistic insights into the system’s behavior (3). Several databases and tools are currently available in metabolomics; for instance, KEGG (4, 5), and MetaboAnalyst 4.0 (6) provides functional annotation of metabolites, and MetaboLights (7), MetaboAnalyst 4.0 (6), ChemRICH (8), and MetExplore (9) provides enrichment analysis of metabolites in the context of function or pathways. However, these tools warrant manipulations of data to perform analysis across various stages, a task that is challenging for an experimental researcher. It would be desirable to have a single comprehensive platform for metabolomics analysis from data deposition to functional annotation, which would significantly enhance the use of metabolomics in deciphering mechanistic biology.

The Metabolomics Workbench (MW), the National Metabolomics Data Repository (NMDR) resource, provides the biomedical research community with a compendium of metabolomics data sets along with a host of tools and user-friendly interfaces. MW delivers the community with capabilities to upload and analyze data seamlessly, linking metabolites to well-defined structures and spectra while offering the ability to perform extensive statistical analysis (10). MW can retrieve data from the database using REpresentational State Transfer (REST) services via HTTP requests. MW also encompasses databases such as RefMet (A <u>Ref</u>erence list of <u>Met</u>abolite names (10), that provides standard nomenclature for metabolites, enabling efficient comparisons between metabolite data across experiments. With MW as a foundation, we developed a workflow, MetENP, that extends the capability of metabolomics enrichment analysis within the MW to include pathway association of enriched metabolites and functional annotation in a species-specific manner. Analysis with MetENP enables a researcher to obtain insights into metabolites altered in comparative measurements, in terms of the metabolite category, the associated biological pathways, and the genes associated with the reactions along with associated enzymatic information. The MetENP R workflow is also available as a web application on Metabolomics Workbench via a user-friendly interface (MetENPWeb).

## IMPLEMENTATION AND FEATURES OF MetENP/MetENPWeb

MetENP is implemented in R, an open-source programming environment. The R package’s current version is available on the GitHub repository (https://github.com/metabolomicsworkbench/MetENP). Users can directly install the R package on their computer. Alternatively, users can run the analysis through the Jupyter notebook. The mybinder.org service can be used to run the Jupyter notebook for free on the web without having to install any software. A link to run the Jupyter notebook on mybinder.org is in the README.md on the GitHub repository. MetENPWeb is available on the Metabolomics Workbench (https://www.metabolomicsworkbench.org/data/analyze.php). It follows the workflow shown in Figure 1. The REST API is used to retrieve metabolomics data from Metabolomics Workbench. The KEGG REST API is employed to get KEGG data for reactions, pathways, and genes associated with metabolites.

**Figure 1:**
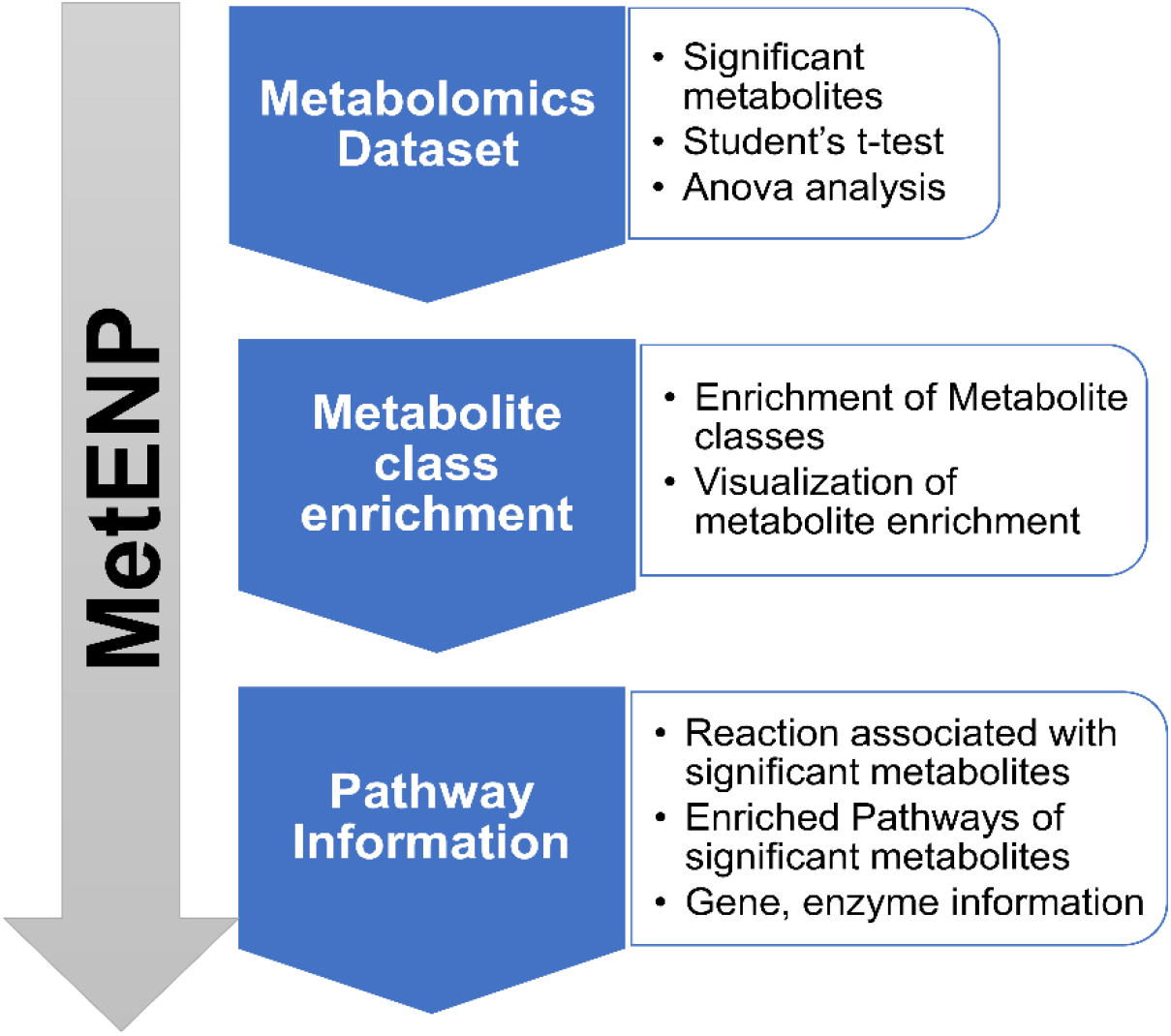
MetENP workflow. MetENP workflow shows the detection of significantly altered metabolites, mapping to metabolite class categories, calculation of metabolite enrichment score, association to KEGG pathways, and generation of functional annotation on significantly altered metabolites. The workflow supports visualization plots at each step.

The following sections provide details for MetENP workflow along with a description of available parameters to customize the workflow:

a. **User input:** The workflow requires metabolomics data and metadata. Users can invoke the metabolomics data and metadata directly from Metabolomics Workbench via study ID for MetENP analysis. Alternatively, users can upload their metabolomics data in a simple tab-delimited format accepted by MetENP. MetENP can take either of the two types of data structures: i) metabolite names are present in the first column. The column names should be the sample names. The second row should have information about experimental factors, e.g., age, disease, condition, time, etc.; and ii) sample names are present in the first column. The second column should include experimental factors, and subsequent columns should have metabolite measurements. (See supplementary file 1 for an example run; also available from https://www.metabolomicsworkbench.org/data/file-upload.php).
b. **Conversion to Refmet metabolite classification:** The *convert_refmet* function within MetENP assigns the RefMet metabolite class category (sub_class, super_class, and main_class) to the metabolites. Refmet is a database containing a standardized nomenclature for metabolite molecules.
c. **Calculation of significant metabolites:** The *significant_met* function with MetENP is used to calculate significant metabolites and rely on the Student’s *t*-test to analyze each metabolite based on chosen experimental groups or factors. Its output provides fold change, log2 fold change, p-value and adjusted p-value (options include Holm (“holm”), Hochberg (“hochberg”), Hommel (“hommel”), and Benjamini & Hochberg (“BH” or its alias “fdr”)) for each metabolite. Missing values are imputed, when appropriate, in the data preprocessing or filtering step. Three filtering methods have been incorporated in the package: a) *half_of_min:* where the NAs are replaced by half of the min values in the data, b) *remove_NAs:* where metabolites with NA values are removed and c) *50percent*: where metabolites with more than 50% NA values are removed. The users may adjust p-values and log2 fold change values to get the list of significantly altered metabolites. Further, *the anova_ana* function is used to analyze independent variables. Significant metabolites are visualized as a volcano plot using *plot_volcano* function within MetENP.
d. **Metabolite class enrichment:** To investigate the enrichment of metabolite classes in significantly altered metabolites, a hypergeometric (HG) test is performed using the *metclassenrichment* function of MetENP based on *phyper* function in R. The formula for the phyper calculation is *phyper (M-1, L, N-L, k, lower. tail=FALSE)*, where N <-all metabolites detected in a study, L <-all significant metabolites detected in a study, M <-all significant metabolites detected in a metabolite class and k <-all metabolites detected in a metabolite class. This gives a p-value of the hypergeometric test for each significantly altered metabolite.
e. **Pathway association:** MetENP provides functionality to link significantly altered metabolites to species-specific KEGG pathways via *met_pathways* function within the MetENP package. *met_pathways* function links metabolites to KEGG reactions and is further associated with KEGG pathways. Only metabolites with linked KEGG reactions are considered for pathway analysis. Pathway analysis can be carried out at both species level and reference pathway level.
f. **Pathway enrichment analysis:** MetENP calculates the pathway enrichment of the associated KEGG pathways using the HG test with *path enrichmentscore* function. This function also utilizes *phyper* function in R using a similar formula as for metabolite class enrichment. This provides a list of pathways and their respective hypergeometric p-values.
g. **Visualization:** MetENP supports the following plotting options: 1) volcano plot of the significant metabolites, 2) bar plots for the metabolite counts, 3) metabolite enrichment score in each metabolite class 4) pathway network 5) heatmap, and 6) dot plot.
h. **Gene information:** The *enzyme_gene_info* function within MetENP can retrieve genes and enzymes involved in the associated pathways. The *react_substrate* function provid**es** information on whether the metabolite is a reactant or a product in the associated reaction.

## WEB APPLICATION DESCRIPTION

The MetENPWeb on MW provides a user interface for analysis of the metabolomics dataset. After selecting the study ID, the metabolomics dataset and metadata are retrieved from the Metabolomics Workbench. Users have the choice to choose from the analysis types and the experimental factor column. Further, the analysis parameters can be selected, where individual groups for comparison, p-value, and log2 fold change thresholds, filtering methods for treatment of NA values, the number of metabolites in a class, etc., can be specified from the list of options by the user. After the analysis is complete, all the visualization graphics and result files are available for download.

## CASE STUDIES

To illustrate the capabilities of MetENP, we performed an analysis of three metabolomics datasets deposited on Metabolomics Workbench with Study ID: ST000915 (11), Study ID: ST001308 (12), and Study ID: ST001140 (13).

### Case study 1 (ST000915): A nonalcoholic fatty liver disease (NAFLD) biomarker study

This was a lipidomics study dealing with detecting lipid metabolites, aqueous intracellular metabolites, SNPs, and mRNA transcripts in patients at different stages of Fatty Liver Disease. The study was done in the liver, plasma, and urine. This example study has data from the liver.

We ran MetENP on this dataset to determine significantly altered metabolites between Normal and Cirrhosis samples and the pathways associated with them. The preprocessing step was applied to handle missing data from the dataset, where metabolites with more than 50% NA values are removed (50percent). We found 85 metabolites to be significantly altered with a p-value cut off 0.05, and log2 fold change cut off 0.5 (**Figure 2(A)**). Using MetENP, we associated these 85 significantly altered metabolites to metabolite classes, using the RefMet classification of Metabolomics Workbench. The metabolite class enrichment on significantly altered metabolites (**Figure 2(B)**), suggested that the several metabolites ranging from Eicosanoids, Ceramides, Triradylglycerols (TAGs) and Cardiolipins metabolite class were elevated in normal and those from Glycerophosphoserines, Cholesterol, and some metabolites of Glycerophospholipids class were elevated in cirrhosis. This result correlated well with the altered metabolites in the liver sample that the authors had published in Armstrong et al. (11). These 85 significantly altered metabolites were associated with nine human-specific pathways from the KEGG database, which can be visualized by a pathway-metabolite network (**Figure 2 (C)**). Pathway-metabolite networks can also be visualized via a heatmap (**Supplementary Dataset 1**). A dot plot can be plotted to view the association of metabolite class categories with pathways (**Supplementary Dataset 1**).

**Figure 2:**
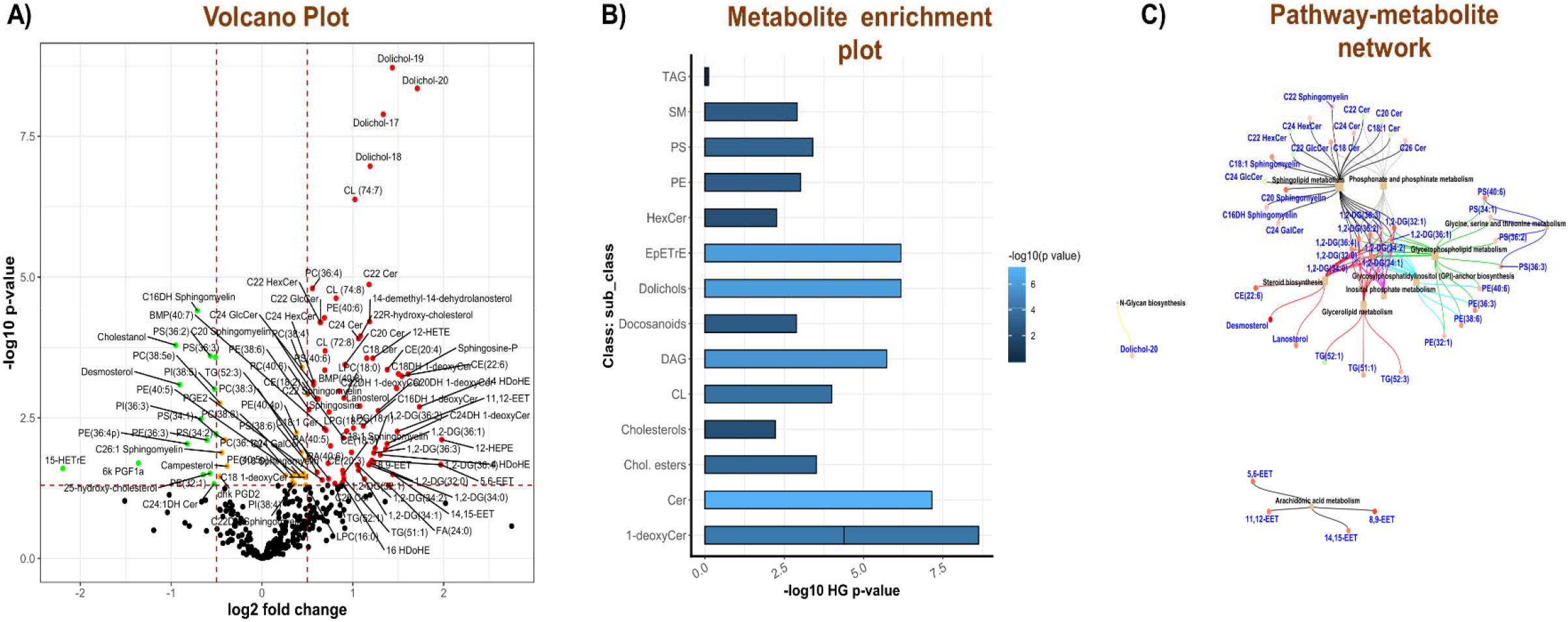
Metabolic analysis of Case Study ST000915. **A)** Volcano plot showing significantly altered metabolites. **B)** Bar plot of metabolite enrichment analysis via hypergeometric score for each metabolite class category. **C)** Pathway-metabolite network. The color of metabolites’ nodes represents the fold change of metabolites (low expression in green and high in red). All metabolites are shown in blue. The size of pathway nodes, shown as square nodes, corresponds to the number of branches representing the metabolites’ number.

In addition to this, we associated 77 reactions, 109 genes, and 46 enzymes to significantly altered metabolites found in this study. Detailed script and results file of this analysis is provided in **Supplementary file 1 and Dataset 1**.

### Case study 2 (ST001308): A growth phenotype study for *Salmonella entrica*

The second case study is an NMR metabolomics project on *Salmonella enterica* (12). This study had three experimental factors: ‘Sample_group,’ ‘Genotype,’ and ‘Supplement.’ With MetENP, users have the flexibility to compare experimental factors individually or combined (all groups: ‘Sample_group,’ ‘Genotype,’ and ‘Supplement’).

In this case study, we compared the experimental group ‘Genotype’ with factors: ridA-mutant and wild type, to get a list of significantly altered metabolites. We used the ‘half_of_min’ normalization method (*see Implementation*) to handle missing data due to metabolites with missing information in a subset of the samples. Five metabolites belonging to Amino acids and Fatty acids metabolite subclass were significantly altered with a p-value cut off 0.05. These five metabolites were observed to be associated with 11 KEGG pathways (Supplementary Dataset 2), belonging to carbohydrate metabolism, amino acid biosynthesis, and biosynthesis of secondary metabolites, respectively. Further, we examined ridA mutants having GlyA damage and Ile damage by comparing the ‘Sample_group’ experimental factor. The downstream metabolites altered due to glycine and isoleucine dependent pathways reflect growth phenotypes of ridA mutants. The analysis showed Pantothenate and CoA biosynthesis, cyanoamino acid metabolism, pyruvate metabolism, and amino acid biosynthesis, among others, to be associated with the altered metabolites. For all the results files, we refer to Supplementary Dataset 3. With the application of MetENP to this case study, we show different species can be analyzed (in this example, bacterial species) in conjunction with different experimental factors.

### Case study 3 (ST001140): Changes in the Canine Plasma Lipidome

Some metabolomics datasets are obtained using different analytical methods such as GC-MS and LC-MS. MetENP can analyze metabolites generated with either a single analytical method or multiple methods simultaneously.

To illustrate this capability, we chose this dataset obtained with three distinct analytical methods viz: i) Reversed-phase MS (for Phospholipids, Cholesterol esters and Diacylglycerols; and Sphingolipids), ii) LC-MS HILIC (For Derivatized Spingosine-1-phosphates), and iii) Normal/Stationary phase MS (For Triacylglycerols) (13). Study ID ST001140 aimed to examine the short-term and long-term exposure of glucocorticoids: Prednisolone and Tetracosactide on Canine plasma (13). The dataset was divided into four analysis types based on metabolite classes examined: a) Phospholipids, Cholesterol esters and Diacylglycerols b) Sphingolipids c) Derivatized Spingosine-1-phosphates and d) Triacylglycerols. We employed MetENP to select all four analytical types used in this study. Alternatively, users have the choice to choose a select number of these method types for comparative analysis. To calculate significantly altered metabolites, we compared the treatment group: prednisolone and tetracosactide irrespective of the exposure duration, using the normalization method ‘50percent’ (*see Implementation*). The p-value and log2 fold change cutoff were 0.05 and 0.5, respectively. Twenty-six metabolites were found to be significantly altered and were associated with 7 KEGG pathways, showing differences in canine plasma lipidome induced by different treatment methods (Supplementary Dataset 4). MetENP can also study lipidome differences due to different exposure durations of these treatment methods (*not shown*). This case study illustrated how MetENP could explore metabolite datasets generated with multiple analytical methods. For all the results files, we refer to Supplementary Dataset 4.

## CONCLUSION

MetENP workflow provides an R pipeline and a web application interface to analyze and visualize the metabolomics datasets within the Metabolomics Workbench or from a commandline entry using custom datasets. It performs metabolite enrichment analysis and aids in the functional and biological interpretation of significantly enriched metabolites for chosen experimental factors. MetENP has been integrated as a web function MetENPWeb on Metabolomics Workbench (https://www.metabolomicsworkbench.org/data/analyze.php) with an interactive GUI for data analysis and visualization. The three different case studies highlight this workflow’s flexibility. Studies of three different species types (Human, bacteria, and dog) coupled with different parameters accurately predicted the significantly altered metabolites and their associated pathways.

Specifically, MetENP R package and MetENPWeb allows users to a) map metabolites to standardized metabolite classes, b) calculate significantly altered metabolites, c) calculate enrichment score of metabolite classes, d) map to KEGG pathway of the species of choice, e) calculate the enrichment score of pathways, f) plot the pathways-metabolites network, g) get the gene, reaction information, and h) obtain enzymatic activities on the metabolites. This tool aims to make the resource more widely available for the scientific community by providing the R source code on the Metabolomics Workbench Github repository. This tool is also available as a web function MetENPWeb for the experimental biology community.

## ACKNOWLEDGEMENT

We would like to thank Dr. Srinivasan Ramachandran and Mr. Kenan Azam for helpful discussion

## FUNDING

This work was supported by the NIH Common Fund Grant, U2CDK119886, to establish the National Metabolomics Data Repository, and NIH grants OT2 OD030544 and R01 LM012595.

## SUPPLEMENTARY FILES

**Supplementary File S1:** An example run using MetENP R package

**Supplementary Dataset S1:** Results generated by running MetENPWeb on Study ID-ST000915

**Supplementary Dataset S2:** Results generated by running MetENPWeb on Study ID-ST001308 on wildtype and ridA mutants

**Supplementary Dataset S3:** Results generated by running MetENPWeb on Study ID-ST001308 on ridA mutants having GlyA damage and Ile damage.

**Supplementary Dataset S4:** Results generated by running MetENPWeb on Study ID-ST001140

## Notes

https://github.com/metabolomicsworkbench/MetENP

https://www.metabolomicsworkbench.org/data/analyze.php

### Competing Interest Statement

The authors have declared no competing interest.

