## Supplementary material for "MetENP/MetENPWeb: An R package and web application for metabolomics enrichment and pathway analysis in Metabolomics Workbench": Supplementary FIle S1.html

Case study 1: Study ST000915


### Case study 1: Study ST000915

###### Kumari Sonal Choudhary

#### 11/12/2020

#### MetENP

MetENP is a R package that enables detection of significant metabolites from metabolite information (names or names and concentration along with metadata information) and provides

Enrichment score of metabolite class, Maps to pathway of the species of choice, Calculate enrichment score of pathways, Plots the pathways and shows the metabolite increase or decrease Gets gene info, reaction info, enzyme info

```
library(MetENP)
library(jsonlite)
```

Data can be fetched via study id from Metabolomics Workbench through getmwstudies. For metabolomics data, ‘data’ is used, for metadata, ‘factors’ is used. If users want to upload their own data, please do at the bottom of the page ST000915 is the study of biomarkers of nonalcoholic fatty liver disease (NAFLD) progression.

```
metdata = getmwstudies('ST000915', 'data')
knitr::kable(head(metdata))
```

| analysis\_id | analysis\_summary | metabolite\_name | metabolite\_id | refmet\_name | NASH001 | NASH002 | NASH003 | NASH004 | NASH005 | NASH006 | NASH007 | NASH009 | NASH010 | NASH012 | NASH013 | NASH014 | NASH015 | NASH016 | NASH017 | NASH018 | NASH057 | NASH071 | NASH072 | NASH073 | NASH075 | NASH076 | NASH077 | NASH078 | NASH079 | NASH080 | NASH081 | NASH082 | NASH083 | NASH084 | NASH085 | NASH086 | NASH087 | NASH088 | NASH089 | NASH090 | NASH091 | NASH011 | NASH019 | NASH020 | NASH021 | NASH022 | NASH023 | NASH024 | NASH026 | NASH027 | NASH028 | NASH029 | NASH030 | NASH031 | NASH032 | NASH033 | NASH034 | NASH035 | NASH036 | NASH037 | NASH038 | NASH039 | NASH040 | NASH041 | NASH042 | NASH043 | NASH044 | NASH045 | NASH046 | NASH047 | NASH048 | NASH049 | NASH050 | NASH051 | NASH052 | NASH053 | NASH054 | NASH055 | NASH056 | NASH058 | NASH059 | NASH060 | NASH061 | NASH062 | NASH064 | NASH065 | NASH066 | NASH067 | NASH068 | NASH069 | NASH070 | NASH074 |
| --- | --- | --- | --- | --- | --- | --- | --- | --- | --- | --- | --- | --- | --- | --- | --- | --- | --- | --- | --- | --- | --- | --- | --- | --- | --- | --- | --- | --- | --- | --- | --- | --- | --- | --- | --- | --- | --- | --- | --- | --- | --- | --- | --- | --- | --- | --- | --- | --- | --- | --- | --- | --- | --- | --- | --- | --- | --- | --- | --- | --- | --- | --- | --- | --- | --- | --- | --- | --- | --- | --- | --- | --- | --- | --- | --- | --- | --- | --- | --- | --- | --- | --- | --- | --- | --- | --- | --- | --- | --- | --- | --- | --- |
| AN001485 | Core G Fatty acids/Eicosanoids | 10 HDoHE | ME230937 | 10-HDoHE | 0.00419 | 0.01396 | 0.00285 | 0.00705 | 0.00020 | 0.00471 | 0.00158 | 0.00463 | 0.00063 | 0.00740 | 0.00435 | 0.00763 | 0.01377 | 0.01151 | 0.00090 | 0.02084 | 0.00265 | 0.00022 | 0.00038 | 0.00189 | 0.00321 | 0.00533 | 0.00152 | 0.00150 | 0.00237 | 0.00059 | 0.00167 | 0.00073 | 0.00641 | 0.00667 | 0.00143 | 0.02844 | 0.02243 | 0.00915 | 0.00366 | 0.00208 | 0.00144 | NA | NA | NA | NA | NA | NA | NA | NA | NA | NA | NA | NA | NA | NA | NA | NA | NA | NA | NA | NA | NA | NA | NA | NA | NA | NA | NA | NA | NA | NA | NA | NA | NA | NA | NA | NA | NA | NA | NA | NA | NA | NA | NA | NA | NA | NA | NA | NA | NA | NA | NA |
| AN001485 | Core G Fatty acids/Eicosanoids | 11,12-diHETrE | ME230961 | 11,12-DiHETrE | 0.01346 | 1.84739 | 0.63897 | 1.87466 | 0.21119 | 2.23019 | 0.29970 | 1.17290 | 0.25164 | 0.90318 | 0.90953 | 0.70945 | 0.43454 | 0.81423 | 0.73651 | 0.16733 | 0.27242 | 0.02657 | 0.01304 | 0.02329 | 0.05239 | 0.02907 | 0.06744 | 0.05461 | 0.05734 | 0.02636 | 0.02764 | 0.06045 | 0.03386 | 0.01953 | 0.02720 | 0.02104 | 0.06057 | 0.03444 | 0.01259 | 0.02891 | 0.05983 | 1.36108 | 0.44150 | 0.02390 | 0.14849 | 0.06360 | 0.07685 | 0.03539 | 0.20321 | 0.09351 | 0.14849 | 0.10522 | 0.06791 | 0.07228 | 0.15854 | 0.04268 | 0.03593 | 0.07740 | 0.10180 | 0.02219 | 0.03319 | 0.03399 | 0.02789 | 0.06747 | 0.13379 | 0.11856 | 0.26173 | 0.15304 | 0.18052 | 0.08712 | 0.19573 | 0.16578 | 0.03662 | 0.09379 | 0.49311 | 0.64944 | 0.17857 | 0.10998 | 0.13538 | 0.03176 | 0.06515 | 0.17947 | 0.12842 | 0.16833 | 0.12114 | 0.32882 | 0.10923 | 0.21540 | 0.24789 | 0.25932 | 0.03551 | 0.05935 |
| AN001485 | Core G Fatty acids/Eicosanoids | 11,12-EET | ME230954 | 11,12-EpETrE | 0.20666 | 0.12915 | 0.18170 | 0.11368 | 0.06081 | 0.34385 | 0.04465 | 0.05223 | 0.04092 | 0.15712 | 0.24605 | 0.22513 | 0.33102 | 0.12221 | 0.15930 | 0.07123 | 0.12722 | 0.14079 | 0.03025 | 0.10854 | 0.26357 | 0.09159 | 0.35800 | 0.22358 | 0.11807 | 0.03547 | 0.06269 | 0.97974 | 0.66245 | 0.19845 | 0.30318 | 0.37735 | 0.67719 | 0.64060 | 0.39347 | 0.22317 | 0.38579 | 0.06281 | 0.20807 | 0.03775 | 0.05421 | 0.02036 | 0.02334 | 0.02255 | 0.03843 | 0.01395 | 0.10388 | 0.03130 | 0.02376 | 0.04410 | 0.06975 | 0.00947 | 0.02032 | 0.02166 | 0.04561 | 0.01040 | 0.01710 | 0.01379 | 0.01428 | 0.03581 | 0.16917 | 0.04188 | 0.06256 | 0.03610 | 0.03925 | 0.01802 | 0.01669 | 0.02359 | 0.02850 | 0.02779 | 0.08402 | 0.03029 | 0.04080 | 0.02240 | 0.01856 | 0.00807 | 0.04488 | 0.22406 | 0.06934 | 0.06330 | 0.04931 | 0.10011 | 0.10355 | 0.16318 | 0.10468 | 0.05115 | 0.08985 | 0.10916 |
| AN001485 | Core G Fatty acids/Eicosanoids | 11b PGE2 | ME230914 | 11beta-PGE2 | 0.00048 | 0.00128 | NA | NA | NA | NA | NA | NA | NA | NA | NA | NA | NA | 0.00533 | NA | 0.00264 | NA | 0.00262 | 0.00693 | 0.00187 | 0.00078 | 0.00072 | NA | NA | 0.00277 | 0.00020 | NA | 0.00098 | 0.00206 | 0.00076 | 0.00213 | 0.00093 | 0.02040 | 0.02086 | 0.00070 | 0.00136 | 0.00969 | NA | 0.00214 | NA | NA | 0.00203 | 0.00174 | 0.00026 | NA | NA | 0.00138 | NA | 0.00016 | 0.00079 | 0.00176 | NA | 0.00205 | 0.00156 | 0.00068 | 0.00298 | 0.00108 | 0.00169 | 0.00104 | 0.00052 | NA | 0.00184 | NA | NA | 0.00175 | 0.00743 | NA | NA | NA | 0.00096 | 0.00528 | NA | NA | 0.00097 | 0.00262 | 0.00178 | NA | NA | 0.00165 | 0.00327 | 0.00153 | NA | NA | NA | NA | 0.00445 | NA | NA |
| AN001485 | Core G Fatty acids/Eicosanoids | 11 HDoHE | ME230941 | 11-HDoHE | 0.00617 | 0.03174 | 0.00958 | 0.00593 | 0.00271 | 0.05614 | 0.00784 | 0.03434 | 0.00303 | 0.02391 | 0.00861 | 0.01333 | 0.03302 | 0.02120 | 0.01284 | 0.02260 | 0.01865 | 0.00433 | 0.00596 | 0.00903 | 0.00737 | 0.01596 | 0.00938 | 0.01685 | 0.00716 | 0.00208 | 0.02655 | 0.01842 | 0.01902 | 0.01049 | 0.00151 | 0.02371 | 0.03326 | 0.00267 | 0.00232 | NA | 0.00584 | 0.01793 | 0.00857 | NA | 0.00473 | NA | 0.00351 | 0.00071 | NA | NA | NA | NA | 0.00398 | 0.00166 | 0.00470 | NA | 0.00488 | 0.00624 | 0.00884 | NA | NA | NA | NA | NA | 0.00600 | 0.02054 | 0.01020 | 0.02115 | 0.01072 | 0.00653 | 0.01549 | 0.01189 | 0.00265 | 0.00751 | 0.02317 | 0.15263 | 0.01830 | 0.00343 | 0.00885 | NA | 0.00737 | 0.03027 | 0.03543 | 0.01280 | 0.00369 | 0.01844 | 0.00649 | 0.03449 | 0.01077 | 0.01869 | 0.00414 | 0.01129 |
| AN001485 | Core G Fatty acids/Eicosanoids | 11-HEPE | ME230917 | 11-HEPE | 0.00642 | NA | 0.01625 | 0.02049 | 0.00115 | 0.01440 | 0.00628 | 0.03413 | NA | 0.01394 | 0.00608 | 0.02614 | 0.00429 | 0.03508 | 0.01276 | 0.00453 | 0.00936 | 0.00567 | 0.00041 | NA | 0.00119 | 0.00098 | 0.00035 | 0.00111 | 0.01423 | 0.00003 | 0.00840 | 0.02481 | 0.00969 | 0.00109 | NA | 0.02083 | 0.00539 | 0.00666 | 0.00745 | 0.01074 | 0.00425 | 0.04633 | 0.00361 | 0.00432 | NA | NA | NA | NA | 0.00018 | 0.00487 | 0.00026 | NA | 0.00325 | NA | 0.00100 | 0.00198 | NA | 0.01562 | NA | NA | NA | NA | NA | 0.00378 | 0.00513 | 0.00391 | 0.01931 | 0.02274 | 0.00528 | 0.00714 | 0.02485 | 0.02938 | 0.05539 | 0.02308 | 0.08151 | 0.00722 | 0.03255 | 0.11625 | 0.01155 | 0.01965 | 0.06180 | 0.01783 | 0.14650 | 0.03245 | 0.01528 | 0.01148 | 0.03754 | 0.01260 | 0.03738 | 0.00957 | 0.00057 | 0.01524 |

Associate metabolomics data to the refmet class

```
refmet_class= convert_refmet(metdata)
```

```
metadata = getmwstudies('ST000915', 'factors')
knitr::kable(head(metadata))
```

| study\_id | local\_sample\_id | subject\_type | factors | Diagnosis |
| --- | --- | --- | --- | --- |
| ST000915 | NASH005 | Human clinical study | Diagnosis:Cirrhosis | Cirrhosis |
| ST000915 | NASH007 | Human clinical study | Diagnosis:Cirrhosis | Cirrhosis |
| ST000915 | NASH009 | Human clinical study | Diagnosis:Cirrhosis | Cirrhosis |
| ST000915 | NASH013 | Human clinical study | Diagnosis:Cirrhosis | Cirrhosis |
| ST000915 | NASH016 | Human clinical study | Diagnosis:Cirrhosis | Cirrhosis |
| ST000915 | NASH022 | Human clinical study | Diagnosis:Cirrhosis | Cirrhosis |

Find the factors you would want to compare. Multiple factors/experimental groups (independent variables) are formatted in multiple columns but you can get information on all the experimental groups by “factors” column. For t-test use the independent variables in the same column. For comparing multipe independent variables use anova by anova\_ana function.

For ex: in this study, we want to compare Diagnosis experimental methods, so we will compare Cirrhosis and Normal samples

```
unique(metadata$factors)
```

```
## [[1]]
## [1] "Diagnosis:Cirrhosis"
## 
## [[2]]
## [1] "Diagnosis:NASH"
## 
## [[3]]
## [1] "Diagnosis:Normal"
## 
## [[4]]
## [1] "Diagnosis:Steatosis"
```

Find different type of analysis mode. This is important, because some studies may have different analysis types, and different analysis types detect different metabolites. In this study, there is only one analysis type.

```
### Find the analysis mode
unique(metdata$analysis_summary)
```

```
## [[1]]
## [1] "Core G Fatty acids/Eicosanoids"
## 
## [[2]]
## [1] "Core J Sterols"
## 
## [[3]]
## [1] "Core K Prenols/Cardiolipins"
## 
## [[4]]
## [1] "Core E Neutral Lipids"
## 
## [[5]]
## [1] "Core I Sphingolipids"
## 
## [[6]]
## [1] "Core H Phospholipids"
```

##### Find significant metabolites, run significance of all the analysis summary together. The analysis summary/modes you got in the previous section

Significant metabolites can be found by using significant\_met function. The parameters are:

metabolomics\_data: metabolomics data associated to refmet class

met\_col: column with metabolite names

analysis\_type: type of analysis ex-GCMS, HILIAC positive ion mode.

metadata: Metadata

factor1: first independent variable

factor2: second independent variable

factor\_col: column name of the independent variables

sample\_col: the column name having samples

p\_adjust: Method for p value adjustment, i.e. “fdr”

normalization: method for normalization a) “half\_of\_min”: where the NAs are replaced by half of min values in the data b) “remove\_NAs”: where Cols with NAs values are removed and c) “50percent”: where cols with more than 50% NAs values are removed

half\_of\_min is ideal when you wish to see which metabolites were present in either group. Very high fold change would mean it was present in either group.

In this example, we show significant metabolite filtered according to pvalue threshold of 0.05 and log2 fold change of 0.5, p adjust method of ‘fdr’ and we use the normalization method “50percent”

```
stats_metabolites = significant_met(metabolomics_data=refmet_class, met_col="metabolite_name",analysis_type=c("Core G Fatty acids/Eicosanoids","Core J Sterols","Core K Prenols/Cardiolipins","Core E Neutral Lipids","Core I Sphingolipids","Core H Phospholipids"), metadata=metadata, factor1='Cirrhosis', factor2='Normal', factor_col='Diagnosis',sample_col='local_sample_id', p_adjust='fdr',normalization="50percent")

sig_metabolites = stats_metabolites[which(stats_metabolites[,"pval"] <= 0.05&abs(stats_metabolites[,"log2Fold_change"])>0.5),]
knitr::kable(head(sig_metabolites))
```

|  | Metabolite | Cirrhosis\_mean | Normal\_mean | Fold\_change | log2Fold\_change | t\_value | pval | padj | metabolite\_id | refmet\_name | super\_class | main\_class | sub\_class | formula |
| --- | --- | --- | --- | --- | --- | --- | --- | --- | --- | --- | --- | --- | --- | --- |
| 5 | 11,12-EET | 0.0609060 | 0.1709519 | 2.8068154 | 1.488934 | -2.946309 | 0.0055361 | 0.1384028 | ME230954 | 11,12-EpETrE | Fatty Acyls | Eicosanoids | EpETrE | C20H32O3 |
| 6 | 12-HEPE | 0.00945950 | 0.03728742 | 3.9417961 | 1.978853 | -2.846260 | 0.0077250 | 0.1448446 | ME230939 | 12-HEPE | Fatty Acyls | Eicosanoids | HEPE | C20H30O3 |
| 7 | 12-HETE | 0.109529 | 0.255101 | 2.3290727 | 1.219756 | -3.994515 | 0.0002766 | 0.0207450 | ME230938 | 12-HETE | Fatty Acyls | Eicosanoids | HETE | C20H32O3 |
| 12 | 14 HDoHE | 0.00825450 | 0.02744806 | 3.3252238 | 1.733451 | -3.316098 | 0.0020102 | 0.0753841 | ME230940 | 14-HDoHE | Fatty Acyls | Docosanoids | Docosanoids | C22H32O3 |
| 14 | 14,15-EET | 0.1171725 | 0.3177545 | 2.7118522 | 1.439278 | -2.236125 | 0.0318876 | 0.2385268 | ME230955 | 14,15-EpETrE | Fatty Acyls | Eicosanoids | EpETrE | C20H32O3 |
| 17 | 15-HETrE | 0.04673850 | 0.01022645 | 0.2188014 | -2.192306 | 2.425914 | 0.0250871 | 0.2090595 | ME230935 | 15-HeTrE | Fatty Acyls | Eicosanoids | ETrE | C20H34O3 |

##### Significant Metabolites can be visualized via Volcano Plot

We will use the same thresholds as used for determining significant metabolites

```
plot_volcano(stats_metabolites, thres_pval= 0.05,thres_log2foldchange = 0.5, TRUE)
```

##### Map metabolite class of the significant metabolites utilzing refmet classification in Metabolomics Workbench

This function maps metabolite to metabolite class

In this example, we will go forward with significant metabolite obtained by t-test

```
sig_metabolites_kegg_id= map_keggid(sig_metabolites)
knitr::kable(head(sig_metabolites_kegg_id))
```

| refmet\_name | Exact mass | KEGG ID | Metabolite | Cirrhosis\_mean | Normal\_mean | Fold\_change | log2Fold\_change | t\_value | pval | padj | metabolite\_id | super\_class | main\_class | sub\_class | formula |
| --- | --- | --- | --- | --- | --- | --- | --- | --- | --- | --- | --- | --- | --- | --- | --- |
| 1-DeoxyCer 18:0;O2/16:0 | 523.5328 |  | C16DH 1-deoxyCer | 0.05623685 | 0.12178748 | 2.165617 | 1.114778 | -3.028001 | 0.0044114 | 0.0251708 | ME231305 | Sphingolipids | Ceramides | 1-deoxyCer | C34H69NO2 |
| 1-DeoxyCer 18:0;O2/18:0 | 551.5641 |  | C18DH 1-deoxyCer | 0.0906803 | 0.2566956 | 2.830776 | 1.501197 | -3.826879 | 0.0005220 | 0.0056713 | ME231306 | Sphingolipids | Ceramides | 1-deoxyCer | C36H73NO2 |
| 1-DeoxyCer 18:0;O2/20:0 | 579.5954 |  | C20DH 1-deoxyCer | 0.0723488 | 0.1756429 | 2.427724 | 1.279604 | -3.268230 | 0.0023663 | 0.0150818 | ME231307 | Sphingolipids | Ceramides | 1-deoxyCer | C38H77NO2 |
| 1-DeoxyCer 18:0;O2/22:0 | 607.6267 |  | C22DH 1-deoxyCer | 0.1513816 | 0.4227716 | 2.792754 | 1.481689 | -3.604410 | 0.0009458 | 0.0076456 | ME231308 | Sphingolipids | Ceramides | 1-deoxyCer | C40H81NO2 |
| 1-DeoxyCer 18:0;O2/24:0 | 635.6580 |  | C24DH 1-deoxyCer | 0.1177141 | 0.3056195 | 2.596286 | 1.376450 | -2.746084 | 0.0091800 | 0.0468664 | ME231310 | Sphingolipids | Ceramides | 1-deoxyCer | C42H85NO2 |
| 1,2-DG 32:0 | 568.5067 | C00641 | 1,2-DG(32:0) | 210.0625 | 479.9210 | 2.284658 | 1.191978 | -2.410346 | 0.0200254 | 0.1066535 | ME231196 | Glycerolipids | Diradylglycerols | DAG | C35H68O5 |

###### Check all your significant metabolites have not been assigned metabolite class

```
setdiff(sig_metabolites$refmet_name, sig_metabolites_kegg_id$refmet_name)
```

```
## character(0)
```

##### Count metabolites in each of the metabolite class and plotting

You may choose from sub\_class, main\_class and super\_class

```
count_changes = metcountplot(df_metclass=sig_metabolites_kegg_id, metclass='sub_class', plotting=TRUE, thres_logfC = 0.5)
```

To see the Plot

```
count_changes$plotimg
```

##### Enrichment class score

Calculate the enrichment score of each metabolite class. Enrichment score is calculated through hypergeometric method. One can specify the no. of significant metabolites in a class while calculating the enrichment score. We advice to use the number of mtabolites in each class as 3 or more. But if someone just wants to know the enrichment score and rest of the information of all the metabolites, then they can choose the number as 1.

```
metenrichment = metclassenrichment(df_metclass=sig_metabolites_kegg_id,refmet_class, metclass="sub_class",enrich_stats="HG",no=3)
knitr::kable(head(metenrichment))
```

| refmet\_name | Exact mass | KEGG ID | Metabolite | Cirrhosis\_mean | Normal\_mean | Fold\_change | log2Fold\_change | t\_value | pval | padj | metabolite\_id | super\_class | main\_class | sub\_class | formula | HG p-value |
| --- | --- | --- | --- | --- | --- | --- | --- | --- | --- | --- | --- | --- | --- | --- | --- | --- |
| 1-DeoxyCer 18:0;O2/16:0 | 523.5328 |  | C16DH 1-deoxyCer | 0.05623685 | 0.12178748 | 2.165617 | 1.114778 | -3.028001 | 0.0044114 | 0.0251708 | ME231305 | Sphingolipids | Ceramides | 1-deoxyCer | C34H69NO2 | 4.12e-05 |
| 1-DeoxyCer 18:0;O2/18:0 | 551.5641 |  | C18DH 1-deoxyCer | 0.0906803 | 0.2566956 | 2.830776 | 1.501197 | -3.826879 | 0.0005220 | 0.0056713 | ME231306 | Sphingolipids | Ceramides | 1-deoxyCer | C36H73NO2 | 5.74e-05 |
| 1-DeoxyCer 18:0;O2/20:0 | 579.5954 |  | C20DH 1-deoxyCer | 0.0723488 | 0.1756429 | 2.427724 | 1.279604 | -3.268230 | 0.0023663 | 0.0150818 | ME231307 | Sphingolipids | Ceramides | 1-deoxyCer | C38H77NO2 | 5.74e-05 |
| 1-DeoxyCer 18:0;O2/22:0 | 607.6267 |  | C22DH 1-deoxyCer | 0.1513816 | 0.4227716 | 2.792754 | 1.481689 | -3.604410 | 0.0009458 | 0.0076456 | ME231308 | Sphingolipids | Ceramides | 1-deoxyCer | C40H81NO2 | 5.74e-05 |
| 1-DeoxyCer 18:0;O2/24:0 | 635.6580 |  | C24DH 1-deoxyCer | 0.1177141 | 0.3056195 | 2.596286 | 1.376450 | -2.746084 | 0.0091800 | 0.0468664 | ME231310 | Sphingolipids | Ceramides | 1-deoxyCer | C42H85NO2 | 5.74e-05 |
| 1,2-DG 32:0 | 568.5067 | C00641 | 1,2-DG(32:0) | 210.0625 | 479.9210 | 2.284658 | 1.191978 | -2.410346 | 0.0200254 | 0.1066535 | ME231196 | Glycerolipids | Diradylglycerols | DAG | C35H68O5 | 1.90e-06 |

###### Plot the enrichment score via function plot\_met\_enrichment

```
plot_met_enrichment(metenrichment, "sub_class","HG", no=3)
```

##### Check the pathways with reactions of all the significant metabolites in Species specific manner.

First parameter is the metenrichment dataframe, while second parameter is the species code from KEGG

Here the subject species is Homo sapiens, and the KEGG species annotation for KEGG is ‘hsa’

```
met_path = met_pathways(df_metenrichment = metenrichment, 'hsa')
knitr::kable(head(met_path))
```

| rxn | refmet\_name | Exact mass | KEGG ID | Metabolite | Cirrhosis\_mean | Normal\_mean | Fold\_change | log2Fold\_change | t\_value | pval | padj | metabolite\_id | super\_class | main\_class | sub\_class | formula | HG p-value | Rxn\_name | PATHWAY | pathway\_id | sps\_path\_id |
| --- | --- | --- | --- | --- | --- | --- | --- | --- | --- | --- | --- | --- | --- | --- | --- | --- | --- | --- | --- | --- | --- |
| R01003 | Dolichol-20 | 1381.2782 | C00381 | Dolichol-20 | 18.54100 | 60.64903 | 3.271076 | 1.709766 | -7.141005 | 0.0000000 | 0.0000000 | ME231044 | Prenol Lipids | Polyprenols | Dolichols | C100H164O | 7.0e-07 | dolichyl-phosphate phosphohydrolase | N-Glycan biosynthesis | rn00510 | hsa00510 |
| R01003 | Dolichol-20 | 1381.2782 | C00381 | Dolichol-20 | 18.54100 | 60.64903 | 3.271076 | 1.709766 | -7.141005 | 0.0000000 | 0.0000000 | ME231044 | Prenol Lipids | Polyprenols | Dolichols | C100H164O | 7.0e-07 | dolichyl-phosphate phosphohydrolase | Metabolic pathways | rn01100 | hsa01100 |
| R01018 | Dolichol-20 | 1381.2782 | C00381 | Dolichol-20 | 18.54100 | 60.64903 | 3.271076 | 1.709766 | -7.141005 | 0.0000000 | 0.0000000 | ME231044 | Prenol Lipids | Polyprenols | Dolichols | C100H164O | 7.0e-07 | CTP:dolichol O-phosphotransferase | N-Glycan biosynthesis | rn00510 | hsa00510 |
| R01018 | Dolichol-20 | 1381.2782 | C00381 | Dolichol-20 | 18.54100 | 60.64903 | 3.271076 | 1.709766 | -7.141005 | 0.0000000 | 0.0000000 | ME231044 | Prenol Lipids | Polyprenols | Dolichols | C100H164O | 7.0e-07 | CTP:dolichol O-phosphotransferase | Metabolic pathways | rn01100 | hsa01100 |
| R01312 | 1,2-DG 34:0 | 596.5380 | C00641 | 1,2-DG(34:0) | 182.8600 | 413.3597 | 2.260525 | 1.176658 | -2.377108 | 0.0217006 | 0.1066535 | ME231203 | Glycerolipids | Diradylglycerols | DAG | C37H72O5 | 1.9e-06 | Phosphatidylcholine cholinephosphohydrolase | Glycerophospholipid metabolism | rn00564 | hsa00564 |
| R01312 | 1,2-DG 36:4 | 616.5067 | C00641 | 1,2-DG(36:4) | 165.5750 | 383.1855 | 2.314271 | 1.210558 | -2.453312 | 0.0180560 | 0.1066535 | ME231210 | Glycerolipids | Diradylglycerols | DAG | C39H68O5 | 1.9e-06 | Phosphatidylcholine cholinephosphohydrolase | Glycerophospholipid metabolism | rn00564 | hsa00564 |

Find metabolites for which no pathways were registered in Kegg and/or no kegg id was found

```
setdiff(metenrichment$Metabolite,unique(met_path$Metabolite))
```

```
##  [1] "C16DH 1-deoxyCer" "C18DH 1-deoxyCer" "C20DH 1-deoxyCer" "C22DH 1-deoxyCer"
##  [5] "C24DH 1-deoxyCer" "14 HDoHE"         "16 HDoHE"         "4 HDoHE"         
##  [9] "CE(18:2)"         "CE(18:3)"         "CE(20:3)"         "CE(20:4)"        
## [13] "CL (72:8)"        "CL (74:7)"        "CL (74:8)"        "Cholestanol"     
## [17] "Dolichol-17"      "Dolichol-18"      "Dolichol-19"
```

##### Get pathway enrichment sore.

Once we have the pathway information, we can calculate enrichment score of pathways. Again, here i have used hypergeometric score.

There are two ways to calculate in MetENP. If users want to calculate based on total no of metabolites detected in the study N= the total number of metabolites in a study, which can be retrieved from refmet\_class dataframe.

Instead, if the users want to calculate HG score such that they can compare two studies, the N is the total no. of cmpds linked to kegg pathway (this is the step which might take long), so I advice to run the script comp\_linkedto\_pathways() just the first time or after 6 months or so, if desired to run the pipeline again. save the result from comp\_linkedto\_pathways() and load it. Loading from saved file would save time for another analysis with another study.

L = all significant metabolites detected in a study M= all significant metabolites detected in a pathway K = all metabolites detected in a pathway

phyper(M,L, N-L, K)

This function also utilizes korg dataset from pathview package.

In this example, the background set N is the total number of metabolites in a study

```
load('C:/Users/bioso/Documents/MetENP/data/ls_path.RData')
load('C:/Users/bioso/Documents/MetENP/data/korg.RData')
kegg_es = path_enrichmentscore(met_path,sig_metabolite_kegg_id=sig_metabolite_kegg_id,ls_path=ls_path,refmet_class=refmet_class,sps='hsa',padj='fdr', kegg_comp_path=FALSE)
knitr::kable(head(kegg_es))
```

|  | Pathway name | No.of mets in study | Total\_no.\_of\_comps\_in\_pathway | pathway\_HG p-value | Padjust |
| --- | --- | --- | --- | --- | --- |
| 2 | Arachidonic acid metabolism | 4 | 75 | 0.9991730 | 0.9991730 |
| 14 | Glycerolipid metabolism | 12 | 38 | 0.0084694 | 0.0211735 |
| 13 | Glycerophospholipid metabolism | 17 | 52 | 0.0009973 | 0.0033243 |
| 17 | Glycine, serine and threonine metabolism | 4 | 50 | 0.9705819 | 0.9991730 |
| 110 | Glycosylphosphatidylinositol (GPI)-anchor biosynthesis | 13 | 16 | 0.0000000 | 0.0000000 |
| 18 | Inositol phosphate metabolism | 9 | 47 | 0.3070207 | 0.4386011 |

##### Plot pathway network

The pathway network is such that it shows metabolites that are connected to different pathways and same metabolite in different pathway. Color of nodes of metabolites are according to the fold change of metabolites (low expression in green and high in red) and size of pathway nodes (square nodes) are according to the number of branches (meaning no of metabolites). All metabolite are written in blue

```
plot_pathway_networks (met_path,kegg_es, TRUE)
```

##### Heatmap

```
plot_heatmap(met_path, shorten_name=TRUE,refmet_name=FALSE, xaxis=8, yaxis=6)
```

##### Dotplot

```
dotplot_met_class_path (met_path, kegg_es,"sub_class",xaxis=10,yaxis=10)
```

##### Get the gene and enzyme info

Here we get the information of genes involved in enriched pathways for specified organism

```
met_gene_info = enzyme_gene_info (metenrichment, "hsa","sub_class")
knitr::kable(head(met_gene_info))
```

| orthology\_id | ORTHOLOGY | gene\_id | gene\_name | DEFINITION | ORGANISM | PATHWAY | DBLINKS | MOTIF | rxn | Metabolite | KEGG ID | sub\_class | Rxn\_name | RCLASS | EQUATION | EQUATION\_more | ENZYME |
| --- | --- | --- | --- | --- | --- | --- | --- | --- | --- | --- | --- | --- | --- | --- | --- | --- | --- |
| K00551 | phosphatidylethanolamine/phosphatidyl-N-methylethanolamine N-methyltransferase [EC:2.1.1.17 2.1.1.71] | 10400 | PEMT, PEAMT, PEMPT, PEMT2, PLMT, PNMT | (RefSeq) phosphatidylethanolamine N-methyltransferase | Homo sapiens (human) | c(hsa00564 = “Glycerophospholipid metabolism”, hsa01100 = “Metabolic pathways”) | c(“NCBI-GeneID: 10400”, “NCBI-ProteinID: NP\_009100”, “OMIM: 602391”, “HGNC: 8830”, “Ensembl: ENSG00000133027”, “Vega: OTTHUMG00000059290”, “Pharos: Q9UBM1(Tbio)”, “UniProt: Q9UBM1”) | Pfam: PEMT Herpes\_UL74 | R02056 | PE(32:1) | C00350 | PE | S-adenosyl-L-methionine:phosphatidylethanolamine N-methyltransferase | c(“RC00003 C00019\_C00021”, “RC00060 C00350\_C01241”) | C00019 + C00350 <=> C00021 + C01241 | S-Adenosyl-L-methionine + Phosphatidylethanolamine <=> S-Adenosyl-L-homocysteine + Phosphatidyl-N-methylethanolamine | 2.1.1.17 |
| K00551 | phosphatidylethanolamine/phosphatidyl-N-methylethanolamine N-methyltransferase [EC:2.1.1.17 2.1.1.71] | 10400 | PEMT, PEAMT, PEMPT, PEMT2, PLMT, PNMT | (RefSeq) phosphatidylethanolamine N-methyltransferase | Homo sapiens (human) | c(hsa00564 = “Glycerophospholipid metabolism”, hsa01100 = “Metabolic pathways”) | c(“NCBI-GeneID: 10400”, “NCBI-ProteinID: NP\_009100”, “OMIM: 602391”, “HGNC: 8830”, “Ensembl: ENSG00000133027”, “Vega: OTTHUMG00000059290”, “Pharos: Q9UBM1(Tbio)”, “UniProt: Q9UBM1”) | Pfam: PEMT Herpes\_UL74 | R02056 | PE(38:6) | C00350 | PE | S-adenosyl-L-methionine:phosphatidylethanolamine N-methyltransferase | c(“RC00003 C00019\_C00021”, “RC00060 C00350\_C01241”) | C00019 + C00350 <=> C00021 + C01241 | S-Adenosyl-L-methionine + Phosphatidylethanolamine <=> S-Adenosyl-L-homocysteine + Phosphatidyl-N-methylethanolamine | 2.1.1.17 |
| K00551 | phosphatidylethanolamine/phosphatidyl-N-methylethanolamine N-methyltransferase [EC:2.1.1.17 2.1.1.71] | 10400 | PEMT, PEAMT, PEMPT, PEMT2, PLMT, PNMT | (RefSeq) phosphatidylethanolamine N-methyltransferase | Homo sapiens (human) | c(hsa00564 = “Glycerophospholipid metabolism”, hsa01100 = “Metabolic pathways”) | c(“NCBI-GeneID: 10400”, “NCBI-ProteinID: NP\_009100”, “OMIM: 602391”, “HGNC: 8830”, “Ensembl: ENSG00000133027”, “Vega: OTTHUMG00000059290”, “Pharos: Q9UBM1(Tbio)”, “UniProt: Q9UBM1”) | Pfam: PEMT Herpes\_UL74 | R02056 | PE(40:6) | C00350 | PE | S-adenosyl-L-methionine:phosphatidylethanolamine N-methyltransferase | c(“RC00003 C00019\_C00021”, “RC00060 C00350\_C01241”) | C00019 + C00350 <=> C00021 + C01241 | S-Adenosyl-L-methionine + Phosphatidylethanolamine <=> S-Adenosyl-L-homocysteine + Phosphatidyl-N-methylethanolamine | 2.1.1.17 |
| K00551 | phosphatidylethanolamine/phosphatidyl-N-methylethanolamine N-methyltransferase [EC:2.1.1.17 2.1.1.71] | 10400 | PEMT, PEAMT, PEMPT, PEMT2, PLMT, PNMT | (RefSeq) phosphatidylethanolamine N-methyltransferase | Homo sapiens (human) | c(hsa00564 = “Glycerophospholipid metabolism”, hsa01100 = “Metabolic pathways”) | c(“NCBI-GeneID: 10400”, “NCBI-ProteinID: NP\_009100”, “OMIM: 602391”, “HGNC: 8830”, “Ensembl: ENSG00000133027”, “Vega: OTTHUMG00000059290”, “Pharos: Q9UBM1(Tbio)”, “UniProt: Q9UBM1”) | Pfam: PEMT Herpes\_UL74 | R02056 | PE(36:3) | C00350 | PE | S-adenosyl-L-methionine:phosphatidylethanolamine N-methyltransferase | c(“RC00003 C00019\_C00021”, “RC00060 C00350\_C01241”) | C00019 + C00350 <=> C00021 + C01241 | S-Adenosyl-L-methionine + Phosphatidylethanolamine <=> S-Adenosyl-L-homocysteine + Phosphatidyl-N-methylethanolamine | 2.1.1.17 |
| K00637 | sterol O-acyltransferase [EC:2.3.1.26] | 6646 | SOAT1, ACACT, ACAT, ACAT-1, ACAT1, SOAT, STAT | (RefSeq) sterol O-acyltransferase 1 | Homo sapiens (human) | c(hsa00100 = “Steroid biosynthesis”, hsa04979 = “Cholesterol metabolism”) | c(“NCBI-GeneID: 6646”, “NCBI-ProteinID: NP\_003092”, “OMIM: 102642”, “HGNC: 11177”, “Ensembl: ENSG00000057252”, “Vega: OTTHUMG00000035253”, “Pharos: P35610(Tchem)”, “UniProt: P35610”) | Pfam: MBOAT COX6C | R01461 | CE(22:6) | C02530 | Chol. esters | Acyl-CoA:cholesterol O-acyltransferase | c(“RC00004 C00010\_C00040”, “RC00055 C00187\_C02530”) | C00040 + C00187 <=> C00010 + C02530 | Acyl-CoA + Cholesterol <=> CoA + Cholesterol ester | 2.3.1.26 |
| K00637 | sterol O-acyltransferase [EC:2.3.1.26] | 8435 | SOAT2, ACACT2, ACAT2, ARGP2 | (RefSeq) sterol O-acyltransferase 2 | Homo sapiens (human) | c(hsa00100 = “Steroid biosynthesis”, hsa04979 = “Cholesterol metabolism”) | c(“NCBI-GeneID: 8435”, “NCBI-ProteinID: NP\_003569”, “OMIM: 601311”, “HGNC: 11178”, “Ensembl: ENSG00000167780”, “Vega: OTTHUMG00000169774”, “Pharos: O75908(Tchem)”, “UniProt: O75908”) | Pfam: MBOAT MBOAT\_2 | R01461 | CE(22:6) | C02530 | Chol. esters | Acyl-CoA:cholesterol O-acyltransferase | c(“RC00004 C00010\_C00040”, “RC00055 C00187\_C02530”) | C00040 + C00187 <=> C00010 + C02530 | Acyl-CoA + Cholesterol <=> CoA + Cholesterol ester | 2.3.1.26 |

###### Get the information if metabolite is a reactant or substrate

```
rclass_info = react_substrate(met_gene_info)
knitr::kable(head(rclass_info))
```

| orthology\_id | ORTHOLOGY | gene\_id | gene\_name | DEFINITION | ORGANISM | PATHWAY | DBLINKS | MOTIF | rxn | Metabolite | KEGG ID | sub\_class | Rxn\_name | RCLASS | EQUATION | EQUATION\_more | ENZYME | reactant\_product |
| --- | --- | --- | --- | --- | --- | --- | --- | --- | --- | --- | --- | --- | --- | --- | --- | --- | --- | --- |
| K00551 | phosphatidylethanolamine/phosphatidyl-N-methylethanolamine N-methyltransferase [EC:2.1.1.17 2.1.1.71] | 10400 | PEMT, PEAMT, PEMPT, PEMT2, PLMT, PNMT | (RefSeq) phosphatidylethanolamine N-methyltransferase | Homo sapiens (human) | c(hsa00564 = “Glycerophospholipid metabolism”, hsa01100 = “Metabolic pathways”) | c(“NCBI-GeneID: 10400”, “NCBI-ProteinID: NP\_009100”, “OMIM: 602391”, “HGNC: 8830”, “Ensembl: ENSG00000133027”, “Vega: OTTHUMG00000059290”, “Pharos: Q9UBM1(Tbio)”, “UniProt: Q9UBM1”) | Pfam: PEMT Herpes\_UL74 | R02056 | PE(32:1) | C00350 | PE | S-adenosyl-L-methionine:phosphatidylethanolamine N-methyltransferase | c(“RC00003 C00019\_C00021”, “RC00060 C00350\_C01241”) | C00019 + C00350 <=> C00021 + C01241 | S-Adenosyl-L-methionine + Phosphatidylethanolamine <=> S-Adenosyl-L-homocysteine + Phosphatidyl-N-methylethanolamine | 2.1.1.17 | Substrate |
| K00551 | phosphatidylethanolamine/phosphatidyl-N-methylethanolamine N-methyltransferase [EC:2.1.1.17 2.1.1.71] | 10400 | PEMT, PEAMT, PEMPT, PEMT2, PLMT, PNMT | (RefSeq) phosphatidylethanolamine N-methyltransferase | Homo sapiens (human) | c(hsa00564 = “Glycerophospholipid metabolism”, hsa01100 = “Metabolic pathways”) | c(“NCBI-GeneID: 10400”, “NCBI-ProteinID: NP\_009100”, “OMIM: 602391”, “HGNC: 8830”, “Ensembl: ENSG00000133027”, “Vega: OTTHUMG00000059290”, “Pharos: Q9UBM1(Tbio)”, “UniProt: Q9UBM1”) | Pfam: PEMT Herpes\_UL74 | R02056 | PE(38:6) | C00350 | PE | S-adenosyl-L-methionine:phosphatidylethanolamine N-methyltransferase | c(“RC00003 C00019\_C00021”, “RC00060 C00350\_C01241”) | C00019 + C00350 <=> C00021 + C01241 | S-Adenosyl-L-methionine + Phosphatidylethanolamine <=> S-Adenosyl-L-homocysteine + Phosphatidyl-N-methylethanolamine | 2.1.1.17 | Substrate |
| K00551 | phosphatidylethanolamine/phosphatidyl-N-methylethanolamine N-methyltransferase [EC:2.1.1.17 2.1.1.71] | 10400 | PEMT, PEAMT, PEMPT, PEMT2, PLMT, PNMT | (RefSeq) phosphatidylethanolamine N-methyltransferase | Homo sapiens (human) | c(hsa00564 = “Glycerophospholipid metabolism”, hsa01100 = “Metabolic pathways”) | c(“NCBI-GeneID: 10400”, “NCBI-ProteinID: NP\_009100”, “OMIM: 602391”, “HGNC: 8830”, “Ensembl: ENSG00000133027”, “Vega: OTTHUMG00000059290”, “Pharos: Q9UBM1(Tbio)”, “UniProt: Q9UBM1”) | Pfam: PEMT Herpes\_UL74 | R02056 | PE(40:6) | C00350 | PE | S-adenosyl-L-methionine:phosphatidylethanolamine N-methyltransferase | c(“RC00003 C00019\_C00021”, “RC00060 C00350\_C01241”) | C00019 + C00350 <=> C00021 + C01241 | S-Adenosyl-L-methionine + Phosphatidylethanolamine <=> S-Adenosyl-L-homocysteine + Phosphatidyl-N-methylethanolamine | 2.1.1.17 | Substrate |
| K00551 | phosphatidylethanolamine/phosphatidyl-N-methylethanolamine N-methyltransferase [EC:2.1.1.17 2.1.1.71] | 10400 | PEMT, PEAMT, PEMPT, PEMT2, PLMT, PNMT | (RefSeq) phosphatidylethanolamine N-methyltransferase | Homo sapiens (human) | c(hsa00564 = “Glycerophospholipid metabolism”, hsa01100 = “Metabolic pathways”) | c(“NCBI-GeneID: 10400”, “NCBI-ProteinID: NP\_009100”, “OMIM: 602391”, “HGNC: 8830”, “Ensembl: ENSG00000133027”, “Vega: OTTHUMG00000059290”, “Pharos: Q9UBM1(Tbio)”, “UniProt: Q9UBM1”) | Pfam: PEMT Herpes\_UL74 | R02056 | PE(36:3) | C00350 | PE | S-adenosyl-L-methionine:phosphatidylethanolamine N-methyltransferase | c(“RC00003 C00019\_C00021”, “RC00060 C00350\_C01241”) | C00019 + C00350 <=> C00021 + C01241 | S-Adenosyl-L-methionine + Phosphatidylethanolamine <=> S-Adenosyl-L-homocysteine + Phosphatidyl-N-methylethanolamine | 2.1.1.17 | Substrate |
| K00637 | sterol O-acyltransferase [EC:2.3.1.26] | 6646 | SOAT1, ACACT, ACAT, ACAT-1, ACAT1, SOAT, STAT | (RefSeq) sterol O-acyltransferase 1 | Homo sapiens (human) | c(hsa00100 = “Steroid biosynthesis”, hsa04979 = “Cholesterol metabolism”) | c(“NCBI-GeneID: 6646”, “NCBI-ProteinID: NP\_003092”, “OMIM: 102642”, “HGNC: 11177”, “Ensembl: ENSG00000057252”, “Vega: OTTHUMG00000035253”, “Pharos: P35610(Tchem)”, “UniProt: P35610”) | Pfam: MBOAT COX6C | R01461 | CE(22:6) | C02530 | Chol. esters | Acyl-CoA:cholesterol O-acyltransferase | c(“RC00004 C00010\_C00040”, “RC00055 C00187\_C02530”) | C00040 + C00187 <=> C00010 + C02530 | Acyl-CoA + Cholesterol <=> CoA + Cholesterol ester | 2.3.1.26 | Product |
| K00637 | sterol O-acyltransferase [EC:2.3.1.26] | 8435 | SOAT2, ACACT2, ACAT2, ARGP2 | (RefSeq) sterol O-acyltransferase 2 | Homo sapiens (human) | c(hsa00100 = “Steroid biosynthesis”, hsa04979 = “Cholesterol metabolism”) | c(“NCBI-GeneID: 8435”, “NCBI-ProteinID: NP\_003569”, “OMIM: 601311”, “HGNC: 11178”, “Ensembl: ENSG00000167780”, “Vega: OTTHUMG00000169774”, “Pharos: O75908(Tchem)”, “UniProt: O75908”) | Pfam: MBOAT MBOAT\_2 | R01461 | CE(22:6) | C02530 | Chol. esters | Acyl-CoA:cholesterol O-acyltransferase | c(“RC00004 C00010\_C00040”, “RC00055 C00187\_C02530”) | C00040 + C00187 <=> C00010 + C02530 | Acyl-CoA + Cholesterol <=> CoA + Cholesterol ester | 2.3.1.26 | Product |

###### Get gene info in short form

```
met_gene_info2=data.table::data.table(rclass_info)[,lapply(.SD, function(x) toString(unique(x))), by = 'Metabolite']
knitr::kable(head(met_gene_info2))
```

| Metabolite | orthology\_id | ORTHOLOGY | gene\_id | gene\_name | DEFINITION | ORGANISM | PATHWAY | DBLINKS | MOTIF | rxn | KEGG ID | sub\_class | Rxn\_name | RCLASS | EQUATION | EQUATION\_more | ENZYME | reactant\_product |
| --- | --- | --- | --- | --- | --- | --- | --- | --- | --- | --- | --- | --- | --- | --- | --- | --- | --- | --- |
| PE(32:1) | K00551, K00993, K01047, K01115, K01613, K05285, K05287, K05288, K05310, K08730, K13512, K13515, K13517, K13644, K14621, K16342, K16343, K16817, K16860 | phosphatidylethanolamine/phosphatidyl-N-methylethanolamine N-methyltransferase [EC:2.1.1.17 2.1.1.71], ethanolaminephosphotransferase [EC:2.7.8.1], secretory phospholipase A2 [EC:3.1.1.4], phospholipase D1/2 [EC:3.1.4.4], phosphatidylserine decarboxylase [EC:4.1.1.65], GPI ethanolamine phosphate transferase 1 [EC:2.7.-.-], GPI ethanolamine phosphate transferase 2/3 subunit F, GPI ethanolamine phosphate transferase 3 subunit O [EC:2.7.-.-], ethanolamine phosphate transferase 2 subunit G [EC:2.7.-.-], phosphatidylserine synthase 2 [EC:2.7.8.29], lysophospholipid acyltransferase [EC:2.3.1.23 2.3.1.-], lysophospholipid acyltransferase 5 [EC:2.3.1.23 2.3.1.-], lysophospholipid acyltransferase 1/2 [EC:2.3.1.51 2.3.1.-], choline/ethanolamine phosphotransferase [EC:2.7.8.1 2.7.8.2], phospholipase B1, membrane-associated [EC:3.1.1.4 3.1.1.5], cytosolic phospholipase A2 [EC:3.1.1.4], calcium-independent phospholipase A2 [EC:3.1.1.4], HRAS-like suppressor 3 [EC:3.1.1.32 3.1.1.4], phospholipase D3/4 [EC:3.1.4.4] | 10400, 85465, 26279, 30814, 391013, 50487, 5319, 5320, 5322, 64600, 81579, 8399, 84647, 5337, 5338, 23761, 23556, 5281, 84720, 54872, 81490, 254531, 10162, 129642, 154141, 10390, 151056, 100137049, 123745, 255189, 283748, 5321, 8605, 8681, 8398, 11145, 122618, 23646 | PEMT, PEAMT, PEMPT, PEMT2, PLMT, PNMT, SELENOI, EPT1, SELI, SEPI, SPG81, PLA2G2D, PLA2IID, SPLASH, sPLA2-IID, sPLA2S, PLA2G2E, GIIE\_sPLA2, sPLA2-IIE, PLA2G2C, UBXN10-AS1, PLA2G3, GIII-SPLA2, SPLA2III, sPLA2-III, PLA2G1B, PLA2, PLA2A, PPLA2, PLA2G2A, MOM1, PLA2, PLA2B, PLA2L, PLA2S, PLAS1, sPLA2, PLA2G5, FRFB, GV-PLA2, PLA2-10, hVPLA(2), PLA2G2F, GIIFsPLA2, sPLA2-IIF, PLA2G12A, GXII, PLA2G12, ROSSY, PLA2G10, GXPLA2, GXSPLA2, SPLA2, sPLA2-X, PLA2G12B, FKSG71, GXIIB, GXIIIsPLA2, PLA2G13, sPLA2-GXIIB, PLD1, CVDD, PLD2, PLD1C, PISD, DJ858B16, LIBF, PSD, PSDC, PSSC, dJ858B16.2, PIGN, MCAHS, MCAHS1, MCD4, MDC4, PIG-N, PIGF, PIGO, HPMRS2, PIGG, GPI7, LAS21, MRT53, PRO4405, RLGS1930, PTDSS2, PSS2, LPCAT4, AGPAT7, AYTL3, LPAAT-eta, LPEAT2, LPCAT3, C3F, LPCAT, LPLAT\_5, LPSAT, MBOAT5, OACT5, nessy, MBOAT2, LPAAT, LPCAT4, LPEAT, LPLAT\_2, OACT2, MBOAT1, LPEAT1, LPLAT, LPLAT\_1, LPSAT, OACT1, dJ434O11.1, CEPT1, PLB1, PLB, PLB/LIP, PLA2G4B, HsT16992, cPLA2-beta, PLA2G4E, PLA2G4F, PLA2G4FZ, PLA2G4D, cPLA2delta, PLA2G4A, GURDP, PLA2G4, cPLA2, cPLA2-alpha, PLA2G4C, CPLA2-gamma, JMJD7-PLA2G4B, HsT16992, cPLA2-beta, PLA2G6, CaI-PLA2, GVI, INAD1, IPLA2-VIA, NBIA2, NBIA2A, NBIA2B, PARK14, PLA2, PNPLA9, iPLA2, iPLA2beta, PLAAT3, AdPLA, H-REV107, H-REV107-1, HRASLS3, HREV107, HREV107-1, HREV107-3, HRSL3, PLA2G16, PLAAT-3, PLD4, C14orf175, PLD3, AD19, HU-K4, HUK4, SCA46 | (RefSeq) phosphatidylethanolamine N-methyltransferase, (RefSeq) selenoprotein I, (RefSeq) phospholipase A2 group IID, (RefSeq) phospholipase A2 group IIE, (RefSeq) phospholipase A2 group IIC, (RefSeq) phospholipase A2 group III, (RefSeq) phospholipase A2 group IB, (RefSeq) phospholipase A2 group IIA, (RefSeq) phospholipase A2 group V, (RefSeq) phospholipase A2 group IIF, (RefSeq) phospholipase A2 group XIIA, (RefSeq) phospholipase A2 group X, (RefSeq) phospholipase A2 group XIIB, (RefSeq) phospholipase D1, (RefSeq) phospholipase D2, (RefSeq) phosphatidylserine decarboxylase, (RefSeq) phosphatidylinositol glycan anchor biosynthesis class N, (RefSeq) phosphatidylinositol glycan anchor biosynthesis class F, (RefSeq) phosphatidylinositol glycan anchor biosynthesis class O, (RefSeq) phosphatidylinositol glycan anchor biosynthesis class G, (RefSeq) phosphatidylserine synthase 2, (RefSeq) lysophosphatidylcholine acyltransferase 4, (RefSeq) lysophosphatidylcholine acyltransferase 3, (RefSeq) membrane bound O-acyltransferase domain containing 2, (RefSeq) membrane bound O-acyltransferase domain containing 1, (RefSeq) choline/ethanolamine phosphotransferase 1, (RefSeq) phospholipase B1, (RefSeq) phospholipase A2 group IVB, (RefSeq) phospholipase A2 group IVE, (RefSeq) phospholipase A2 group IVF, (RefSeq) phospholipase A2 group IVD, (RefSeq) phospholipase A2 group IVA, (RefSeq) phospholipase A2 group IVC, (RefSeq) JMJD7-PLA2G4B readthrough, (RefSeq) phospholipase A2 group VI, (RefSeq) phospholipase A and acyltransferase 3, (RefSeq) phospholipase D family member 4, (RefSeq) phospholipase D family member 3 | Homo sapiens (human) | c(hsa00564 = “Glycerophospholipid metabolism”, hsa01100 = “Metabolic pathways”), c(hsa00440 = “Phosphonate and phosphinate metabolism”, hsa00564 = “Glycerophospholipid metabolism”, hsa00565 = “Ether lipid metabolism”, hsa01100 = “Metabolic pathways”), c(hsa00564 = “Glycerophospholipid metabolism”, hsa00565 = “Ether lipid metabolism”, hsa00590 = “Arachidonic acid metabolism”, hsa00591 = “Linoleic acid metabolism”, hsa00592 = “alpha-Linolenic acid metabolism”, hsa01100 = “Metabolic pathways”, hsa04014 = “Ras signaling pathway”, hsa04270 = “Vascular smooth muscle contraction”, hsa04972 = “Pancreatic secretion”, hsa04975 = “Fat digestion and absorption”), c(hsa00564 = “Glycerophospholipid metabolism”, hsa00565 = “Ether lipid metabolism”, hsa01100 = “Metabolic pathways”, hsa04014 = “Ras signaling pathway”, hsa04024 = “cAMP signaling pathway”, hsa04071 = “Sphingolipid signaling pathway”, hsa04072 = “Phospholipase D signaling pathway”, hsa04144 = “Endocytosis”, hsa04666 = “Fc gamma R-mediated phagocytosis”, hsa04724 = “Glutamatergic synapse”, hsa04912 = “GnRH signaling pathway”, hsa04928 = “Parathyroid hormone synthesis, secretion and action”, hsa05200 = “Pathways in cancer”, |  |  |  |  |  |  |  |  |  |  |  |
| hsa05212 = "P | ancreatic cancer“, hsa05231 =”Choline metabolism in cancer“), c(hsa00563 =”Glycosylphosphatidylinositol (GPI)-anchor biosynthesis“, hsa01100 =”Metabol | ic pathways“), Glycosylphosphatidylinositol (GPI)-anchor biosynthesis, c(hsa00564 =”Glycerophospholipid metabolism“, hsa00565 =”Ether lipid metabolism“, hsa01100 =”Metabolic pathways“), c(hsa00564 =”Glycerophospholipid metabolism“, hsa04216 =”Ferroptosis“), c(hsa00561 =”Glycerolipid metabolism“, hsa00564 =”Glycerophospholipid metabolism“, hsa01100 =”Metabolic pathways“), c(hsa00564 =”Glycerophospholipid metabolism“, hsa00565 =”Ether lipid metabolism“, hsa00590 =”Arachidonic acid metabolism“, hsa00591 =”Linoleic acid metabolism“, hsa00592 =”alpha-Linolenic acid metabolism“, hsa01100 =”Metabolic pathways“, hsa04977 =”Vitamin digestion and absorption“), c(hsa00564 =”Glycerophospholipid metabolism“, hsa00565 =”Ether lipid metabolism“, hsa00590 =”Arachidonic acid metabolism“, hsa00591 =”Linoleic acid metabolism“, hsa00592 =”alpha-Linolenic acid metabolism“, hsa01100 =”Metabolic pathways“, hsa04010 =”MAPK signaling pathway“, hsa04014 =”Ras signaling pathway“, hsa04072 =”Phospholipase D | signaling pathway“, hsa04217 =”Necroptosis“, hsa04270 =”Vascular smooth muscle contraction“, hsa04370 =”VEGF signaling pathway“, hsa04611 =”Platelet activation", |  |  |  |  |  |  |  |  |  |  |  |  |  |  |  |
| hsa04664 = "F | c epsilon RI signaling pathway“, hsa04666 =”Fc gamma R-mediated phagocytosis“, hsa04724 =”Glutamatergic synapse“, hsa04726 =”Serotonergic synapse", hs | a04730 = “Long-term depression”, hsa04750 = “Inflammatory mediator regulation of TRP channels”, hsa04912 = “GnRH signaling pathway”, hsa04913 = “Ovarian steroidogenesis”, hsa04921 = “Oxytocin signaling pathway”, hsa05231 = “Choline metabolism in cancer”), c(hsa00564 = “Glycerophospholipid metabolism”, hsa00565 = “Ether lipid metabolism”, hsa00590 = “Arachidonic acid metabolism”, hsa00591 = “Linoleic acid metabolism”, hsa00592 = “alpha-Linolenic acid metabolism”, hsa01100 = “Metabolic pathways”, hsa04014 = “Ras signaling pathway”, hsa04270 = “Vascular smooth muscle contraction”, hsa04666 = “Fc gamma R-mediated phagocytosis”, hsa04750 = “Inflammatory mediator regulation of TRP channels”), c(hsa00564 = “Glycerophospholipid metabolism”, hsa00565 = “Ether lipid metabolism”, hsa00590 = “Arachidonic acid metabolism”, hsa00591 = “Linoleic acid metabolism”, hsa00592 = “alpha-Linolenic acid metabolism”, hsa01100 = “Metabolic pathways”, hsa04014 = “Ras signaling pathway”, hsa04923 = "Regulation of lipolysis in adi | pocytes“) c(”NCBI-GeneID: 10400“,”NCBI-ProteinID: NP\_009100“,”OMIM: 602391“,”HGNC: 8830“,”Ensembl: ENSG00000133027“,”Vega: OTTHUMG00000059290“,”Pharos: Q9UBM1(Tbio)“,”UniProt: Q9UBM1“), c(”NCBI-GeneID: 85465“,”NCBI-ProteinID: NP\_277040“,”OMIM: 607915“,”HGNC | : 29361“,”Ensembl: ENSG00000138018“,”Vega: OTTHUMG00000151931“,”Pharos: Q9C0D9(Tbio)“,”UniProt: Q9C0D9“), c(”NCBI-GeneID: 26279“,”NCBI-ProteinID: NP\_036532“,”OMIM: 605630“,”HGNC: 9033“,”Ensembl: ENSG00000117215“,”Vega: OTTHUMG00000002701“,”Pharos: Q9UNK4(Tchem)“,”UniProt: Q9UNK4“), c(”NCBI-GeneID: 30814“,”NCBI-ProteinID: NP\_055404“,”OMIM: 618320“,”HGNC: 13414“,”Ensembl: ENSG00000188784“,”Vega: OTTHUMG00000002702“,”Pharos: Q9NZK7(Tchem)“,”UniProt: Q9NZK7“), c(”NCBI-GeneID: 391013“,”NCBI-ProteinID: NP\_001303651“,”HGNC: 9032“,”Ensembl: ENSG00000187980 ENSG00000225986“,”Vega: OTTHUMG00000002705“), c(”NCBI-GeneID: 50487“,”NCBI-ProteinID: NP\_056530“,”OMIM: 611651“,”HGNC: 17934“,”Ensembl: ENSG00000100078“,”Vega: OTTHUMG00000151255“,”Pharos: Q9NZ20(Tbio)“,”UniProt: Q9NZ20“), c(”NCBI-GeneID: 5319“,”NCBI-ProteinID: NP\_000919“,”OMIM: 172410“,”HGNC: 9030“,”Ensembl: ENSG00000170890“,”Vega: OTTHUMG00000169343“,”Pharos: P04054(Tchem)“,”UniProt: P04054“), c(”NCBI-GeneID: 5320“,”NCBI-ProteinID: NP\_000291“,”OMIM: 172411“,”HGNC: 9031“,”Ensembl: ENSG00000188257“,”Vega: OTTHUMG00000002699“,”Pharos: P14555(Tchem)“,”UniProt: P14555 A0A024RA96“), c(”NCBI-GeneID: 5322“,”NCBI-ProteinID: NP\_000920“,”OMIM: 601192“,”HGNC: 9038“,”Ensembl: ENSG00000127472“,”Vega: OTTHUMG00000002698“,”Pharos: P39877(Tchem)“,”UniProt: P39877"), c | (“NCBI-GeneID: 64600”, “NCBI-ProteinID: NP\_073730”, “OMIM: 616793”, “HGNC: 30040”, “Ensembl: ENSG00000158786”, “Vega: OTTHUMG00000002704”, “Pharos: Q9BZM2(Tchem)”, “UniProt: Q9BZM2”), c(“NCBI-GeneID: 81579”, “NCBI-ProteinID: NP\_110448”, “OMIM: 611652”, “HGNC: 18554”, “Ensembl: ENSG00000123739”, “Vega: OTTHUMG00000131915”, “Pharos: Q9BZM1(Tbio)”, “UniProt: Q9BZM1 Q542Y6”), c(“NCBI-GeneID: 8399”, “NCBI-ProteinID: NP\_003552”, “OMIM: 603603”, “HGNC: 9029”, “Ensembl: ENSG00000069764”, “Vega: OTTHUMG00000048069”, “Pharos: O15496(Tchem)”, “UniProt: O15496”), c(“NCBI-GeneID: 84647”, “NCBI-ProteinID: NP\_115951”, “OMIM: 611653”, “HGNC: 18555”, “Ensembl: ENSG00000138308”, “Vega: OTTHUMG00000018446”, “Pharos: Q9BX93(Tdark)”, “UniProt: Q9BX93”), c(“NCBI-GeneID: 5337”, “NCBI-ProteinID: NP\_002653”, “OMIM: 602382”, “HGNC: 9067”, “Ensembl: ENSG00000075651”, “Vega: OTTHUMG00000156947”, “Pharos: Q13393(Tchem)”, “UniProt: Q13393 Q59EA4”), c(“NCBI-GeneID: 5338”, “NCBI-ProteinID: NP\_002654”, “OMIM: 602384”, “HGNC: 9068”, “Ensembl: ENSG00000129219”, “Vega: OTTHUMG00000090779”, “Pharos: O14939(Tchem)”, “UniProt: O14939”), c(“NCBI-GeneID: 23761”, “NCBI-ProteinID: NP\_001313340”, “OMIM: 612770”, “HGNC: 8999”, “Ensembl: ENSG00000241878”, “Vega: OTTHUMG00000030252”, “Pharos: Q9UG56(Tbio)”, “UniProt: Q9UG56”), c(“NCBI-GeneID: 23556”, “NCBI-ProteinID: NP\_036459”, “OMIM: 606097”, “HGNC: 8967”, “Ensembl: ENSG00000197563”, “Vega: OTTHUMG00000180098”, “Pharos: O95427(Tbio)”, “UniProt: O95427 A0A024R2C3”), c(“NCBI-GeneID: 5281”, “NCBI-ProteinID: NP\_002634”, “OMIM: 600153”, “HGNC: 8962”, “Ensembl: ENSG00000151665”, "Vega: OTTHUMG00 | 000128816“,”Pharos: Q0 | 7326(Tbio)“,”UniProt: Q07326 Q6IB04“), c(”NCBI-GeneID: 84720“,”NCBI-ProteinID: NP\_116023“,”OMIM: 614730“,”HGNC: 23215“,”Ensembl: ENSG00000165282“,”Vega: OTTHUMG00000019854“,”Pharos: Q8TEQ8(Tbio)“,”UniProt: Q8TEQ8“), c(”NCBI-GeneID: 54872“,”NCBI-ProteinID: NP\_001120650“,”OMIM: 616918“,”HGNC: 25985“,”Ensembl: ENSG00000174227“,”Vega: OTTHUMG00000112457“,”Pharos: Q5H8A4(Tbio)“,”UniProt: Q5H8A4“), c(”NCBI-GeneID: 81490“,”NCBI-ProteinID: NP\_110410“,”OMIM: 612793“,”HGNC: 15463“,”Ensembl: ENSG00000174915“,”Vega: OTTHUMG00000119087“,”Pharos: Q9BVG9(Tbio)“,”UniProt: Q9BVG9 A0A024RC97“), c(”NCBI-GeneID: 254531“,”NCBI-ProteinID: NP\_705841“,”OMIM: 612039“,”HGNC: 30059“,”Ensembl: ENSG00000176454“,”Vega: OTTHUMG00000172349“,”Pharos: Q643R3(Tdark)“,”UniProt: Q643R3“), c(”NCBI-GeneID: 10162“,”NCBI-ProteinID: NP\_005759“,”OMIM: 611950“,”HGNC: 30244“,”Ensembl: ENSG00000111684“,”Vega: OTTHUMG00000168970“,”Pharos: Q6P1A2(Tbio)“,”UniProt: Q6P1A2“), c(”NCBI-GeneID: 129642“,”NCBI-ProteinID: NP\_620154“,”OMIM: 611949“,”HGNC: 25193“,”Ensembl: ENSG00000143797“,”Vega: OTTHUMG00000090363“,”Pharos: Q6ZWT7(Tdark)“,”UniProt: Q6ZWT7“), c(”NCBI-GeneID: 154141“,”NCBI-ProteinID: NP\_001073949“,”OMIM: 611732“,”HGNC: 21579“,”Ensembl: ENSG00000172197“,”Vega: OTTHUMG00000014334“,”Pharos: Q6ZNC8(Tdark)“,”UniProt: Q6ZNC8“), c(”NCBI-GeneID: 10390“,”NCBI-ProteinID: NP\_001007795“,”OMIM: 616751“,”HGNC: 24289“,”Ensembl: ENSG00000134255“,”Vega: OTTHUMG00000012357“,”Pharos: Q9Y6K0(Tbio)“,”UniProt: Q9Y6K0 A1PL14“), c(”NCBI-GeneID: 151056“,”NCBI-ProteinID: NP\_694566“,”OMIM: 610179“,”HGNC: 30041“,”Ensembl: ENSG00000163803“,”Vega: OTTHUMG00000152014“,”Pharos: Q6P1J6(Tdark)“,”UniProt: Q6P1J6 B2RWP8“), c(”NCBI-GeneID: 100137049“,”NCBI-ProteinID: NP\_001108105“,”OMIM: 606088“,”HGNC: 9036“,”Ensembl: ENSG00000243708“,”Vega: OTTHUMG00000156809“,”UniProt: P0C869“), c(”NCBI-GeneID: 123745“,”NCBI-ProteinID: NP\_001193599“,”HGNC: 24791“,”Ensembl: ENSG00000188089“,”Vega: OTTHUMG00000130371“,”Pharos: Q3MJ16(Tdark)“,”UniProt: Q3MJ16“), c(”NCBI-GeneID: 255189“,”NCBI-ProteinID: NP\_998765“,”HGNC: 27396“,”Ensembl: ENSG00000168907“,”Vega: OTTHUMG00000172782“,”Pharos: Q68DD2(Tdark)“,”UniProt: Q68DD2 A5PKZ7“), c(”NCBI-GeneID: 283748“,”NCBI-ProteinID: NP\_828848“,”OMIM: 612864“,”HGNC: 30038“,”Ensembl: ENSG00000159337“,”Vega: OTTHUMG00000172587“,”Pharos: Q86XP0(Tdark)“,”UniProt: Q86XP0“), c(”NCBI-GeneID: 5321“,”NCBI-ProteinID: NP\_077734“,”OMIM: 600522“,”HGNC: 9035“,”Ensembl: ENSG00000116711“,”Vega: OTTHUMG00000035512“,”Pharos: P47712(Tchem)“,”UniProt: P47712“), c(”NCBI-GeneID: 8605“,”NCBI-ProteinID: NP\_003697“,”OMIM: 603602“,”HGNC: 9037“,”Ensembl: ENSG00000105499“,”Vega: OTTHUMG00000183185“,”Pharos: Q9UP65(Tchem)“,”UniProt: Q9UP65 A0A024QZH0“), c(”NCBI-GeneID: 8681“,”NCBI-ProteinID: NP\_001185517“,”HGNC: 34449“,”Ensembl: ENSG00000168970“,”Vega: OTTHUMG00000044442“,”Pharos: P0C869(Tchem)“,”UniProt: P0C869“), c(”NCBI-GeneID: 8398“,”NCBI-ProteinID: NP\_003551“,”OMIM: 603604“,”HGNC: 9039“,”Ensembl: ENSG00000184381“,”Vega: OTTHUMG00000151246“,”Pharos: O60733(Tchem)“,”UniProt: O60733“), c(”NCBI-GeneID: 11145“,”NCBI-ProteinID: NP\_001121675“,”OMIM: 613867“,”HGNC: 17825“,”Ensembl: ENSG00000176485“,”Vega: OTTHUMG00000167852“,”Pharos: P53816(Tbio)“,”UniProt: P53816 A0A024R561“), c(”NCBI-GeneID: 122618“,”NCBI-ProteinID: NP\_620145“,”OMIM: 618488“,”HGNC: 23792“,”Ensembl: ENSG00000166428“,”Vega: OTTHUMG00000144167“,”Pharos: Q96BZ4(Tbio)“,”UniProt: Q96BZ4 B4DI07 B4DJQ6“), c(”NCBI-GeneID: 23646“,”NCBI-ProteinID: NP\_001026866“,”OMIM: 615698“,”HGNC: 17158“,”Ensembl: ENSG00000105223“,”Vega: OTTHUMG00000152736“,”Pharos: Q8IV08(Tbio)“,”U | niProt: Q8IV08 A0A024R0Q4“) Pfam: PEMT Herpes\_UL74, Pfam: CDP-OH\_P\_transf, Pfam: Phospholip\_A2\_1 Parvo\_coat\_N DUF5460, Pfam: Phospholip\_A2\_1, Pfam: Phospholip\_A2\_2 Chromadorea\_ALT, Pfam: Phospholip\_A2\_1 Parvo\_coat\_N Phospholip\_A2\_2 DUF1644, Pfam: Phospholip\_A2\_1 Phospholip\_A2\_2, Pfam: Phospholip\_A2\_1 Parvo\_coat\_N, Pfam: PLA2G12, Pfam: PLDc\_2 PX PLDc PH DUF3785, Pfam: PX PLDc\_2 PLDc PH PH\_2, Pfam: PS\_Dcarbxylase, Pfam: PigN Phosphodiest Sulfatase Metalloenzyme, Pfam: PIG-F, Pfam: Phosphodiest Metalloenzyme Sulfatase DUF1501, Pfam: Phosphodiest Metalloenzyme, Pfam: PSS TctB, Pfam: Acyltransferase JmjN, Pfam: MBOAT DUF5659, Pfam: MBOAT, Pfam: Lipase\_GDSL Lipase\_GDSL\_2, Pfam: PLA2\_B cPLA2\_C2 C2, Pfam: cPLA2\_C2 PLA2\_B C2, Pfam: PLA2\_B C2, Pfam: PLA2\_B, Pfam: Cupin\_8 PLA2\_B cPLA2\_C2 C2, Pfam: Ank\_2 Ank\_4 Ank\_5 Ank Ank\_3 Patatin, Pfam: LRAT Calici\_PP\_N Peptidase\_C97 Churchill, Pfam: PLDc\_3 PLDc\_2 PLDc R02056, R02057, R02053, R02051, R02055, R12351, R05924, R05923, R08107, R07376, R04480, R02054 C00350 PE S-adenosyl-L-methionine:phosphatidylethanolamine N-methyltransferase, CDPethanolamine:1,2-diacylglycerol ethanolaminephosphotransferase, phosphatidylethanolamine 2-acylhydrolase, phosphatidylethanolamine phosphatidohydrolase, Phsophatidyl-L-serine carboxy-lyase, NULL, L-1-phosphatidylethanolamine:L-serine phosphatidyltransferase, Acyl-CoA:1-acyl-sn-glycero-3-phosphoethanolamine O-acyltransferase, phosphatidylethanolamine 1-acylhydrolase c(”RC00003 C00019\_C00021“,”RC00060 C00350\_C01241“), c(”RC00002 C00055\_C00570“,”RC00017 C00350\_C00641“), c(”RC00037 C00350\_C04438“,”RC00094 C00162\_C00350“), c(”RC00017 C00189\_C00350“,”RC00425 C00350\_C00416“), RC00299 C00350\_C02737, NULL, RC00017 C00350\_C00641, c(”RC00017 C00065\_C02737 C00189\_C00350“,”RC02950 C00350\_C02737“), c(”RC00004 C00010\_C00040“,”RC00037 C00350\_C04438“), c(”RC00020 C00162\_C00350“,”RC00041 C00350\_C05973") C00019 + C00350 <=> C00021 + C01241, C00570 + C00641 <=> C00055 + C00350, C00350 + C00001 <=> C04438 + C00162, C00350 + C00001 <=> C00189 + C00416, C02737 <=> C00350 + C00011, G13128 + C00350 <=> G00148 + C00641, C00350 + G00141 <=> C00641 + G00151, C00350 + G00140 <=> C00641 + G00141, C00350 + G00149 <=> C00641 + G13044, C00350 + C00065 <=> C02737 + C00189, C00350 + C00010 <=> C04438 + C00040, C00350 + C00001 <=> C05973 + C00162 S-Adenosyl-L-methionine + Phosphatidylethanolamine <=> S-Adenosyl-L-homocysteine + Phosphatidyl-N-methylethanolamine, CDP-ethanolamine + 1,2-Diacyl-sn-glycerol <=> CMP + Phosphatidylethanolamine, Phosphatidylethanolamine + H2O <=> 1-Acyl-sn-glycero-3-phosphoethanolamine + Fatty acid, Phosphatidylethanolamine + H2O <=> Ethanolamine + Phosphatidate, Phosphatidylserine <=> Phosphatidylethanolamine + CO2, G13128 + Phosphatidylethanolamine <=> G00148 + 1,2-Diacyl-sn-glycerol, Phosphatidylethanolamine + G00141 <=> 1,2-Diacyl-sn-glycerol + G00151, Phosphatidylethanolamine + G00140 <=> 1,2-Diacyl-sn-glycerol + G00141, Phosphatidylethanolamine + G00149 <=> 1,2-Diacyl-sn-glycerol + G13044, Phosphatidylethanolamine + L-Serine <=> Phosphatidylserine + Ethanolamine, Phosphatidylethanolamine + CoA <=> 1-Acyl-sn-glycero-3-phosphoethanolamine + Acyl-CoA, Phosphatidylethanolamine + H2O <=> 2-Acyl-sn-glycero-3-phosphoethanolamine + Fatty acid 2.1.1.17, 2.7.8.1, 3.1.1.4, 3.1.4.4, 4.1.1.65, 2.7.-.-, NULL, 2.7.8.29, 2.3.1.23, 3.1.1.32 Substrate, Product |  |  |  |  |  |  |  |  |  |  |
| PE(38:6) | K00551, K00993, K01047, K01115, K01613, K05285, K05287, K05288, K05310, K08730, K13512, K13515, K13517, K13644, K14621, K16342, K16343, K16817, K16860 | phosphatidylethanolamine/phosphatidyl-N-methylethanolamine N-methyltransferase [EC:2.1.1.17 2.1.1.71], ethanolaminephosphotransferase [EC:2.7.8.1], secretory phospholipase A2 [EC:3.1.1.4], phospholipase D1/2 [EC:3.1.4.4], phosphatidylserine decarboxylase [EC:4.1.1.65], GPI ethanolamine phosphate transferase 1 [EC:2.7.-.-], GPI ethanolamine phosphate transferase 2/3 subunit F, GPI ethanolamine phosphate transferase 3 subunit O [EC:2.7.-.-], ethanolamine phosphate transferase 2 subunit G [EC:2.7.-.-], phosphatidylserine synthase 2 [EC:2.7.8.29], lysophospholipid acyltransferase [EC:2.3.1.23 2.3.1.-], lysophospholipid acyltransferase 5 [EC:2.3.1.23 2.3.1.-], lysophospholipid acyltransferase 1/2 [EC:2.3.1.51 2.3.1.-], choline/ethanolamine phosphotransferase [EC:2.7.8.1 2.7.8.2], phospholipase B1, membrane-associated [EC:3.1.1.4 3.1.1.5], cytosolic phospholipase A2 [EC:3.1.1.4], calcium-independent phospholipase A2 [EC:3.1.1.4], HRAS-like suppressor 3 [EC:3.1.1.32 3.1.1.4], phospholipase D3/4 [EC:3.1.4.4] | 10400, 85465, 26279, 30814, 391013, 50487, 5319, 5320, 5322, 64600, 81579, 8399, 84647, 5337, 5338, 23761, 23556, 5281, 84720, 54872, 81490, 254531, 10162, 129642, 154141, 10390, 151056, 100137049, 123745, 255189, 283748, 5321, 8605, 8681, 8398, 11145, 122618, 23646 | PEMT, PEAMT, PEMPT, PEMT2, PLMT, PNMT, SELENOI, EPT1, SELI, SEPI, SPG81, PLA2G2D, PLA2IID, SPLASH, sPLA2-IID, sPLA2S, PLA2G2E, GIIE\_sPLA2, sPLA2-IIE, PLA2G2C, UBXN10-AS1, PLA2G3, GIII-SPLA2, SPLA2III, sPLA2-III, PLA2G1B, PLA2, PLA2A, PPLA2, PLA2G2A, MOM1, PLA2, PLA2B, PLA2L, PLA2S, PLAS1, sPLA2, PLA2G5, FRFB, GV-PLA2, PLA2-10, hVPLA(2), PLA2G2F, GIIFsPLA2, sPLA2-IIF, PLA2G12A, GXII, PLA2G12, ROSSY, PLA2G10, GXPLA2, GXSPLA2, SPLA2, sPLA2-X, PLA2G12B, FKSG71, GXIIB, GXIIIsPLA2, PLA2G13, sPLA2-GXIIB, PLD1, CVDD, PLD2, PLD1C, PISD, DJ858B16, LIBF, PSD, PSDC, PSSC, dJ858B16.2, PIGN, MCAHS, MCAHS1, MCD4, MDC4, PIG-N, PIGF, PIGO, HPMRS2, PIGG, GPI7, LAS21, MRT53, PRO4405, RLGS1930, PTDSS2, PSS2, LPCAT4, AGPAT7, AYTL3, LPAAT-eta, LPEAT2, LPCAT3, C3F, LPCAT, LPLAT\_5, LPSAT, MBOAT5, OACT5, nessy, MBOAT2, LPAAT, LPCAT4, LPEAT, LPLAT\_2, OACT2, MBOAT1, LPEAT1, LPLAT, LPLAT\_1, LPSAT, OACT1, dJ434O11.1, CEPT1, PLB1, PLB, PLB/LIP, PLA2G4B, HsT16992, cPLA2-beta, PLA2G4E, PLA2G4F, PLA2G4FZ, PLA2G4D, cPLA2delta, PLA2G4A, GURDP, PLA2G4, cPLA2, cPLA2-alpha, PLA2G4C, CPLA2-gamma, JMJD7-PLA2G4B, HsT16992, cPLA2-beta, PLA2G6, CaI-PLA2, GVI, INAD1, IPLA2-VIA, NBIA2, NBIA2A, NBIA2B, PARK14, PLA2, PNPLA9, iPLA2, iPLA2beta, PLAAT3, AdPLA, H-REV107, H-REV107-1, HRASLS3, HREV107, HREV107-1, HREV107-3, HRSL3, PLA2G16, PLAAT-3, PLD4, C14orf175, PLD3, AD19, HU-K4, HUK4, SCA46 | (RefSeq) phosphatidylethanolamine N-methyltransferase, (RefSeq) selenoprotein I, (RefSeq) phospholipase A2 group IID, (RefSeq) phospholipase A2 group IIE, (RefSeq) phospholipase A2 group IIC, (RefSeq) phospholipase A2 group III, (RefSeq) phospholipase A2 group IB, (RefSeq) phospholipase A2 group IIA, (RefSeq) phospholipase A2 group V, (RefSeq) phospholipase A2 group IIF, (RefSeq) phospholipase A2 group XIIA, (RefSeq) phospholipase A2 group X, (RefSeq) phospholipase A2 group XIIB, (RefSeq) phospholipase D1, (RefSeq) phospholipase D2, (RefSeq) phosphatidylserine decarboxylase, (RefSeq) phosphatidylinositol glycan anchor biosynthesis class N, (RefSeq) phosphatidylinositol glycan anchor biosynthesis class F, (RefSeq) phosphatidylinositol glycan anchor biosynthesis class O, (RefSeq) phosphatidylinositol glycan anchor biosynthesis class G, (RefSeq) phosphatidylserine synthase 2, (RefSeq) lysophosphatidylcholine acyltransferase 4, (RefSeq) lysophosphatidylcholine acyltransferase 3, (RefSeq) membrane bound O-acyltransferase domain containing 2, (RefSeq) membrane bound O-acyltransferase domain containing 1, (RefSeq) choline/ethanolamine phosphotransferase 1, (RefSeq) phospholipase B1, (RefSeq) phospholipase A2 group IVB, (RefSeq) phospholipase A2 group IVE, (RefSeq) phospholipase A2 group IVF, (RefSeq) phospholipase A2 group IVD, (RefSeq) phospholipase A2 group IVA, (RefSeq) phospholipase A2 group IVC, (RefSeq) JMJD7-PLA2G4B readthrough, (RefSeq) phospholipase A2 group VI, (RefSeq) phospholipase A and acyltransferase 3, (RefSeq) phospholipase D family member 4, (RefSeq) phospholipase D family member 3 | Homo sapiens (human) | c(hsa00564 = “Glycerophospholipid metabolism”, hsa01100 = “Metabolic pathways”), c(hsa00440 = “Phosphonate and phosphinate metabolism”, hsa00564 = “Glycerophospholipid metabolism”, hsa00565 = “Ether lipid metabolism”, hsa01100 = “Metabolic pathways”), c(hsa00564 = “Glycerophospholipid metabolism”, hsa00565 = “Ether lipid metabolism”, hsa00590 = “Arachidonic acid metabolism”, hsa00591 = “Linoleic acid metabolism”, hsa00592 = “alpha-Linolenic acid metabolism”, hsa01100 = “Metabolic pathways”, hsa04014 = “Ras signaling pathway”, hsa04270 = “Vascular smooth muscle contraction”, hsa04972 = “Pancreatic secretion”, hsa04975 = “Fat digestion and absorption”), c(hsa00564 = “Glycerophospholipid metabolism”, hsa00565 = “Ether lipid metabolism”, hsa01100 = “Metabolic pathways”, hsa04014 = “Ras signaling pathway”, hsa04024 = “cAMP signaling pathway”, hsa04071 = “Sphingolipid signaling pathway”, hsa04072 = “Phospholipase D signaling pathway”, hsa04144 = “Endocytosis”, hsa04666 = “Fc gamma R-mediated phagocytosis”, hsa04724 = “Glutamatergic synapse”, hsa04912 = “GnRH signaling pathway”, hsa04928 = “Parathyroid hormone synthesis, secretion and action”, hsa05200 = “Pathways in cancer”, |  |  |  |  |  |  |  |  |  |  |  |
| hsa05212 = "P | ancreatic cancer“, hsa05231 =”Choline metabolism in cancer“), c(hsa00563 =”Glycosylphosphatidylinositol (GPI)-anchor biosynthesis“, hsa01100 =”Metabol | ic pathways“), Glycosylphosphatidylinositol (GPI)-anchor biosynthesis, c(hsa00564 =”Glycerophospholipid metabolism“, hsa00565 =”Ether lipid metabolism“, hsa01100 =”Metabolic pathways“), c(hsa00564 =”Glycerophospholipid metabolism“, hsa04216 =”Ferroptosis“), c(hsa00561 =”Glycerolipid metabolism“, hsa00564 =”Glycerophospholipid metabolism“, hsa01100 =”Metabolic pathways“), c(hsa00564 =”Glycerophospholipid metabolism“, hsa00565 =”Ether lipid metabolism“, hsa00590 =”Arachidonic acid metabolism“, hsa00591 =”Linoleic acid metabolism“, hsa00592 =”alpha-Linolenic acid metabolism“, hsa01100 =”Metabolic pathways“, hsa04977 =”Vitamin digestion and absorption“), c(hsa00564 =”Glycerophospholipid metabolism“, hsa00565 =”Ether lipid metabolism“, hsa00590 =”Arachidonic acid metabolism“, hsa00591 =”Linoleic acid metabolism“, hsa00592 =”alpha-Linolenic acid metabolism“, hsa01100 =”Metabolic pathways“, hsa04010 =”MAPK signaling pathway“, hsa04014 =”Ras signaling pathway“, hsa04072 =”Phospholipase D | signaling pathway“, hsa04217 =”Necroptosis“, hsa04270 =”Vascular smooth muscle contraction“, hsa04370 =”VEGF signaling pathway“, hsa04611 =”Platelet activation", |  |  |  |  |  |  |  |  |  |  |  |  |  |  |  |
| hsa04664 = "F | c epsilon RI signaling pathway“, hsa04666 =”Fc gamma R-mediated phagocytosis“, hsa04724 =”Glutamatergic synapse“, hsa04726 =”Serotonergic synapse", hs | a04730 = “Long-term depression”, hsa04750 = “Inflammatory mediator regulation of TRP channels”, hsa04912 = “GnRH signaling pathway”, hsa04913 = “Ovarian steroidogenesis”, hsa04921 = “Oxytocin signaling pathway”, hsa05231 = “Choline metabolism in cancer”), c(hsa00564 = “Glycerophospholipid metabolism”, hsa00565 = “Ether lipid metabolism”, hsa00590 = “Arachidonic acid metabolism”, hsa00591 = “Linoleic acid metabolism”, hsa00592 = “alpha-Linolenic acid metabolism”, hsa01100 = “Metabolic pathways”, hsa04014 = “Ras signaling pathway”, hsa04270 = “Vascular smooth muscle contraction”, hsa04666 = “Fc gamma R-mediated phagocytosis”, hsa04750 = “Inflammatory mediator regulation of TRP channels”), c(hsa00564 = “Glycerophospholipid metabolism”, hsa00565 = “Ether lipid metabolism”, hsa00590 = “Arachidonic acid metabolism”, hsa00591 = “Linoleic acid metabolism”, hsa00592 = “alpha-Linolenic acid metabolism”, hsa01100 = “Metabolic pathways”, hsa04014 = “Ras signaling pathway”, hsa04923 = "Regulation of lipolysis in adi | pocytes“) c(”NCBI-GeneID: 10400“,”NCBI-ProteinID: NP\_009100“,”OMIM: 602391“,”HGNC: 8830“,”Ensembl: ENSG00000133027“,”Vega: OTTHUMG00000059290“,”Pharos: Q9UBM1(Tbio)“,”UniProt: Q9UBM1“), c(”NCBI-GeneID: 85465“,”NCBI-ProteinID: NP\_277040“,”OMIM: 607915“,”HGNC | : 29361“,”Ensembl: ENSG00000138018“,”Vega: OTTHUMG00000151931“,”Pharos: Q9C0D9(Tbio)“,”UniProt: Q9C0D9“), c(”NCBI-GeneID: 26279“,”NCBI-ProteinID: NP\_036532“,”OMIM: 605630“,”HGNC: 9033“,”Ensembl: ENSG00000117215“,”Vega: OTTHUMG00000002701“,”Pharos: Q9UNK4(Tchem)“,”UniProt: Q9UNK4“), c(”NCBI-GeneID: 30814“,”NCBI-ProteinID: NP\_055404“,”OMIM: 618320“,”HGNC: 13414“,”Ensembl: ENSG00000188784“,”Vega: OTTHUMG00000002702“,”Pharos: Q9NZK7(Tchem)“,”UniProt: Q9NZK7“), c(”NCBI-GeneID: 391013“,”NCBI-ProteinID: NP\_001303651“,”HGNC: 9032“,”Ensembl: ENSG00000187980 ENSG00000225986“,”Vega: OTTHUMG00000002705“), c(”NCBI-GeneID: 50487“,”NCBI-ProteinID: NP\_056530“,”OMIM: 611651“,”HGNC: 17934“,”Ensembl: ENSG00000100078“,”Vega: OTTHUMG00000151255“,”Pharos: Q9NZ20(Tbio)“,”UniProt: Q9NZ20“), c(”NCBI-GeneID: 5319“,”NCBI-ProteinID: NP\_000919“,”OMIM: 172410“,”HGNC: 9030“,”Ensembl: ENSG00000170890“,”Vega: OTTHUMG00000169343“,”Pharos: P04054(Tchem)“,”UniProt: P04054“), c(”NCBI-GeneID: 5320“,”NCBI-ProteinID: NP\_000291“,”OMIM: 172411“,”HGNC: 9031“,”Ensembl: ENSG00000188257“,”Vega: OTTHUMG00000002699“,”Pharos: P14555(Tchem)“,”UniProt: P14555 A0A024RA96“), c(”NCBI-GeneID: 5322“,”NCBI-ProteinID: NP\_000920“,”OMIM: 601192“,”HGNC: 9038“,”Ensembl: ENSG00000127472“,”Vega: OTTHUMG00000002698“,”Pharos: P39877(Tchem)“,”UniProt: P39877"), c | (“NCBI-GeneID: 64600”, “NCBI-ProteinID: NP\_073730”, “OMIM: 616793”, “HGNC: 30040”, “Ensembl: ENSG00000158786”, “Vega: OTTHUMG00000002704”, “Pharos: Q9BZM2(Tchem)”, “UniProt: Q9BZM2”), c(“NCBI-GeneID: 81579”, “NCBI-ProteinID: NP\_110448”, “OMIM: 611652”, “HGNC: 18554”, “Ensembl: ENSG00000123739”, “Vega: OTTHUMG00000131915”, “Pharos: Q9BZM1(Tbio)”, “UniProt: Q9BZM1 Q542Y6”), c(“NCBI-GeneID: 8399”, “NCBI-ProteinID: NP\_003552”, “OMIM: 603603”, “HGNC: 9029”, “Ensembl: ENSG00000069764”, “Vega: OTTHUMG00000048069”, “Pharos: O15496(Tchem)”, “UniProt: O15496”), c(“NCBI-GeneID: 84647”, “NCBI-ProteinID: NP\_115951”, “OMIM: 611653”, “HGNC: 18555”, “Ensembl: ENSG00000138308”, “Vega: OTTHUMG00000018446”, “Pharos: Q9BX93(Tdark)”, “UniProt: Q9BX93”), c(“NCBI-GeneID: 5337”, “NCBI-ProteinID: NP\_002653”, “OMIM: 602382”, “HGNC: 9067”, “Ensembl: ENSG00000075651”, “Vega: OTTHUMG00000156947”, “Pharos: Q13393(Tchem)”, “UniProt: Q13393 Q59EA4”), c(“NCBI-GeneID: 5338”, “NCBI-ProteinID: NP\_002654”, “OMIM: 602384”, “HGNC: 9068”, “Ensembl: ENSG00000129219”, “Vega: OTTHUMG00000090779”, “Pharos: O14939(Tchem)”, “UniProt: O14939”), c(“NCBI-GeneID: 23761”, “NCBI-ProteinID: NP\_001313340”, “OMIM: 612770”, “HGNC: 8999”, “Ensembl: ENSG00000241878”, “Vega: OTTHUMG00000030252”, “Pharos: Q9UG56(Tbio)”, “UniProt: Q9UG56”), c(“NCBI-GeneID: 23556”, “NCBI-ProteinID: NP\_036459”, “OMIM: 606097”, “HGNC: 8967”, “Ensembl: ENSG00000197563”, “Vega: OTTHUMG00000180098”, “Pharos: O95427(Tbio)”, “UniProt: O95427 A0A024R2C3”), c(“NCBI-GeneID: 5281”, “NCBI-ProteinID: NP\_002634”, “OMIM: 600153”, “HGNC: 8962”, “Ensembl: ENSG00000151665”, "Vega: OTTHUMG00 | 000128816“,”Pharos: Q0 | 7326(Tbio)“,”UniProt: Q07326 Q6IB04“), c(”NCBI-GeneID: 84720“,”NCBI-ProteinID: NP\_116023“,”OMIM: 614730“,”HGNC: 23215“,”Ensembl: ENSG00000165282“,”Vega: OTTHUMG00000019854“,”Pharos: Q8TEQ8(Tbio)“,”UniProt: Q8TEQ8“), c(”NCBI-GeneID: 54872“,”NCBI-ProteinID: NP\_001120650“,”OMIM: 616918“,”HGNC: 25985“,”Ensembl: ENSG00000174227“,”Vega: OTTHUMG00000112457“,”Pharos: Q5H8A4(Tbio)“,”UniProt: Q5H8A4“), c(”NCBI-GeneID: 81490“,”NCBI-ProteinID: NP\_110410“,”OMIM: 612793“,”HGNC: 15463“,”Ensembl: ENSG00000174915“,”Vega: OTTHUMG00000119087“,”Pharos: Q9BVG9(Tbio)“,”UniProt: Q9BVG9 A0A024RC97“), c(”NCBI-GeneID: 254531“,”NCBI-ProteinID: NP\_705841“,”OMIM: 612039“,”HGNC: 30059“,”Ensembl: ENSG00000176454“,”Vega: OTTHUMG00000172349“,”Pharos: Q643R3(Tdark)“,”UniProt: Q643R3“), c(”NCBI-GeneID: 10162“,”NCBI-ProteinID: NP\_005759“,”OMIM: 611950“,”HGNC: 30244“,”Ensembl: ENSG00000111684“,”Vega: OTTHUMG00000168970“,”Pharos: Q6P1A2(Tbio)“,”UniProt: Q6P1A2“), c(”NCBI-GeneID: 129642“,”NCBI-ProteinID: NP\_620154“,”OMIM: 611949“,”HGNC: 25193“,”Ensembl: ENSG00000143797“,”Vega: OTTHUMG00000090363“,”Pharos: Q6ZWT7(Tdark)“,”UniProt: Q6ZWT7“), c(”NCBI-GeneID: 154141“,”NCBI-ProteinID: NP\_001073949“,”OMIM: 611732“,”HGNC: 21579“,”Ensembl: ENSG00000172197“,”Vega: OTTHUMG00000014334“,”Pharos: Q6ZNC8(Tdark)“,”UniProt: Q6ZNC8“), c(”NCBI-GeneID: 10390“,”NCBI-ProteinID: NP\_001007795“,”OMIM: 616751“,”HGNC: 24289“,”Ensembl: ENSG00000134255“,”Vega: OTTHUMG00000012357“,”Pharos: Q9Y6K0(Tbio)“,”UniProt: Q9Y6K0 A1PL14“), c(”NCBI-GeneID: 151056“,”NCBI-ProteinID: NP\_694566“,”OMIM: 610179“,”HGNC: 30041“,”Ensembl: ENSG00000163803“,”Vega: OTTHUMG00000152014“,”Pharos: Q6P1J6(Tdark)“,”UniProt: Q6P1J6 B2RWP8“), c(”NCBI-GeneID: 100137049“,”NCBI-ProteinID: NP\_001108105“,”OMIM: 606088“,”HGNC: 9036“,”Ensembl: ENSG00000243708“,”Vega: OTTHUMG00000156809“,”UniProt: P0C869“), c(”NCBI-GeneID: 123745“,”NCBI-ProteinID: NP\_001193599“,”HGNC: 24791“,”Ensembl: ENSG00000188089“,”Vega: OTTHUMG00000130371“,”Pharos: Q3MJ16(Tdark)“,”UniProt: Q3MJ16“), c(”NCBI-GeneID: 255189“,”NCBI-ProteinID: NP\_998765“,”HGNC: 27396“,”Ensembl: ENSG00000168907“,”Vega: OTTHUMG00000172782“,”Pharos: Q68DD2(Tdark)“,”UniProt: Q68DD2 A5PKZ7“), c(”NCBI-GeneID: 283748“,”NCBI-ProteinID: NP\_828848“,”OMIM: 612864“,”HGNC: 30038“,”Ensembl: ENSG00000159337“,”Vega: OTTHUMG00000172587“,”Pharos: Q86XP0(Tdark)“,”UniProt: Q86XP0“), c(”NCBI-GeneID: 5321“,”NCBI-ProteinID: NP\_077734“,”OMIM: 600522“,”HGNC: 9035“,”Ensembl: ENSG00000116711“,”Vega: OTTHUMG00000035512“,”Pharos: P47712(Tchem)“,”UniProt: P47712“), c(”NCBI-GeneID: 8605“,”NCBI-ProteinID: NP\_003697“,”OMIM: 603602“,”HGNC: 9037“,”Ensembl: ENSG00000105499“,”Vega: OTTHUMG00000183185“,”Pharos: Q9UP65(Tchem)“,”UniProt: Q9UP65 A0A024QZH0“), c(”NCBI-GeneID: 8681“,”NCBI-ProteinID: NP\_001185517“,”HGNC: 34449“,”Ensembl: ENSG00000168970“,”Vega: OTTHUMG00000044442“,”Pharos: P0C869(Tchem)“,”UniProt: P0C869“), c(”NCBI-GeneID: 8398“,”NCBI-ProteinID: NP\_003551“,”OMIM: 603604“,”HGNC: 9039“,”Ensembl: ENSG00000184381“,”Vega: OTTHUMG00000151246“,”Pharos: O60733(Tchem)“,”UniProt: O60733“), c(”NCBI-GeneID: 11145“,”NCBI-ProteinID: NP\_001121675“,”OMIM: 613867“,”HGNC: 17825“,”Ensembl: ENSG00000176485“,”Vega: OTTHUMG00000167852“,”Pharos: P53816(Tbio)“,”UniProt: P53816 A0A024R561“), c(”NCBI-GeneID: 122618“,”NCBI-ProteinID: NP\_620145“,”OMIM: 618488“,”HGNC: 23792“,”Ensembl: ENSG00000166428“,”Vega: OTTHUMG00000144167“,”Pharos: Q96BZ4(Tbio)“,”UniProt: Q96BZ4 B4DI07 B4DJQ6“), c(”NCBI-GeneID: 23646“,”NCBI-ProteinID: NP\_001026866“,”OMIM: 615698“,”HGNC: 17158“,”Ensembl: ENSG00000105223“,”Vega: OTTHUMG00000152736“,”Pharos: Q8IV08(Tbio)“,”U | niProt: Q8IV08 A0A024R0Q4“) Pfam: PEMT Herpes\_UL74, Pfam: CDP-OH\_P\_transf, Pfam: Phospholip\_A2\_1 Parvo\_coat\_N DUF5460, Pfam: Phospholip\_A2\_1, Pfam: Phospholip\_A2\_2 Chromadorea\_ALT, Pfam: Phospholip\_A2\_1 Parvo\_coat\_N Phospholip\_A2\_2 DUF1644, Pfam: Phospholip\_A2\_1 Phospholip\_A2\_2, Pfam: Phospholip\_A2\_1 Parvo\_coat\_N, Pfam: PLA2G12, Pfam: PLDc\_2 PX PLDc PH DUF3785, Pfam: PX PLDc\_2 PLDc PH PH\_2, Pfam: PS\_Dcarbxylase, Pfam: PigN Phosphodiest Sulfatase Metalloenzyme, Pfam: PIG-F, Pfam: Phosphodiest Metalloenzyme Sulfatase DUF1501, Pfam: Phosphodiest Metalloenzyme, Pfam: PSS TctB, Pfam: Acyltransferase JmjN, Pfam: MBOAT DUF5659, Pfam: MBOAT, Pfam: Lipase\_GDSL Lipase\_GDSL\_2, Pfam: PLA2\_B cPLA2\_C2 C2, Pfam: cPLA2\_C2 PLA2\_B C2, Pfam: PLA2\_B C2, Pfam: PLA2\_B, Pfam: Cupin\_8 PLA2\_B cPLA2\_C2 C2, Pfam: Ank\_2 Ank\_4 Ank\_5 Ank Ank\_3 Patatin, Pfam: LRAT Calici\_PP\_N Peptidase\_C97 Churchill, Pfam: PLDc\_3 PLDc\_2 PLDc R02056, R02057, R02053, R02051, R02055, R12351, R05924, R05923, R08107, R07376, R04480, R02054 C00350 PE S-adenosyl-L-methionine:phosphatidylethanolamine N-methyltransferase, CDPethanolamine:1,2-diacylglycerol ethanolaminephosphotransferase, phosphatidylethanolamine 2-acylhydrolase, phosphatidylethanolamine phosphatidohydrolase, Phsophatidyl-L-serine carboxy-lyase, NULL, L-1-phosphatidylethanolamine:L-serine phosphatidyltransferase, Acyl-CoA:1-acyl-sn-glycero-3-phosphoethanolamine O-acyltransferase, phosphatidylethanolamine 1-acylhydrolase c(”RC00003 C00019\_C00021“,”RC00060 C00350\_C01241“), c(”RC00002 C00055\_C00570“,”RC00017 C00350\_C00641“), c(”RC00037 C00350\_C04438“,”RC00094 C00162\_C00350“), c(”RC00017 C00189\_C00350“,”RC00425 C00350\_C00416“), RC00299 C00350\_C02737, NULL, RC00017 C00350\_C00641, c(”RC00017 C00065\_C02737 C00189\_C00350“,”RC02950 C00350\_C02737“), c(”RC00004 C00010\_C00040“,”RC00037 C00350\_C04438“), c(”RC00020 C00162\_C00350“,”RC00041 C00350\_C05973") C00019 + C00350 <=> C00021 + C01241, C00570 + C00641 <=> C00055 + C00350, C00350 + C00001 <=> C04438 + C00162, C00350 + C00001 <=> C00189 + C00416, C02737 <=> C00350 + C00011, G13128 + C00350 <=> G00148 + C00641, C00350 + G00141 <=> C00641 + G00151, C00350 + G00140 <=> C00641 + G00141, C00350 + G00149 <=> C00641 + G13044, C00350 + C00065 <=> C02737 + C00189, C00350 + C00010 <=> C04438 + C00040, C00350 + C00001 <=> C05973 + C00162 S-Adenosyl-L-methionine + Phosphatidylethanolamine <=> S-Adenosyl-L-homocysteine + Phosphatidyl-N-methylethanolamine, CDP-ethanolamine + 1,2-Diacyl-sn-glycerol <=> CMP + Phosphatidylethanolamine, Phosphatidylethanolamine + H2O <=> 1-Acyl-sn-glycero-3-phosphoethanolamine + Fatty acid, Phosphatidylethanolamine + H2O <=> Ethanolamine + Phosphatidate, Phosphatidylserine <=> Phosphatidylethanolamine + CO2, G13128 + Phosphatidylethanolamine <=> G00148 + 1,2-Diacyl-sn-glycerol, Phosphatidylethanolamine + G00141 <=> 1,2-Diacyl-sn-glycerol + G00151, Phosphatidylethanolamine + G00140 <=> 1,2-Diacyl-sn-glycerol + G00141, Phosphatidylethanolamine + G00149 <=> 1,2-Diacyl-sn-glycerol + G13044, Phosphatidylethanolamine + L-Serine <=> Phosphatidylserine + Ethanolamine, Phosphatidylethanolamine + CoA <=> 1-Acyl-sn-glycero-3-phosphoethanolamine + Acyl-CoA, Phosphatidylethanolamine + H2O <=> 2-Acyl-sn-glycero-3-phosphoethanolamine + Fatty acid 2.1.1.17, 2.7.8.1, 3.1.1.4, 3.1.4.4, 4.1.1.65, 2.7.-.-, NULL, 2.7.8.29, 2.3.1.23, 3.1.1.32 Substrate, Product |  |  |  |  |  |  |  |  |  |  |
| PE(40:6) | K00551, K00993, K01047, K01115, K01613, K05285, K05287, K05288, K05310, K08730, K13512, K13515, K13517, K13644, K14621, K16342, K16343, K16817, K16860 | phosphatidylethanolamine/phosphatidyl-N-methylethanolamine N-methyltransferase [EC:2.1.1.17 2.1.1.71], ethanolaminephosphotransferase [EC:2.7.8.1], secretory phospholipase A2 [EC:3.1.1.4], phospholipase D1/2 [EC:3.1.4.4], phosphatidylserine decarboxylase [EC:4.1.1.65], GPI ethanolamine phosphate transferase 1 [EC:2.7.-.-], GPI ethanolamine phosphate transferase 2/3 subunit F, GPI ethanolamine phosphate transferase 3 subunit O [EC:2.7.-.-], ethanolamine phosphate transferase 2 subunit G [EC:2.7.-.-], phosphatidylserine synthase 2 [EC:2.7.8.29], lysophospholipid acyltransferase [EC:2.3.1.23 2.3.1.-], lysophospholipid acyltransferase 5 [EC:2.3.1.23 2.3.1.-], lysophospholipid acyltransferase 1/2 [EC:2.3.1.51 2.3.1.-], choline/ethanolamine phosphotransferase [EC:2.7.8.1 2.7.8.2], phospholipase B1, membrane-associated [EC:3.1.1.4 3.1.1.5], cytosolic phospholipase A2 [EC:3.1.1.4], calcium-independent phospholipase A2 [EC:3.1.1.4], HRAS-like suppressor 3 [EC:3.1.1.32 3.1.1.4], phospholipase D3/4 [EC:3.1.4.4] | 10400, 85465, 26279, 30814, 391013, 50487, 5319, 5320, 5322, 64600, 81579, 8399, 84647, 5337, 5338, 23761, 23556, 5281, 84720, 54872, 81490, 254531, 10162, 129642, 154141, 10390, 151056, 100137049, 123745, 255189, 283748, 5321, 8605, 8681, 8398, 11145, 122618, 23646 | PEMT, PEAMT, PEMPT, PEMT2, PLMT, PNMT, SELENOI, EPT1, SELI, SEPI, SPG81, PLA2G2D, PLA2IID, SPLASH, sPLA2-IID, sPLA2S, PLA2G2E, GIIE\_sPLA2, sPLA2-IIE, PLA2G2C, UBXN10-AS1, PLA2G3, GIII-SPLA2, SPLA2III, sPLA2-III, PLA2G1B, PLA2, PLA2A, PPLA2, PLA2G2A, MOM1, PLA2, PLA2B, PLA2L, PLA2S, PLAS1, sPLA2, PLA2G5, FRFB, GV-PLA2, PLA2-10, hVPLA(2), PLA2G2F, GIIFsPLA2, sPLA2-IIF, PLA2G12A, GXII, PLA2G12, ROSSY, PLA2G10, GXPLA2, GXSPLA2, SPLA2, sPLA2-X, PLA2G12B, FKSG71, GXIIB, GXIIIsPLA2, PLA2G13, sPLA2-GXIIB, PLD1, CVDD, PLD2, PLD1C, PISD, DJ858B16, LIBF, PSD, PSDC, PSSC, dJ858B16.2, PIGN, MCAHS, MCAHS1, MCD4, MDC4, PIG-N, PIGF, PIGO, HPMRS2, PIGG, GPI7, LAS21, MRT53, PRO4405, RLGS1930, PTDSS2, PSS2, LPCAT4, AGPAT7, AYTL3, LPAAT-eta, LPEAT2, LPCAT3, C3F, LPCAT, LPLAT\_5, LPSAT, MBOAT5, OACT5, nessy, MBOAT2, LPAAT, LPCAT4, LPEAT, LPLAT\_2, OACT2, MBOAT1, LPEAT1, LPLAT, LPLAT\_1, LPSAT, OACT1, dJ434O11.1, CEPT1, PLB1, PLB, PLB/LIP, PLA2G4B, HsT16992, cPLA2-beta, PLA2G4E, PLA2G4F, PLA2G4FZ, PLA2G4D, cPLA2delta, PLA2G4A, GURDP, PLA2G4, cPLA2, cPLA2-alpha, PLA2G4C, CPLA2-gamma, JMJD7-PLA2G4B, HsT16992, cPLA2-beta, PLA2G6, CaI-PLA2, GVI, INAD1, IPLA2-VIA, NBIA2, NBIA2A, NBIA2B, PARK14, PLA2, PNPLA9, iPLA2, iPLA2beta, PLAAT3, AdPLA, H-REV107, H-REV107-1, HRASLS3, HREV107, HREV107-1, HREV107-3, HRSL3, PLA2G16, PLAAT-3, PLD4, C14orf175, PLD3, AD19, HU-K4, HUK4, SCA46 | (RefSeq) phosphatidylethanolamine N-methyltransferase, (RefSeq) selenoprotein I, (RefSeq) phospholipase A2 group IID, (RefSeq) phospholipase A2 group IIE, (RefSeq) phospholipase A2 group IIC, (RefSeq) phospholipase A2 group III, (RefSeq) phospholipase A2 group IB, (RefSeq) phospholipase A2 group IIA, (RefSeq) phospholipase A2 group V, (RefSeq) phospholipase A2 group IIF, (RefSeq) phospholipase A2 group XIIA, (RefSeq) phospholipase A2 group X, (RefSeq) phospholipase A2 group XIIB, (RefSeq) phospholipase D1, (RefSeq) phospholipase D2, (RefSeq) phosphatidylserine decarboxylase, (RefSeq) phosphatidylinositol glycan anchor biosynthesis class N, (RefSeq) phosphatidylinositol glycan anchor biosynthesis class F, (RefSeq) phosphatidylinositol glycan anchor biosynthesis class O, (RefSeq) phosphatidylinositol glycan anchor biosynthesis class G, (RefSeq) phosphatidylserine synthase 2, (RefSeq) lysophosphatidylcholine acyltransferase 4, (RefSeq) lysophosphatidylcholine acyltransferase 3, (RefSeq) membrane bound O-acyltransferase domain containing 2, (RefSeq) membrane bound O-acyltransferase domain containing 1, (RefSeq) choline/ethanolamine phosphotransferase 1, (RefSeq) phospholipase B1, (RefSeq) phospholipase A2 group IVB, (RefSeq) phospholipase A2 group IVE, (RefSeq) phospholipase A2 group IVF, (RefSeq) phospholipase A2 group IVD, (RefSeq) phospholipase A2 group IVA, (RefSeq) phospholipase A2 group IVC, (RefSeq) JMJD7-PLA2G4B readthrough, (RefSeq) phospholipase A2 group VI, (RefSeq) phospholipase A and acyltransferase 3, (RefSeq) phospholipase D family member 4, (RefSeq) phospholipase D family member 3 | Homo sapiens (human) | c(hsa00564 = “Glycerophospholipid metabolism”, hsa01100 = “Metabolic pathways”), c(hsa00440 = “Phosphonate and phosphinate metabolism”, hsa00564 = “Glycerophospholipid metabolism”, hsa00565 = “Ether lipid metabolism”, hsa01100 = “Metabolic pathways”), c(hsa00564 = “Glycerophospholipid metabolism”, hsa00565 = “Ether lipid metabolism”, hsa00590 = “Arachidonic acid metabolism”, hsa00591 = “Linoleic acid metabolism”, hsa00592 = “alpha-Linolenic acid metabolism”, hsa01100 = “Metabolic pathways”, hsa04014 = “Ras signaling pathway”, hsa04270 = “Vascular smooth muscle contraction”, hsa04972 = “Pancreatic secretion”, hsa04975 = “Fat digestion and absorption”), c(hsa00564 = “Glycerophospholipid metabolism”, hsa00565 = “Ether lipid metabolism”, hsa01100 = “Metabolic pathways”, hsa04014 = “Ras signaling pathway”, hsa04024 = “cAMP signaling pathway”, hsa04071 = “Sphingolipid signaling pathway”, hsa04072 = “Phospholipase D signaling pathway”, hsa04144 = “Endocytosis”, hsa04666 = “Fc gamma R-mediated phagocytosis”, hsa04724 = “Glutamatergic synapse”, hsa04912 = “GnRH signaling pathway”, hsa04928 = “Parathyroid hormone synthesis, secretion and action”, hsa05200 = “Pathways in cancer”, |  |  |  |  |  |  |  |  |  |  |  |
| hsa05212 = "P | ancreatic cancer“, hsa05231 =”Choline metabolism in cancer“), c(hsa00563 =”Glycosylphosphatidylinositol (GPI)-anchor biosynthesis“, hsa01100 =”Metabol | ic pathways“), Glycosylphosphatidylinositol (GPI)-anchor biosynthesis, c(hsa00564 =”Glycerophospholipid metabolism“, hsa00565 =”Ether lipid metabolism“, hsa01100 =”Metabolic pathways“), c(hsa00564 =”Glycerophospholipid metabolism“, hsa04216 =”Ferroptosis“), c(hsa00561 =”Glycerolipid metabolism“, hsa00564 =”Glycerophospholipid metabolism“, hsa01100 =”Metabolic pathways“), c(hsa00564 =”Glycerophospholipid metabolism“, hsa00565 =”Ether lipid metabolism“, hsa00590 =”Arachidonic acid metabolism“, hsa00591 =”Linoleic acid metabolism“, hsa00592 =”alpha-Linolenic acid metabolism“, hsa01100 =”Metabolic pathways“, hsa04977 =”Vitamin digestion and absorption“), c(hsa00564 =”Glycerophospholipid metabolism“, hsa00565 =”Ether lipid metabolism“, hsa00590 =”Arachidonic acid metabolism“, hsa00591 =”Linoleic acid metabolism“, hsa00592 =”alpha-Linolenic acid metabolism“, hsa01100 =”Metabolic pathways“, hsa04010 =”MAPK signaling pathway“, hsa04014 =”Ras signaling pathway“, hsa04072 =”Phospholipase D | signaling pathway“, hsa04217 =”Necroptosis“, hsa04270 =”Vascular smooth muscle contraction“, hsa04370 =”VEGF signaling pathway“, hsa04611 =”Platelet activation", |  |  |  |  |  |  |  |  |  |  |  |  |  |  |  |
| hsa04664 = "F | c epsilon RI signaling pathway“, hsa04666 =”Fc gamma R-mediated phagocytosis“, hsa04724 =”Glutamatergic synapse“, hsa04726 =”Serotonergic synapse", hs | a04730 = “Long-term depression”, hsa04750 = “Inflammatory mediator regulation of TRP channels”, hsa04912 = “GnRH signaling pathway”, hsa04913 = “Ovarian steroidogenesis”, hsa04921 = “Oxytocin signaling pathway”, hsa05231 = “Choline metabolism in cancer”), c(hsa00564 = “Glycerophospholipid metabolism”, hsa00565 = “Ether lipid metabolism”, hsa00590 = “Arachidonic acid metabolism”, hsa00591 = “Linoleic acid metabolism”, hsa00592 = “alpha-Linolenic acid metabolism”, hsa01100 = “Metabolic pathways”, hsa04014 = “Ras signaling pathway”, hsa04270 = “Vascular smooth muscle contraction”, hsa04666 = “Fc gamma R-mediated phagocytosis”, hsa04750 = “Inflammatory mediator regulation of TRP channels”), c(hsa00564 = “Glycerophospholipid metabolism”, hsa00565 = “Ether lipid metabolism”, hsa00590 = “Arachidonic acid metabolism”, hsa00591 = “Linoleic acid metabolism”, hsa00592 = “alpha-Linolenic acid metabolism”, hsa01100 = “Metabolic pathways”, hsa04014 = “Ras signaling pathway”, hsa04923 = "Regulation of lipolysis in adi | pocytes“) c(”NCBI-GeneID: 10400“,”NCBI-ProteinID: NP\_009100“,”OMIM: 602391“,”HGNC: 8830“,”Ensembl: ENSG00000133027“,”Vega: OTTHUMG00000059290“,”Pharos: Q9UBM1(Tbio)“,”UniProt: Q9UBM1“), c(”NCBI-GeneID: 85465“,”NCBI-ProteinID: NP\_277040“,”OMIM: 607915“,”HGNC | : 29361“,”Ensembl: ENSG00000138018“,”Vega: OTTHUMG00000151931“,”Pharos: Q9C0D9(Tbio)“,”UniProt: Q9C0D9“), c(”NCBI-GeneID: 26279“,”NCBI-ProteinID: NP\_036532“,”OMIM: 605630“,”HGNC: 9033“,”Ensembl: ENSG00000117215“,”Vega: OTTHUMG00000002701“,”Pharos: Q9UNK4(Tchem)“,”UniProt: Q9UNK4“), c(”NCBI-GeneID: 30814“,”NCBI-ProteinID: NP\_055404“,”OMIM: 618320“,”HGNC: 13414“,”Ensembl: ENSG00000188784“,”Vega: OTTHUMG00000002702“,”Pharos: Q9NZK7(Tchem)“,”UniProt: Q9NZK7“), c(”NCBI-GeneID: 391013“,”NCBI-ProteinID: NP\_001303651“,”HGNC: 9032“,”Ensembl: ENSG00000187980 ENSG00000225986“,”Vega: OTTHUMG00000002705“), c(”NCBI-GeneID: 50487“,”NCBI-ProteinID: NP\_056530“,”OMIM: 611651“,”HGNC: 17934“,”Ensembl: ENSG00000100078“,”Vega: OTTHUMG00000151255“,”Pharos: Q9NZ20(Tbio)“,”UniProt: Q9NZ20“), c(”NCBI-GeneID: 5319“,”NCBI-ProteinID: NP\_000919“,”OMIM: 172410“,”HGNC: 9030“,”Ensembl: ENSG00000170890“,”Vega: OTTHUMG00000169343“,”Pharos: P04054(Tchem)“,”UniProt: P04054“), c(”NCBI-GeneID: 5320“,”NCBI-ProteinID: NP\_000291“,”OMIM: 172411“,”HGNC: 9031“,”Ensembl: ENSG00000188257“,”Vega: OTTHUMG00000002699“,”Pharos: P14555(Tchem)“,”UniProt: P14555 A0A024RA96“), c(”NCBI-GeneID: 5322“,”NCBI-ProteinID: NP\_000920“,”OMIM: 601192“,”HGNC: 9038“,”Ensembl: ENSG00000127472“,”Vega: OTTHUMG00000002698“,”Pharos: P39877(Tchem)“,”UniProt: P39877"), c | (“NCBI-GeneID: 64600”, “NCBI-ProteinID: NP\_073730”, “OMIM: 616793”, “HGNC: 30040”, “Ensembl: ENSG00000158786”, “Vega: OTTHUMG00000002704”, “Pharos: Q9BZM2(Tchem)”, “UniProt: Q9BZM2”), c(“NCBI-GeneID: 81579”, “NCBI-ProteinID: NP\_110448”, “OMIM: 611652”, “HGNC: 18554”, “Ensembl: ENSG00000123739”, “Vega: OTTHUMG00000131915”, “Pharos: Q9BZM1(Tbio)”, “UniProt: Q9BZM1 Q542Y6”), c(“NCBI-GeneID: 8399”, “NCBI-ProteinID: NP\_003552”, “OMIM: 603603”, “HGNC: 9029”, “Ensembl: ENSG00000069764”, “Vega: OTTHUMG00000048069”, “Pharos: O15496(Tchem)”, “UniProt: O15496”), c(“NCBI-GeneID: 84647”, “NCBI-ProteinID: NP\_115951”, “OMIM: 611653”, “HGNC: 18555”, “Ensembl: ENSG00000138308”, “Vega: OTTHUMG00000018446”, “Pharos: Q9BX93(Tdark)”, “UniProt: Q9BX93”), c(“NCBI-GeneID: 5337”, “NCBI-ProteinID: NP\_002653”, “OMIM: 602382”, “HGNC: 9067”, “Ensembl: ENSG00000075651”, “Vega: OTTHUMG00000156947”, “Pharos: Q13393(Tchem)”, “UniProt: Q13393 Q59EA4”), c(“NCBI-GeneID: 5338”, “NCBI-ProteinID: NP\_002654”, “OMIM: 602384”, “HGNC: 9068”, “Ensembl: ENSG00000129219”, “Vega: OTTHUMG00000090779”, “Pharos: O14939(Tchem)”, “UniProt: O14939”), c(“NCBI-GeneID: 23761”, “NCBI-ProteinID: NP\_001313340”, “OMIM: 612770”, “HGNC: 8999”, “Ensembl: ENSG00000241878”, “Vega: OTTHUMG00000030252”, “Pharos: Q9UG56(Tbio)”, “UniProt: Q9UG56”), c(“NCBI-GeneID: 23556”, “NCBI-ProteinID: NP\_036459”, “OMIM: 606097”, “HGNC: 8967”, “Ensembl: ENSG00000197563”, “Vega: OTTHUMG00000180098”, “Pharos: O95427(Tbio)”, “UniProt: O95427 A0A024R2C3”), c(“NCBI-GeneID: 5281”, “NCBI-ProteinID: NP\_002634”, “OMIM: 600153”, “HGNC: 8962”, “Ensembl: ENSG00000151665”, "Vega: OTTHUMG00 | 000128816“,”Pharos: Q0 | 7326(Tbio)“,”UniProt: Q07326 Q6IB04“), c(”NCBI-GeneID: 84720“,”NCBI-ProteinID: NP\_116023“,”OMIM: 614730“,”HGNC: 23215“,”Ensembl: ENSG00000165282“,”Vega: OTTHUMG00000019854“,”Pharos: Q8TEQ8(Tbio)“,”UniProt: Q8TEQ8“), c(”NCBI-GeneID: 54872“,”NCBI-ProteinID: NP\_001120650“,”OMIM: 616918“,”HGNC: 25985“,”Ensembl: ENSG00000174227“,”Vega: OTTHUMG00000112457“,”Pharos: Q5H8A4(Tbio)“,”UniProt: Q5H8A4“), c(”NCBI-GeneID: 81490“,”NCBI-ProteinID: NP\_110410“,”OMIM: 612793“,”HGNC: 15463“,”Ensembl: ENSG00000174915“,”Vega: OTTHUMG00000119087“,”Pharos: Q9BVG9(Tbio)“,”UniProt: Q9BVG9 A0A024RC97“), c(”NCBI-GeneID: 254531“,”NCBI-ProteinID: NP\_705841“,”OMIM: 612039“,”HGNC: 30059“,”Ensembl: ENSG00000176454“,”Vega: OTTHUMG00000172349“,”Pharos: Q643R3(Tdark)“,”UniProt: Q643R3“), c(”NCBI-GeneID: 10162“,”NCBI-ProteinID: NP\_005759“,”OMIM: 611950“,”HGNC: 30244“,”Ensembl: ENSG00000111684“,”Vega: OTTHUMG00000168970“,”Pharos: Q6P1A2(Tbio)“,”UniProt: Q6P1A2“), c(”NCBI-GeneID: 129642“,”NCBI-ProteinID: NP\_620154“,”OMIM: 611949“,”HGNC: 25193“,”Ensembl: ENSG00000143797“,”Vega: OTTHUMG00000090363“,”Pharos: Q6ZWT7(Tdark)“,”UniProt: Q6ZWT7“), c(”NCBI-GeneID: 154141“,”NCBI-ProteinID: NP\_001073949“,”OMIM: 611732“,”HGNC: 21579“,”Ensembl: ENSG00000172197“,”Vega: OTTHUMG00000014334“,”Pharos: Q6ZNC8(Tdark)“,”UniProt: Q6ZNC8“), c(”NCBI-GeneID: 10390“,”NCBI-ProteinID: NP\_001007795“,”OMIM: 616751“,”HGNC: 24289“,”Ensembl: ENSG00000134255“,”Vega: OTTHUMG00000012357“,”Pharos: Q9Y6K0(Tbio)“,”UniProt: Q9Y6K0 A1PL14“), c(”NCBI-GeneID: 151056“,”NCBI-ProteinID: NP\_694566“,”OMIM: 610179“,”HGNC: 30041“,”Ensembl: ENSG00000163803“,”Vega: OTTHUMG00000152014“,”Pharos: Q6P1J6(Tdark)“,”UniProt: Q6P1J6 B2RWP8“), c(”NCBI-GeneID: 100137049“,”NCBI-ProteinID: NP\_001108105“,”OMIM: 606088“,”HGNC: 9036“,”Ensembl: ENSG00000243708“,”Vega: OTTHUMG00000156809“,”UniProt: P0C869“), c(”NCBI-GeneID: 123745“,”NCBI-ProteinID: NP\_001193599“,”HGNC: 24791“,”Ensembl: ENSG00000188089“,”Vega: OTTHUMG00000130371“,”Pharos: Q3MJ16(Tdark)“,”UniProt: Q3MJ16“), c(”NCBI-GeneID: 255189“,”NCBI-ProteinID: NP\_998765“,”HGNC: 27396“,”Ensembl: ENSG00000168907“,”Vega: OTTHUMG00000172782“,”Pharos: Q68DD2(Tdark)“,”UniProt: Q68DD2 A5PKZ7“), c(”NCBI-GeneID: 283748“,”NCBI-ProteinID: NP\_828848“,”OMIM: 612864“,”HGNC: 30038“,”Ensembl: ENSG00000159337“,”Vega: OTTHUMG00000172587“,”Pharos: Q86XP0(Tdark)“,”UniProt: Q86XP0“), c(”NCBI-GeneID: 5321“,”NCBI-ProteinID: NP\_077734“,”OMIM: 600522“,”HGNC: 9035“,”Ensembl: ENSG00000116711“,”Vega: OTTHUMG00000035512“,”Pharos: P47712(Tchem)“,”UniProt: P47712“), c(”NCBI-GeneID: 8605“,”NCBI-ProteinID: NP\_003697“,”OMIM: 603602“,”HGNC: 9037“,”Ensembl: ENSG00000105499“,”Vega: OTTHUMG00000183185“,”Pharos: Q9UP65(Tchem)“,”UniProt: Q9UP65 A0A024QZH0“), c(”NCBI-GeneID: 8681“,”NCBI-ProteinID: NP\_001185517“,”HGNC: 34449“,”Ensembl: ENSG00000168970“,”Vega: OTTHUMG00000044442“,”Pharos: P0C869(Tchem)“,”UniProt: P0C869“), c(”NCBI-GeneID: 8398“,”NCBI-ProteinID: NP\_003551“,”OMIM: 603604“,”HGNC: 9039“,”Ensembl: ENSG00000184381“,”Vega: OTTHUMG00000151246“,”Pharos: O60733(Tchem)“,”UniProt: O60733“), c(”NCBI-GeneID: 11145“,”NCBI-ProteinID: NP\_001121675“,”OMIM: 613867“,”HGNC: 17825“,”Ensembl: ENSG00000176485“,”Vega: OTTHUMG00000167852“,”Pharos: P53816(Tbio)“,”UniProt: P53816 A0A024R561“), c(”NCBI-GeneID: 122618“,”NCBI-ProteinID: NP\_620145“,”OMIM: 618488“,”HGNC: 23792“,”Ensembl: ENSG00000166428“,”Vega: OTTHUMG00000144167“,”Pharos: Q96BZ4(Tbio)“,”UniProt: Q96BZ4 B4DI07 B4DJQ6“), c(”NCBI-GeneID: 23646“,”NCBI-ProteinID: NP\_001026866“,”OMIM: 615698“,”HGNC: 17158“,”Ensembl: ENSG00000105223“,”Vega: OTTHUMG00000152736“,”Pharos: Q8IV08(Tbio)“,”U | niProt: Q8IV08 A0A024R0Q4“) Pfam: PEMT Herpes\_UL74, Pfam: CDP-OH\_P\_transf, Pfam: Phospholip\_A2\_1 Parvo\_coat\_N DUF5460, Pfam: Phospholip\_A2\_1, Pfam: Phospholip\_A2\_2 Chromadorea\_ALT, Pfam: Phospholip\_A2\_1 Parvo\_coat\_N Phospholip\_A2\_2 DUF1644, Pfam: Phospholip\_A2\_1 Phospholip\_A2\_2, Pfam: Phospholip\_A2\_1 Parvo\_coat\_N, Pfam: PLA2G12, Pfam: PLDc\_2 PX PLDc PH DUF3785, Pfam: PX PLDc\_2 PLDc PH PH\_2, Pfam: PS\_Dcarbxylase, Pfam: PigN Phosphodiest Sulfatase Metalloenzyme, Pfam: PIG-F, Pfam: Phosphodiest Metalloenzyme Sulfatase DUF1501, Pfam: Phosphodiest Metalloenzyme, Pfam: PSS TctB, Pfam: Acyltransferase JmjN, Pfam: MBOAT DUF5659, Pfam: MBOAT, Pfam: Lipase\_GDSL Lipase\_GDSL\_2, Pfam: PLA2\_B cPLA2\_C2 C2, Pfam: cPLA2\_C2 PLA2\_B C2, Pfam: PLA2\_B C2, Pfam: PLA2\_B, Pfam: Cupin\_8 PLA2\_B cPLA2\_C2 C2, Pfam: Ank\_2 Ank\_4 Ank\_5 Ank Ank\_3 Patatin, Pfam: LRAT Calici\_PP\_N Peptidase\_C97 Churchill, Pfam: PLDc\_3 PLDc\_2 PLDc R02056, R02057, R02053, R02051, R02055, R12351, R05923, R08107, R05924, R07376, R04480, R02054 C00350 PE S-adenosyl-L-methionine:phosphatidylethanolamine N-methyltransferase, CDPethanolamine:1,2-diacylglycerol ethanolaminephosphotransferase, phosphatidylethanolamine 2-acylhydrolase, phosphatidylethanolamine phosphatidohydrolase, Phsophatidyl-L-serine carboxy-lyase, NULL, L-1-phosphatidylethanolamine:L-serine phosphatidyltransferase, Acyl-CoA:1-acyl-sn-glycero-3-phosphoethanolamine O-acyltransferase, phosphatidylethanolamine 1-acylhydrolase c(”RC00003 C00019\_C00021“,”RC00060 C00350\_C01241“), c(”RC00002 C00055\_C00570“,”RC00017 C00350\_C00641“), c(”RC00037 C00350\_C04438“,”RC00094 C00162\_C00350“), c(”RC00017 C00189\_C00350“,”RC00425 C00350\_C00416“), RC00299 C00350\_C02737, NULL, RC00017 C00350\_C00641, c(”RC00017 C00065\_C02737 C00189\_C00350“,”RC02950 C00350\_C02737“), c(”RC00004 C00010\_C00040“,”RC00037 C00350\_C04438“), c(”RC00020 C00162\_C00350“,”RC00041 C00350\_C05973") C00019 + C00350 <=> C00021 + C01241, C00570 + C00641 <=> C00055 + C00350, C00350 + C00001 <=> C04438 + C00162, C00350 + C00001 <=> C00189 + C00416, C02737 <=> C00350 + C00011, G13128 + C00350 <=> G00148 + C00641, C00350 + G00140 <=> C00641 + G00141, C00350 + G00149 <=> C00641 + G13044, C00350 + G00141 <=> C00641 + G00151, C00350 + C00065 <=> C02737 + C00189, C00350 + C00010 <=> C04438 + C00040, C00350 + C00001 <=> C05973 + C00162 S-Adenosyl-L-methionine + Phosphatidylethanolamine <=> S-Adenosyl-L-homocysteine + Phosphatidyl-N-methylethanolamine, CDP-ethanolamine + 1,2-Diacyl-sn-glycerol <=> CMP + Phosphatidylethanolamine, Phosphatidylethanolamine + H2O <=> 1-Acyl-sn-glycero-3-phosphoethanolamine + Fatty acid, Phosphatidylethanolamine + H2O <=> Ethanolamine + Phosphatidate, Phosphatidylserine <=> Phosphatidylethanolamine + CO2, G13128 + Phosphatidylethanolamine <=> G00148 + 1,2-Diacyl-sn-glycerol, Phosphatidylethanolamine + G00140 <=> 1,2-Diacyl-sn-glycerol + G00141, Phosphatidylethanolamine + G00149 <=> 1,2-Diacyl-sn-glycerol + G13044, Phosphatidylethanolamine + G00141 <=> 1,2-Diacyl-sn-glycerol + G00151, Phosphatidylethanolamine + L-Serine <=> Phosphatidylserine + Ethanolamine, Phosphatidylethanolamine + CoA <=> 1-Acyl-sn-glycero-3-phosphoethanolamine + Acyl-CoA, Phosphatidylethanolamine + H2O <=> 2-Acyl-sn-glycero-3-phosphoethanolamine + Fatty acid 2.1.1.17, 2.7.8.1, 3.1.1.4, 3.1.4.4, 4.1.1.65, 2.7.-.-, NULL, 2.7.8.29, 2.3.1.23, 3.1.1.32 Substrate, Product |  |  |  |  |  |  |  |  |  |  |
| PE(36:3) | K00551, K00993, K01047, K01115, K01613, K05285, K05287, K05288, K05310, K08730, K13512, K13515, K13517, K13644, K14621, K16342, K16343, K16817, K16860 | phosphatidylethanolamine/phosphatidyl-N-methylethanolamine N-methyltransferase [EC:2.1.1.17 2.1.1.71], ethanolaminephosphotransferase [EC:2.7.8.1], secretory phospholipase A2 [EC:3.1.1.4], phospholipase D1/2 [EC:3.1.4.4], phosphatidylserine decarboxylase [EC:4.1.1.65], GPI ethanolamine phosphate transferase 1 [EC:2.7.-.-], GPI ethanolamine phosphate transferase 2/3 subunit F, GPI ethanolamine phosphate transferase 3 subunit O [EC:2.7.-.-], ethanolamine phosphate transferase 2 subunit G [EC:2.7.-.-], phosphatidylserine synthase 2 [EC:2.7.8.29], lysophospholipid acyltransferase [EC:2.3.1.23 2.3.1.-], lysophospholipid acyltransferase 5 [EC:2.3.1.23 2.3.1.-], lysophospholipid acyltransferase 1/2 [EC:2.3.1.51 2.3.1.-], choline/ethanolamine phosphotransferase [EC:2.7.8.1 2.7.8.2], phospholipase B1, membrane-associated [EC:3.1.1.4 3.1.1.5], cytosolic phospholipase A2 [EC:3.1.1.4], calcium-independent phospholipase A2 [EC:3.1.1.4], HRAS-like suppressor 3 [EC:3.1.1.32 3.1.1.4], phospholipase D3/4 [EC:3.1.4.4] | 10400, 85465, 26279, 30814, 391013, 50487, 5319, 5320, 5322, 64600, 81579, 8399, 84647, 5337, 5338, 23761, 23556, 5281, 84720, 54872, 81490, 254531, 10162, 129642, 154141, 10390, 151056, 100137049, 123745, 255189, 283748, 5321, 8605, 8681, 8398, 11145, 122618, 23646 | PEMT, PEAMT, PEMPT, PEMT2, PLMT, PNMT, SELENOI, EPT1, SELI, SEPI, SPG81, PLA2G2D, PLA2IID, SPLASH, sPLA2-IID, sPLA2S, PLA2G2E, GIIE\_sPLA2, sPLA2-IIE, PLA2G2C, UBXN10-AS1, PLA2G3, GIII-SPLA2, SPLA2III, sPLA2-III, PLA2G1B, PLA2, PLA2A, PPLA2, PLA2G2A, MOM1, PLA2, PLA2B, PLA2L, PLA2S, PLAS1, sPLA2, PLA2G5, FRFB, GV-PLA2, PLA2-10, hVPLA(2), PLA2G2F, GIIFsPLA2, sPLA2-IIF, PLA2G12A, GXII, PLA2G12, ROSSY, PLA2G10, GXPLA2, GXSPLA2, SPLA2, sPLA2-X, PLA2G12B, FKSG71, GXIIB, GXIIIsPLA2, PLA2G13, sPLA2-GXIIB, PLD1, CVDD, PLD2, PLD1C, PISD, DJ858B16, LIBF, PSD, PSDC, PSSC, dJ858B16.2, PIGN, MCAHS, MCAHS1, MCD4, MDC4, PIG-N, PIGF, PIGO, HPMRS2, PIGG, GPI7, LAS21, MRT53, PRO4405, RLGS1930, PTDSS2, PSS2, LPCAT4, AGPAT7, AYTL3, LPAAT-eta, LPEAT2, LPCAT3, C3F, LPCAT, LPLAT\_5, LPSAT, MBOAT5, OACT5, nessy, MBOAT2, LPAAT, LPCAT4, LPEAT, LPLAT\_2, OACT2, MBOAT1, LPEAT1, LPLAT, LPLAT\_1, LPSAT, OACT1, dJ434O11.1, CEPT1, PLB1, PLB, PLB/LIP, PLA2G4B, HsT16992, cPLA2-beta, PLA2G4E, PLA2G4F, PLA2G4FZ, PLA2G4D, cPLA2delta, PLA2G4A, GURDP, PLA2G4, cPLA2, cPLA2-alpha, PLA2G4C, CPLA2-gamma, JMJD7-PLA2G4B, HsT16992, cPLA2-beta, PLA2G6, CaI-PLA2, GVI, INAD1, IPLA2-VIA, NBIA2, NBIA2A, NBIA2B, PARK14, PLA2, PNPLA9, iPLA2, iPLA2beta, PLAAT3, AdPLA, H-REV107, H-REV107-1, HRASLS3, HREV107, HREV107-1, HREV107-3, HRSL3, PLA2G16, PLAAT-3, PLD4, C14orf175, PLD3, AD19, HU-K4, HUK4, SCA46 | (RefSeq) phosphatidylethanolamine N-methyltransferase, (RefSeq) selenoprotein I, (RefSeq) phospholipase A2 group IID, (RefSeq) phospholipase A2 group IIE, (RefSeq) phospholipase A2 group IIC, (RefSeq) phospholipase A2 group III, (RefSeq) phospholipase A2 group IB, (RefSeq) phospholipase A2 group IIA, (RefSeq) phospholipase A2 group V, (RefSeq) phospholipase A2 group IIF, (RefSeq) phospholipase A2 group XIIA, (RefSeq) phospholipase A2 group X, (RefSeq) phospholipase A2 group XIIB, (RefSeq) phospholipase D1, (RefSeq) phospholipase D2, (RefSeq) phosphatidylserine decarboxylase, (RefSeq) phosphatidylinositol glycan anchor biosynthesis class N, (RefSeq) phosphatidylinositol glycan anchor biosynthesis class F, (RefSeq) phosphatidylinositol glycan anchor biosynthesis class O, (RefSeq) phosphatidylinositol glycan anchor biosynthesis class G, (RefSeq) phosphatidylserine synthase 2, (RefSeq) lysophosphatidylcholine acyltransferase 4, (RefSeq) lysophosphatidylcholine acyltransferase 3, (RefSeq) membrane bound O-acyltransferase domain containing 2, (RefSeq) membrane bound O-acyltransferase domain containing 1, (RefSeq) choline/ethanolamine phosphotransferase 1, (RefSeq) phospholipase B1, (RefSeq) phospholipase A2 group IVB, (RefSeq) phospholipase A2 group IVE, (RefSeq) phospholipase A2 group IVF, (RefSeq) phospholipase A2 group IVD, (RefSeq) phospholipase A2 group IVA, (RefSeq) phospholipase A2 group IVC, (RefSeq) JMJD7-PLA2G4B readthrough, (RefSeq) phospholipase A2 group VI, (RefSeq) phospholipase A and acyltransferase 3, (RefSeq) phospholipase D family member 4, (RefSeq) phospholipase D family member 3 | Homo sapiens (human) | c(hsa00564 = “Glycerophospholipid metabolism”, hsa01100 = “Metabolic pathways”), c(hsa00440 = “Phosphonate and phosphinate metabolism”, hsa00564 = “Glycerophospholipid metabolism”, hsa00565 = “Ether lipid metabolism”, hsa01100 = “Metabolic pathways”), c(hsa00564 = “Glycerophospholipid metabolism”, hsa00565 = “Ether lipid metabolism”, hsa00590 = “Arachidonic acid metabolism”, hsa00591 = “Linoleic acid metabolism”, hsa00592 = “alpha-Linolenic acid metabolism”, hsa01100 = “Metabolic pathways”, hsa04014 = “Ras signaling pathway”, hsa04270 = “Vascular smooth muscle contraction”, hsa04972 = “Pancreatic secretion”, hsa04975 = “Fat digestion and absorption”), c(hsa00564 = “Glycerophospholipid metabolism”, hsa00565 = “Ether lipid metabolism”, hsa01100 = “Metabolic pathways”, hsa04014 = “Ras signaling pathway”, hsa04024 = “cAMP signaling pathway”, hsa04071 = “Sphingolipid signaling pathway”, hsa04072 = “Phospholipase D signaling pathway”, hsa04144 = “Endocytosis”, hsa04666 = “Fc gamma R-mediated phagocytosis”, hsa04724 = “Glutamatergic synapse”, hsa04912 = “GnRH signaling pathway”, hsa04928 = “Parathyroid hormone synthesis, secretion and action”, hsa05200 = “Pathways in cancer”, |  |  |  |  |  |  |  |  |  |  |  |
| hsa05212 = "P | ancreatic cancer“, hsa05231 =”Choline metabolism in cancer“), c(hsa00563 =”Glycosylphosphatidylinositol (GPI)-anchor biosynthesis“, hsa01100 =”Metabol | ic pathways“), Glycosylphosphatidylinositol (GPI)-anchor biosynthesis, c(hsa00564 =”Glycerophospholipid metabolism“, hsa00565 =”Ether lipid metabolism“, hsa01100 =”Metabolic pathways“), c(hsa00564 =”Glycerophospholipid metabolism“, hsa04216 =”Ferroptosis“), c(hsa00561 =”Glycerolipid metabolism“, hsa00564 =”Glycerophospholipid metabolism“, hsa01100 =”Metabolic pathways“), c(hsa00564 =”Glycerophospholipid metabolism“, hsa00565 =”Ether lipid metabolism“, hsa00590 =”Arachidonic acid metabolism“, hsa00591 =”Linoleic acid metabolism“, hsa00592 =”alpha-Linolenic acid metabolism“, hsa01100 =”Metabolic pathways“, hsa04977 =”Vitamin digestion and absorption“), c(hsa00564 =”Glycerophospholipid metabolism“, hsa00565 =”Ether lipid metabolism“, hsa00590 =”Arachidonic acid metabolism“, hsa00591 =”Linoleic acid metabolism“, hsa00592 =”alpha-Linolenic acid metabolism“, hsa01100 =”Metabolic pathways“, hsa04010 =”MAPK signaling pathway“, hsa04014 =”Ras signaling pathway“, hsa04072 =”Phospholipase D | signaling pathway“, hsa04217 =”Necroptosis“, hsa04270 =”Vascular smooth muscle contraction“, hsa04370 =”VEGF signaling pathway“, hsa04611 =”Platelet activation", |  |  |  |  |  |  |  |  |  |  |  |  |  |  |  |
| hsa04664 = "F | c epsilon RI signaling pathway“, hsa04666 =”Fc gamma R-mediated phagocytosis“, hsa04724 =”Glutamatergic synapse“, hsa04726 =”Serotonergic synapse", hs | a04730 = “Long-term depression”, hsa04750 = “Inflammatory mediator regulation of TRP channels”, hsa04912 = “GnRH signaling pathway”, hsa04913 = “Ovarian steroidogenesis”, hsa04921 = “Oxytocin signaling pathway”, hsa05231 = “Choline metabolism in cancer”), c(hsa00564 = “Glycerophospholipid metabolism”, hsa00565 = “Ether lipid metabolism”, hsa00590 = “Arachidonic acid metabolism”, hsa00591 = “Linoleic acid metabolism”, hsa00592 = “alpha-Linolenic acid metabolism”, hsa01100 = “Metabolic pathways”, hsa04014 = “Ras signaling pathway”, hsa04270 = “Vascular smooth muscle contraction”, hsa04666 = “Fc gamma R-mediated phagocytosis”, hsa04750 = “Inflammatory mediator regulation of TRP channels”), c(hsa00564 = “Glycerophospholipid metabolism”, hsa00565 = “Ether lipid metabolism”, hsa00590 = “Arachidonic acid metabolism”, hsa00591 = “Linoleic acid metabolism”, hsa00592 = “alpha-Linolenic acid metabolism”, hsa01100 = “Metabolic pathways”, hsa04014 = “Ras signaling pathway”, hsa04923 = "Regulation of lipolysis in adi | pocytes“) c(”NCBI-GeneID: 10400“,”NCBI-ProteinID: NP\_009100“,”OMIM: 602391“,”HGNC: 8830“,”Ensembl: ENSG00000133027“,”Vega: OTTHUMG00000059290“,”Pharos: Q9UBM1(Tbio)“,”UniProt: Q9UBM1“), c(”NCBI-GeneID: 85465“,”NCBI-ProteinID: NP\_277040“,”OMIM: 607915“,”HGNC | : 29361“,”Ensembl: ENSG00000138018“,”Vega: OTTHUMG00000151931“,”Pharos: Q9C0D9(Tbio)“,”UniProt: Q9C0D9“), c(”NCBI-GeneID: 26279“,”NCBI-ProteinID: NP\_036532“,”OMIM: 605630“,”HGNC: 9033“,”Ensembl: ENSG00000117215“,”Vega: OTTHUMG00000002701“,”Pharos: Q9UNK4(Tchem)“,”UniProt: Q9UNK4“), c(”NCBI-GeneID: 30814“,”NCBI-ProteinID: NP\_055404“,”OMIM: 618320“,”HGNC: 13414“,”Ensembl: ENSG00000188784“,”Vega: OTTHUMG00000002702“,”Pharos: Q9NZK7(Tchem)“,”UniProt: Q9NZK7“), c(”NCBI-GeneID: 391013“,”NCBI-ProteinID: NP\_001303651“,”HGNC: 9032“,”Ensembl: ENSG00000187980 ENSG00000225986“,”Vega: OTTHUMG00000002705“), c(”NCBI-GeneID: 50487“,”NCBI-ProteinID: NP\_056530“,”OMIM: 611651“,”HGNC: 17934“,”Ensembl: ENSG00000100078“,”Vega: OTTHUMG00000151255“,”Pharos: Q9NZ20(Tbio)“,”UniProt: Q9NZ20“), c(”NCBI-GeneID: 5319“,”NCBI-ProteinID: NP\_000919“,”OMIM: 172410“,”HGNC: 9030“,”Ensembl: ENSG00000170890“,”Vega: OTTHUMG00000169343“,”Pharos: P04054(Tchem)“,”UniProt: P04054“), c(”NCBI-GeneID: 5320“,”NCBI-ProteinID: NP\_000291“,”OMIM: 172411“,”HGNC: 9031“,”Ensembl: ENSG00000188257“,”Vega: OTTHUMG00000002699“,”Pharos: P14555(Tchem)“,”UniProt: P14555 A0A024RA96“), c(”NCBI-GeneID: 5322“,”NCBI-ProteinID: NP\_000920“,”OMIM: 601192“,”HGNC: 9038“,”Ensembl: ENSG00000127472“,”Vega: OTTHUMG00000002698“,”Pharos: P39877(Tchem)“,”UniProt: P39877"), c | (“NCBI-GeneID: 64600”, “NCBI-ProteinID: NP\_073730”, “OMIM: 616793”, “HGNC: 30040”, “Ensembl: ENSG00000158786”, “Vega: OTTHUMG00000002704”, “Pharos: Q9BZM2(Tchem)”, “UniProt: Q9BZM2”), c(“NCBI-GeneID: 81579”, “NCBI-ProteinID: NP\_110448”, “OMIM: 611652”, “HGNC: 18554”, “Ensembl: ENSG00000123739”, “Vega: OTTHUMG00000131915”, “Pharos: Q9BZM1(Tbio)”, “UniProt: Q9BZM1 Q542Y6”), c(“NCBI-GeneID: 8399”, “NCBI-ProteinID: NP\_003552”, “OMIM: 603603”, “HGNC: 9029”, “Ensembl: ENSG00000069764”, “Vega: OTTHUMG00000048069”, “Pharos: O15496(Tchem)”, “UniProt: O15496”), c(“NCBI-GeneID: 84647”, “NCBI-ProteinID: NP\_115951”, “OMIM: 611653”, “HGNC: 18555”, “Ensembl: ENSG00000138308”, “Vega: OTTHUMG00000018446”, “Pharos: Q9BX93(Tdark)”, “UniProt: Q9BX93”), c(“NCBI-GeneID: 5337”, “NCBI-ProteinID: NP\_002653”, “OMIM: 602382”, “HGNC: 9067”, “Ensembl: ENSG00000075651”, “Vega: OTTHUMG00000156947”, “Pharos: Q13393(Tchem)”, “UniProt: Q13393 Q59EA4”), c(“NCBI-GeneID: 5338”, “NCBI-ProteinID: NP\_002654”, “OMIM: 602384”, “HGNC: 9068”, “Ensembl: ENSG00000129219”, “Vega: OTTHUMG00000090779”, “Pharos: O14939(Tchem)”, “UniProt: O14939”), c(“NCBI-GeneID: 23761”, “NCBI-ProteinID: NP\_001313340”, “OMIM: 612770”, “HGNC: 8999”, “Ensembl: ENSG00000241878”, “Vega: OTTHUMG00000030252”, “Pharos: Q9UG56(Tbio)”, “UniProt: Q9UG56”), c(“NCBI-GeneID: 23556”, “NCBI-ProteinID: NP\_036459”, “OMIM: 606097”, “HGNC: 8967”, “Ensembl: ENSG00000197563”, “Vega: OTTHUMG00000180098”, “Pharos: O95427(Tbio)”, “UniProt: O95427 A0A024R2C3”), c(“NCBI-GeneID: 5281”, “NCBI-ProteinID: NP\_002634”, “OMIM: 600153”, “HGNC: 8962”, “Ensembl: ENSG00000151665”, "Vega: OTTHUMG00 | 000128816“,”Pharos: Q0 | 7326(Tbio)“,”UniProt: Q07326 Q6IB04“), c(”NCBI-GeneID: 84720“,”NCBI-ProteinID: NP\_116023“,”OMIM: 614730“,”HGNC: 23215“,”Ensembl: ENSG00000165282“,”Vega: OTTHUMG00000019854“,”Pharos: Q8TEQ8(Tbio)“,”UniProt: Q8TEQ8“), c(”NCBI-GeneID: 54872“,”NCBI-ProteinID: NP\_001120650“,”OMIM: 616918“,”HGNC: 25985“,”Ensembl: ENSG00000174227“,”Vega: OTTHUMG00000112457“,”Pharos: Q5H8A4(Tbio)“,”UniProt: Q5H8A4“), c(”NCBI-GeneID: 81490“,”NCBI-ProteinID: NP\_110410“,”OMIM: 612793“,”HGNC: 15463“,”Ensembl: ENSG00000174915“,”Vega: OTTHUMG00000119087“,”Pharos: Q9BVG9(Tbio)“,”UniProt: Q9BVG9 A0A024RC97“), c(”NCBI-GeneID: 254531“,”NCBI-ProteinID: NP\_705841“,”OMIM: 612039“,”HGNC: 30059“,”Ensembl: ENSG00000176454“,”Vega: OTTHUMG00000172349“,”Pharos: Q643R3(Tdark)“,”UniProt: Q643R3“), c(”NCBI-GeneID: 10162“,”NCBI-ProteinID: NP\_005759“,”OMIM: 611950“,”HGNC: 30244“,”Ensembl: ENSG00000111684“,”Vega: OTTHUMG00000168970“,”Pharos: Q6P1A2(Tbio)“,”UniProt: Q6P1A2“), c(”NCBI-GeneID: 129642“,”NCBI-ProteinID: NP\_620154“,”OMIM: 611949“,”HGNC: 25193“,”Ensembl: ENSG00000143797“,”Vega: OTTHUMG00000090363“,”Pharos: Q6ZWT7(Tdark)“,”UniProt: Q6ZWT7“), c(”NCBI-GeneID: 154141“,”NCBI-ProteinID: NP\_001073949“,”OMIM: 611732“,”HGNC: 21579“,”Ensembl: ENSG00000172197“,”Vega: OTTHUMG00000014334“,”Pharos: Q6ZNC8(Tdark)“,”UniProt: Q6ZNC8“), c(”NCBI-GeneID: 10390“,”NCBI-ProteinID: NP\_001007795“,”OMIM: 616751“,”HGNC: 24289“,”Ensembl: ENSG00000134255“,”Vega: OTTHUMG00000012357“,”Pharos: Q9Y6K0(Tbio)“,”UniProt: Q9Y6K0 A1PL14“), c(”NCBI-GeneID: 151056“,”NCBI-ProteinID: NP\_694566“,”OMIM: 610179“,”HGNC: 30041“,”Ensembl: ENSG00000163803“,”Vega: OTTHUMG00000152014“,”Pharos: Q6P1J6(Tdark)“,”UniProt: Q6P1J6 B2RWP8“), c(”NCBI-GeneID: 100137049“,”NCBI-ProteinID: NP\_001108105“,”OMIM: 606088“,”HGNC: 9036“,”Ensembl: ENSG00000243708“,”Vega: OTTHUMG00000156809“,”UniProt: P0C869“), c(”NCBI-GeneID: 123745“,”NCBI-ProteinID: NP\_001193599“,”HGNC: 24791“,”Ensembl: ENSG00000188089“,”Vega: OTTHUMG00000130371“,”Pharos: Q3MJ16(Tdark)“,”UniProt: Q3MJ16“), c(”NCBI-GeneID: 255189“,”NCBI-ProteinID: NP\_998765“,”HGNC: 27396“,”Ensembl: ENSG00000168907“,”Vega: OTTHUMG00000172782“,”Pharos: Q68DD2(Tdark)“,”UniProt: Q68DD2 A5PKZ7“), c(”NCBI-GeneID: 283748“,”NCBI-ProteinID: NP\_828848“,”OMIM: 612864“,”HGNC: 30038“,”Ensembl: ENSG00000159337“,”Vega: OTTHUMG00000172587“,”Pharos: Q86XP0(Tdark)“,”UniProt: Q86XP0“), c(”NCBI-GeneID: 5321“,”NCBI-ProteinID: NP\_077734“,”OMIM: 600522“,”HGNC: 9035“,”Ensembl: ENSG00000116711“,”Vega: OTTHUMG00000035512“,”Pharos: P47712(Tchem)“,”UniProt: P47712“), c(”NCBI-GeneID: 8605“,”NCBI-ProteinID: NP\_003697“,”OMIM: 603602“,”HGNC: 9037“,”Ensembl: ENSG00000105499“,”Vega: OTTHUMG00000183185“,”Pharos: Q9UP65(Tchem)“,”UniProt: Q9UP65 A0A024QZH0“), c(”NCBI-GeneID: 8681“,”NCBI-ProteinID: NP\_001185517“,”HGNC: 34449“,”Ensembl: ENSG00000168970“,”Vega: OTTHUMG00000044442“,”Pharos: P0C869(Tchem)“,”UniProt: P0C869“), c(”NCBI-GeneID: 8398“,”NCBI-ProteinID: NP\_003551“,”OMIM: 603604“,”HGNC: 9039“,”Ensembl: ENSG00000184381“,”Vega: OTTHUMG00000151246“,”Pharos: O60733(Tchem)“,”UniProt: O60733“), c(”NCBI-GeneID: 11145“,”NCBI-ProteinID: NP\_001121675“,”OMIM: 613867“,”HGNC: 17825“,”Ensembl: ENSG00000176485“,”Vega: OTTHUMG00000167852“,”Pharos: P53816(Tbio)“,”UniProt: P53816 A0A024R561“), c(”NCBI-GeneID: 122618“,”NCBI-ProteinID: NP\_620145“,”OMIM: 618488“,”HGNC: 23792“,”Ensembl: ENSG00000166428“,”Vega: OTTHUMG00000144167“,”Pharos: Q96BZ4(Tbio)“,”UniProt: Q96BZ4 B4DI07 B4DJQ6“), c(”NCBI-GeneID: 23646“,”NCBI-ProteinID: NP\_001026866“,”OMIM: 615698“,”HGNC: 17158“,”Ensembl: ENSG00000105223“,”Vega: OTTHUMG00000152736“,”Pharos: Q8IV08(Tbio)“,”U | niProt: Q8IV08 A0A024R0Q4“) Pfam: PEMT Herpes\_UL74, Pfam: CDP-OH\_P\_transf, Pfam: Phospholip\_A2\_1 Parvo\_coat\_N DUF5460, Pfam: Phospholip\_A2\_1, Pfam: Phospholip\_A2\_2 Chromadorea\_ALT, Pfam: Phospholip\_A2\_1 Parvo\_coat\_N Phospholip\_A2\_2 DUF1644, Pfam: Phospholip\_A2\_1 Phospholip\_A2\_2, Pfam: Phospholip\_A2\_1 Parvo\_coat\_N, Pfam: PLA2G12, Pfam: PLDc\_2 PX PLDc PH DUF3785, Pfam: PX PLDc\_2 PLDc PH PH\_2, Pfam: PS\_Dcarbxylase, Pfam: PigN Phosphodiest Sulfatase Metalloenzyme, Pfam: PIG-F, Pfam: Phosphodiest Metalloenzyme Sulfatase DUF1501, Pfam: Phosphodiest Metalloenzyme, Pfam: PSS TctB, Pfam: Acyltransferase JmjN, Pfam: MBOAT DUF5659, Pfam: MBOAT, Pfam: Lipase\_GDSL Lipase\_GDSL\_2, Pfam: PLA2\_B cPLA2\_C2 C2, Pfam: cPLA2\_C2 PLA2\_B C2, Pfam: PLA2\_B C2, Pfam: PLA2\_B, Pfam: Cupin\_8 PLA2\_B cPLA2\_C2 C2, Pfam: Ank\_2 Ank\_4 Ank\_5 Ank Ank\_3 Patatin, Pfam: LRAT Calici\_PP\_N Peptidase\_C97 Churchill, Pfam: PLDc\_3 PLDc\_2 PLDc R02056, R02057, R02053, R02051, R02055, R12351, R08107, R05924, R05923, R07376, R04480, R02054 C00350 PE S-adenosyl-L-methionine:phosphatidylethanolamine N-methyltransferase, CDPethanolamine:1,2-diacylglycerol ethanolaminephosphotransferase, phosphatidylethanolamine 2-acylhydrolase, phosphatidylethanolamine phosphatidohydrolase, Phsophatidyl-L-serine carboxy-lyase, NULL, L-1-phosphatidylethanolamine:L-serine phosphatidyltransferase, Acyl-CoA:1-acyl-sn-glycero-3-phosphoethanolamine O-acyltransferase, phosphatidylethanolamine 1-acylhydrolase c(”RC00003 C00019\_C00021“,”RC00060 C00350\_C01241“), c(”RC00002 C00055\_C00570“,”RC00017 C00350\_C00641“), c(”RC00037 C00350\_C04438“,”RC00094 C00162\_C00350“), c(”RC00017 C00189\_C00350“,”RC00425 C00350\_C00416“), RC00299 C00350\_C02737, NULL, RC00017 C00350\_C00641, c(”RC00017 C00065\_C02737 C00189\_C00350“,”RC02950 C00350\_C02737“), c(”RC00004 C00010\_C00040“,”RC00037 C00350\_C04438“), c(”RC00020 C00162\_C00350“,”RC00041 C00350\_C05973") C00019 + C00350 <=> C00021 + C01241, C00570 + C00641 <=> C00055 + C00350, C00350 + C00001 <=> C04438 + C00162, C00350 + C00001 <=> C00189 + C00416, C02737 <=> C00350 + C00011, G13128 + C00350 <=> G00148 + C00641, C00350 + G00149 <=> C00641 + G13044, C00350 + G00141 <=> C00641 + G00151, C00350 + G00140 <=> C00641 + G00141, C00350 + C00065 <=> C02737 + C00189, C00350 + C00010 <=> C04438 + C00040, C00350 + C00001 <=> C05973 + C00162 S-Adenosyl-L-methionine + Phosphatidylethanolamine <=> S-Adenosyl-L-homocysteine + Phosphatidyl-N-methylethanolamine, CDP-ethanolamine + 1,2-Diacyl-sn-glycerol <=> CMP + Phosphatidylethanolamine, Phosphatidylethanolamine + H2O <=> 1-Acyl-sn-glycero-3-phosphoethanolamine + Fatty acid, Phosphatidylethanolamine + H2O <=> Ethanolamine + Phosphatidate, Phosphatidylserine <=> Phosphatidylethanolamine + CO2, G13128 + Phosphatidylethanolamine <=> G00148 + 1,2-Diacyl-sn-glycerol, Phosphatidylethanolamine + G00149 <=> 1,2-Diacyl-sn-glycerol + G13044, Phosphatidylethanolamine + G00141 <=> 1,2-Diacyl-sn-glycerol + G00151, Phosphatidylethanolamine + G00140 <=> 1,2-Diacyl-sn-glycerol + G00141, Phosphatidylethanolamine + L-Serine <=> Phosphatidylserine + Ethanolamine, Phosphatidylethanolamine + CoA <=> 1-Acyl-sn-glycero-3-phosphoethanolamine + Acyl-CoA, Phosphatidylethanolamine + H2O <=> 2-Acyl-sn-glycero-3-phosphoethanolamine + Fatty acid 2.1.1.17, 2.7.8.1, 3.1.1.4, 3.1.4.4, 4.1.1.65, 2.7.-.-, NULL, 2.7.8.29, 2.3.1.23, 3.1.1.32 Substrate, Product |  |  |  |  |  |  |  |  |  |  |
| CE(22:6) | K00637, K01052, K12298 | sterol O-acyltransferase [EC:2.3.1.26], lysosomal acid lipase/cholesteryl ester hydrolase [EC:3.1.1.13], bile salt-stimulated lipase [EC:3.1.1.3 3.1.1.13] | 6646, 8435, 3988, 1056 | SOAT1, ACACT, ACAT, ACAT-1, ACAT1, SOAT, STAT, SOAT2, ACACT2, ACAT2, ARGP2, LIPA, CESD, LAL, CEL, BAL, BSDL, BSSL, CELL, CEase, FAP, FAPP, LIPA, MODY8 | (RefSeq) sterol O-acyltransferase 1, (RefSeq) sterol O-acyltransferase 2, (RefSeq) lipase A, lysosomal acid type, (RefSeq) carboxyl ester lipase | Homo sapiens (human) | c(hsa00100 = “Steroid biosynthesis”, hsa04979 = “Cholesterol metabolism”), c(hsa00100 = “Steroid biosynthesis”, hsa04142 = “Lysosome”, hsa04979 = “Cholesterol metabolism”), c(hsa00100 = “Steroid biosynthesis”, hsa00561 = “Glycerolipid metabolism”, hsa01100 = “Metabolic pathways”, hsa04972 = “Pancreatic secretion”, hsa04975 = “Fat digestion and absorption”) | c(“NCBI-GeneID: 6646”, “NCBI-ProteinID: NP\_003092”, “OMIM: 102642”, “HGNC: 11177”, “Ensembl: ENSG00000057252”, “Vega: OTTHUMG00000035253”, “Pharos: P35610(Tchem)”, “UniProt: P35610”), c(“NCBI-GeneID: 8435”, “NCBI-ProteinID: NP\_003569”, “OMIM: 601311”, “HGNC: 11178”, “Ensembl: ENSG00000167780”, “Vega: OTTHUMG00000169774”, “Pharos: O75908(Tchem)”, “UniProt: O75908”), c(“NCBI-GeneID: 3988”, “NCBI-ProteinID: NP\_000226”, “OMIM: 613497”, “HGNC: 6617”, “Ensembl: ENSG00000107798”, “Vega: OTTHUMG00000018716”, “Pharos: P38571(Tchem)”, “UniProt: P38571”), c(“NCBI-GeneID: 1056”, “NCBI-ProteinID: NP\_001798”, “OMIM: 114840”, “HGNC: 1848”, “Ensembl: ENSG00000170835”, “Vega: OTTHUMG00000020855”, “UniProt: O75612 Q86SR3 B4DSX9 X6R868”) | Pfam: MBOAT COX6C, Pfam: MBOAT MBOAT\_2, Pfam: Abhydrolase\_1 Abhydro\_lipase Hydrolase\_4 Abhydrolase\_6 DUF900 YjbF DUF779, Pfam: COesterase Mucin-like Abhydrolase\_3 Say1\_Mug180 | R01461, R01462 | C02530 | Chol. esters | Acyl-CoA:cholesterol O-acyltransferase, cholesterol ester acylhydrolase | c(“RC00004 C00010\_C00040”, “RC00055 C00187\_C02530”), RC00055 C00187\_C02530 | C00040 + C00187 <=> C00010 + C02530, C02530 + C00001 <=> C00187 + C00162 | Acyl-CoA + Cholesterol <=> CoA + Cholesterol ester, Cholesterol ester + H2O <=> Cholesterol + Fatty acid | 2.3.1.26, 3.1.1.13 | Product, Substrate |
| C24 GalCer | K00720, K01201, K07553, K12309, K17108 | ceramide glucosyltransferase [EC:2.4.1.80], glucosylceramidase [EC:3.2.1.45], beta-1,4-galactosyltransferase 6 [EC:2.4.1.274], beta-galactosidase [EC:3.2.1.23], non-lysosomal glucosylceramidase [EC:3.2.1.45] | 7357, 2629, 9331, 2720, 57704 | UGCG, GCS, GLCT1, GBA, GBA1, GCB, GLUC, B4GALT6, B4Gal-T6, beta4Gal-T6, GLB1, EBP, ELNR1, MPS4B, GBA2, AD035, NLGase, SPG46 | (RefSeq) UDP-glucose ceramide glucosyltransferase, (RefSeq) glucosylceramidase beta, (RefSeq) beta-1,4-galactosyltransferase 6, (RefSeq) galactosidase beta 1, (RefSeq) glucosylceramidase beta 2 | Homo sapiens (human) | c(hsa00600 = “Sphingolipid metabolism”, hsa01100 = “Metabolic pathways”), c(hsa00511 = “Other glycan degradation”, hsa00600 = “Sphingolipid metabolism”, hsa01100 = “Metabolic pathways”, hsa04142 = “Lysosome”), c(hsa00052 = “Galactose metabolism”, hsa00511 = “Other glycan degradation”, hsa00531 = “Glycosaminoglycan degradation”, hsa00600 = “Sphingolipid metabolism”, hsa00604 = “Glycosphingolipid biosynthesis - ganglio series”, hsa01100 = “Metabolic pathways”, hsa04142 = “Lysosome”), c(hsa00511 = “Other glycan degradation”, hsa00600 = “Sphingolipid metabolism”, hsa01100 = “Metabolic pathways”) | c(“NCBI-GeneID: 7357”, “NCBI-ProteinID: NP\_003349”, “OMIM: 602874”, “HGNC: 12524”, “Ensembl: ENSG00000148154”, “Vega: OTTHUMG00000020498”, “Pharos: Q16739(Tclin)”, “UniProt: Q16739 A0A024R157”), c(“NCBI-GeneID: 2629”, “NCBI-ProteinID: NP\_000148”, “OMIM: 606463”, “HGNC: 4177”, “Ensembl: ENSG00000177628”, “Vega: OTTHUMG00000035841”, “Pharos: P04062(Tchem)”, “UniProt: P04062 A0A068F658”), c(“NCBI-GeneID: 9331”, “NCBI-ProteinID: NP\_004766”, “OMIM: 604017”, “HGNC: 929”, “Ensembl: ENSG00000118276”, “Vega: OTTHUMG00000131980”, “Pharos: Q9UBX8(Tbio)”, “UniProt: Q9UBX8”), c(“NCBI-GeneID: 2720”, “NCBI-ProteinID: NP\_000395”, “OMIM: 611458”, “HGNC: 4298”, “Ensembl: ENSG00000170266”, “Vega: OTTHUMG00000155781”, “Pharos: P16278(Tchem)”, “UniProt: P16278”), c(“NCBI-GeneID: 57704”, “NCBI-ProteinID: NP\_065995”, “OMIM: 609471”, “HGNC: 18986”, “Ensembl: ENSG00000070610”, “Vega: OTTHUMG00000021024”, “Pharos: Q9HCG7(Tchem)”, “UniProt: Q9HCG7”) | Pfam: Glyco\_transf\_21 Glyco\_tranf\_2\_3 Glyco\_trans\_2\_3 Glycos\_transf\_2 Chitin\_synth\_2, Pfam: Glyco\_hydro\_30 Glyco\_hydro\_30C Glyco\_hydro\_59, Pfam: Glyco\_transf\_7N Glyco\_transf\_7C, Pfam: Glyco\_hydro\_35 Glyco\_hydro\_42 BetaGal\_dom4\_5, Pfam: DUF608 Glyco\_hydr\_116N Bac\_rhamnosid6H GDE\_C DUF5127 | R01497, R01498, R03354, R03355 | C01190 | HexCer | UDP-glucose:N-acylsphingosine D-glucosyltransferase, D-Glucosyl-N-acylsphingosine glucohydrolase, UDP-alpha-D-galactose:beta-D-glucosyl-(1<->1)-ceramide 4-beta-D-galactosyltransferase, beta-D-Galactosyl-1,4-beta-D-glucosylceramide galactohydrolase | c(“RC00005 C00015\_C00029”, “RC00059 C00195\_C01190”), c(“RC00059 C00195\_C01190”, “RC00451 C00031\_C01190”), c(“RC00005 C00015\_C00052”, “RC00049 C01190\_C01290”), RC00049 C00124\_C01290 C01190\_C01290 | C00029 + C00195 <=> C00015 + C01190, C01190 + C00001 <=> C00031 + C00195, C01190 + C00052 <=> C01290 + C00015, C01290 + C00001 <=> C01190 + C00124 | UDP-glucose + N-Acylsphingosine <=> UDP + Glucosylceramide, Glucosylceramide + H2O <=> D-Glucose + N-Acylsphingosine, Glucosylceramide + UDP-alpha-D-galactose <=> Lactosylceramide + UDP, Lactosylceramide + H2O <=> Glucosylceramide + D-Galactose | 2.4.1.80, c(“3.2.1.45”, “3.2.1.62”), 2.4.1.274, 3.2.1.23 | Product, Substrate |

##### When loading your own dataset

Load your own dataset. If your dataset has sample name in 1st column, enter the third parameter as true. If the dataset has 1st column as metabolite names, enter the third parameter as false. Please check the example files in the data folder of the package.

```
data_own <- separate_data('C:/MAMP/htdocs/webdev/human_cachexia.csv',"data",TRUE)

knitr::kable(head(data_own))
```

|  | metabolite\_name | PIF\_178 | PIF\_087 | PIF\_090 | NETL\_005\_V1 | PIF\_115 | PIF\_110 | NETL\_019\_V1 | NETCR\_014\_V1 | NETCR\_014\_V2 | PIF\_154 | NETL\_022\_V1 | NETL\_022\_V2 | NETL\_008\_V1 | PIF\_146 | PIF\_119 | PIF\_099 | PIF\_162 | PIF\_160 | PIF\_113 | PIF\_143 | NETCR\_007\_V1 | NETCR\_007\_V2 | PIF\_137 | PIF\_100 | NETL\_004\_V1 | PIF\_094 | PIF\_132 | PIF\_163 | NETCR\_003\_V1 | NETL\_028\_V1 | NETL\_028\_V2 | NETCR\_013\_V1 | NETL\_020\_V1 | NETL\_020\_V2 | PIF\_192 | NETCR\_012\_V1 | NETCR\_012\_V2 | PIF\_089 | NETCR\_002\_V1 | PIF\_179 | PIF\_114 | NETCR\_006\_V1 | PIF\_141 | NETCR\_025\_V1 | NETCR\_025\_V2 | NETCR\_016\_V1 | PIF\_116 | PIF\_191 | PIF\_164 | NETL\_013\_V1 | PIF\_188 | PIF\_195 | NETCR\_015\_V1 | PIF\_102 | NETL\_010\_V1 | NETL\_010\_V2 | NETL\_001\_V1 | NETCR\_015\_V2 | NETCR\_005\_V1 | PIF\_111 | PIF\_171 | NETCR\_008\_V1 | NETCR\_008\_V2 | NETL\_017\_V1 | NETL\_017\_V2 | NETL\_002\_V1 | NETL\_002\_V2 | PIF\_190 | NETCR\_009\_V1 | NETCR\_009\_V2 | NETL\_007\_V1 | PIF\_112 | NETCR\_019\_V2 | NETL\_012\_V1 | NETL\_012\_V2 | NETL\_003\_V1 | NETL\_003\_V2 |
| --- | --- | --- | --- | --- | --- | --- | --- | --- | --- | --- | --- | --- | --- | --- | --- | --- | --- | --- | --- | --- | --- | --- | --- | --- | --- | --- | --- | --- | --- | --- | --- | --- | --- | --- | --- | --- | --- | --- | --- | --- | --- | --- | --- | --- | --- | --- | --- | --- | --- | --- | --- | --- | --- | --- | --- | --- | --- | --- | --- | --- | --- | --- | --- | --- | --- | --- | --- | --- | --- | --- | --- | --- | --- | --- | --- | --- | --- | --- |
| 2 | 1,6-Anhydro-beta-D-glucose | 40.85 | 62.18 | 270.43 | 154.47 | 22.20 | 212.72 | 151.41 | 31.50 | 51.42 | 117.92 | 20.70 | 127.74 | 59.74 | 89.12 | 23.57 | 41.26 | 589.93 | 112.17 | 167.34 | 183.09 | 208.51 | 34.81 | 333.62 | 32.46 | 4.71 | 68.72 | 214.86 | 304.90 | 37.71 | 45.60 | 34.12 | 107.77 | 13.33 | 27.94 | 141.17 | 14.01 | 244.69 | 123.97 | 141.17 | 35.16 | 685.40 | 278.66 | 15.80 | 29.96 | 16.95 | 292.95 | 29.67 | 18.92 | 127.74 | 34.81 | 65.37 | 15.18 | 70.81 | 25.28 | 34.47 | 18.54 | 37.34 | 33.78 | 22.42 | 146.94 | 64.07 | 32.46 | 113.30 | 22.20 | 46.53 | 192.48 | 528.48 | 28.79 | 181.27 | 47.47 | 15.96 | 22.87 | 35.16 | 16.95 | 9.39 | 37.71 | 38.47 |
| 3 | 1-Methylnicotinamide | 65.37 | 340.36 | 64.72 | 52.98 | 73.70 | 31.82 | 36.60 | 6.82 | 30.27 | 52.46 | 221.41 | 177.68 | 50.91 | 32.79 | 6.89 | 8.67 | 21.98 | 25.28 | 19.89 | 90.92 | 53.52 | 95.58 | 35.87 | 9.68 | 11.13 | 13.87 | 127.74 | 25.79 | 10.80 | 473.43 | 92.76 | 16.61 | 50.91 | 80.64 | 68.03 | 46.06 | 116.75 | 81.45 | 28.50 | 26.58 | 36.23 | 40.45 | 23.57 | 96.54 | 114.43 | 57.97 | 70.11 | 24.53 | 1032.77 | 12.30 | 24.05 | 94.63 | 75.94 | 101.49 | 12.81 | 8.41 | 55.15 | 53.52 | 55.15 | 10.07 | 6.42 | 14.01 | 43.38 | 20.70 | 9.78 | 108.85 | 225.88 | 9.21 | 48.42 | 7.69 | 16.12 | 10.38 | 52.46 | 15.80 | 14.01 | 18.17 | 12.55 |
| 4 | 2-Aminobutyrate | 18.73 | 24.29 | 12.18 | 172.43 | 15.64 | 18.36 | 8.67 | 4.18 | 7.54 | 19.49 | 15.18 | 12.68 | 6.82 | 10.38 | 2.12 | 2.56 | 15.18 | 15.49 | 13.46 | 8.94 | 5.26 | 23.57 | 7.92 | 3.90 | 43.38 | 12.18 | 31.50 | 27.11 | 5.00 | 16.28 | 8.25 | 26.84 | 2.92 | 15.80 | 40.85 | 29.08 | 40.04 | 55.15 | 20.29 | 5.21 | 32.46 | 55.15 | 17.99 | 6.55 | 2.53 | 167.34 | 5.58 | 3.29 | 8.58 | 5.87 | 4.71 | 11.36 | 22.65 | 8.33 | 3.78 | 3.78 | 7.39 | 18.17 | 20.70 | 6.30 | 28.79 | 2.97 | 4.66 | 7.85 | 3.10 | 7.77 | 13.46 | 5.53 | 8.94 | 4.06 | 1.93 | 1.28 | 13.87 | 10.49 | 5.16 | 26.05 | 15.03 |
| 5 | 2-Hydroxyisobutyrate | 26.05 | 41.68 | 65.37 | 74.44 | 83.93 | 80.64 | 42.52 | 12.94 | 34.81 | 72.24 | 28.79 | 15.03 | 46.06 | 32.14 | 7.85 | 7.85 | 46.06 | 47.94 | 31.19 | 64.07 | 47.94 | 68.03 | 54.60 | 11.02 | 30.88 | 25.03 | 33.78 | 40.45 | 8.25 | 63.43 | 16.61 | 32.46 | 40.85 | 64.72 | 12.81 | 24.53 | 61.56 | 70.81 | 14.30 | 30.27 | 85.63 | 51.42 | 37.34 | 65.37 | 77.48 | 82.27 | 18.73 | 10.49 | 66.02 | 15.18 | 15.80 | 8.17 | 60.95 | 59.15 | 8.33 | 4.85 | 36.23 | 46.53 | 38.47 | 27.94 | 18.92 | 5.16 | 27.11 | 19.69 | 9.30 | 46.06 | 93.69 | 17.64 | 51.94 | 9.30 | 15.80 | 5.58 | 44.26 | 22.42 | 23.57 | 15.03 | 12.55 |
| 6 | 2-Oxoglutarate | 71.52 | 67.36 | 23.81 | 1199.91 | 33.12 | 47.94 | 223.63 | 25.03 | 80.64 | 73.70 | 357.81 | 68.03 | 111.05 | 32.46 | 8.33 | 6.89 | 32.79 | 28.79 | 47.94 | 20.49 | 212.72 | 287.15 | 20.49 | 170.72 | 104.58 | 28.22 | 88.23 | 70.81 | 11.70 | 221.41 | 55.15 | 62.80 | 46.99 | 88.23 | 26.05 | 64.07 | 174.16 | 92.76 | 97.51 | 7.39 | 25.03 | 74.44 | 21.33 | 1053.63 | 2465.13 | 468.72 | 5.53 | 9.68 | 38.09 | 16.78 | 7.24 | 5.64 | 230.44 | 88.23 | 14.30 | 8.08 | 75.94 | 81.45 | 164.02 | 24.05 | 85.63 | 8.08 | 22.42 | 38.47 | 10.59 | 55.15 | 230.44 | 14.44 | 982.40 | 65.37 | 25.28 | 8.50 | 99.48 | 62.80 | 46.99 | 23.34 | 22.20 |
| 7 | 3-Aminoisobutyrate | 1480.30 | 116.75 | 14.30 | 555.57 | 29.67 | 17.46 | 56.26 | 8.67 | 17.99 | 57.97 | 93.69 | 105.64 | 8.08 | 43.38 | 2.97 | 6.36 | 31.82 | 16.12 | 79.04 | 18.73 | 50.40 | 104.58 | 63.43 | 2.97 | 54.05 | 72.97 | 64.07 | 126.47 | 8.41 | 15.49 | 3.39 | 29.67 | 22.42 | 11.70 | 21.76 | 13.07 | 53.52 | 561.16 | 8.41 | 8.41 | 184.93 | 354.25 | 26.84 | 14.15 | 19.49 | 53.52 | 2.61 | 26.84 | 66.69 | 11.25 | 3.13 | 5.99 | 53.52 | 22.65 | 24.29 | 22.87 | 9.87 | 44.70 | 206.44 | 14.88 | 31.82 | 5.99 | 27.11 | 9.30 | 13.20 | 7.03 | 10.80 | 15.49 | 198.34 | 50.40 | 13.46 | 13.74 | 208.51 | 10.91 | 13.33 | 33.45 | 21.33 |

```
metadata_own <- separate_data('C:/MAMP/htdocs/webdev/human_cachexia.csv',"metadata",TRUE)
knitr::kable(head(metadata_own))
```

|  | local\_sample\_id | Muscle loss |
| --- | --- | --- |
| 2 | PIF\_178 | cachexic |
| 3 | PIF\_087 | cachexic |
| 4 | PIF\_090 | cachexic |
| 5 | NETL\_005\_V1 | cachexic |
| 6 | PIF\_115 | cachexic |
| 7 | PIF\_110 | cachexic |

```
refmet_names=convert_refmet(data_own)
knitr::kable(head(refmet_names))
```

| metabolite\_name | PIF\_178 | PIF\_087 | PIF\_090 | NETL\_005\_V1 | PIF\_115 | PIF\_110 | NETL\_019\_V1 | NETCR\_014\_V1 | NETCR\_014\_V2 | PIF\_154 | NETL\_022\_V1 | NETL\_022\_V2 | NETL\_008\_V1 | PIF\_146 | PIF\_119 | PIF\_099 | PIF\_162 | PIF\_160 | PIF\_113 | PIF\_143 | NETCR\_007\_V1 | NETCR\_007\_V2 | PIF\_137 | PIF\_100 | NETL\_004\_V1 | PIF\_094 | PIF\_132 | PIF\_163 | NETCR\_003\_V1 | NETL\_028\_V1 | NETL\_028\_V2 | NETCR\_013\_V1 | NETL\_020\_V1 | NETL\_020\_V2 | PIF\_192 | NETCR\_012\_V1 | NETCR\_012\_V2 | PIF\_089 | NETCR\_002\_V1 | PIF\_179 | PIF\_114 | NETCR\_006\_V1 | PIF\_141 | NETCR\_025\_V1 | NETCR\_025\_V2 | NETCR\_016\_V1 | PIF\_116 | PIF\_191 | PIF\_164 | NETL\_013\_V1 | PIF\_188 | PIF\_195 | NETCR\_015\_V1 | PIF\_102 | NETL\_010\_V1 | NETL\_010\_V2 | NETL\_001\_V1 | NETCR\_015\_V2 | NETCR\_005\_V1 | PIF\_111 | PIF\_171 | NETCR\_008\_V1 | NETCR\_008\_V2 | NETL\_017\_V1 | NETL\_017\_V2 | NETL\_002\_V1 | NETL\_002\_V2 | PIF\_190 | NETCR\_009\_V1 | NETCR\_009\_V2 | NETL\_007\_V1 | PIF\_112 | NETCR\_019\_V2 | NETL\_012\_V1 | NETL\_012\_V2 | NETL\_003\_V1 | NETL\_003\_V2 | refmet\_name | formula | super\_class | main\_class | sub\_class |
| --- | --- | --- | --- | --- | --- | --- | --- | --- | --- | --- | --- | --- | --- | --- | --- | --- | --- | --- | --- | --- | --- | --- | --- | --- | --- | --- | --- | --- | --- | --- | --- | --- | --- | --- | --- | --- | --- | --- | --- | --- | --- | --- | --- | --- | --- | --- | --- | --- | --- | --- | --- | --- | --- | --- | --- | --- | --- | --- | --- | --- | --- | --- | --- | --- | --- | --- | --- | --- | --- | --- | --- | --- | --- | --- | --- | --- | --- | --- | --- | --- | --- | --- |
| 1-Methylnicotinamide | 65.37 | 340.36 | 64.72 | 52.98 | 73.70 | 31.82 | 36.60 | 6.82 | 30.27 | 52.46 | 221.41 | 177.68 | 50.91 | 32.79 | 6.89 | 8.67 | 21.98 | 25.28 | 19.89 | 90.92 | 53.52 | 95.58 | 35.87 | 9.68 | 11.13 | 13.87 | 127.74 | 25.79 | 10.80 | 473.43 | 92.76 | 16.61 | 50.91 | 80.64 | 68.03 | 46.06 | 116.75 | 81.45 | 28.50 | 26.58 | 36.23 | 40.45 | 23.57 | 96.54 | 114.43 | 57.97 | 70.11 | 24.53 | 1032.77 | 12.30 | 24.05 | 94.63 | 75.94 | 101.49 | 12.81 | 8.41 | 55.15 | 53.52 | 55.15 | 10.07 | 6.42 | 14.01 | 43.38 | 20.70 | 9.78 | 108.85 | 225.88 | 9.21 | 48.42 | 7.69 | 16.12 | 10.38 | 52.46 | 15.80 | 14.01 | 18.17 | 12.55 | 1-Methyl nicotinamide | C7H9N2O | Organoheterocyclic compounds | Pyridinecarboxylic acids | Nicotinamides |
| 1,6-Anhydro-beta-D-glucose | 40.85 | 62.18 | 270.43 | 154.47 | 22.20 | 212.72 | 151.41 | 31.50 | 51.42 | 117.92 | 20.70 | 127.74 | 59.74 | 89.12 | 23.57 | 41.26 | 589.93 | 112.17 | 167.34 | 183.09 | 208.51 | 34.81 | 333.62 | 32.46 | 4.71 | 68.72 | 214.86 | 304.90 | 37.71 | 45.60 | 34.12 | 107.77 | 13.33 | 27.94 | 141.17 | 14.01 | 244.69 | 123.97 | 141.17 | 35.16 | 685.40 | 278.66 | 15.80 | 29.96 | 16.95 | 292.95 | 29.67 | 18.92 | 127.74 | 34.81 | 65.37 | 15.18 | 70.81 | 25.28 | 34.47 | 18.54 | 37.34 | 33.78 | 22.42 | 146.94 | 64.07 | 32.46 | 113.30 | 22.20 | 46.53 | 192.48 | 528.48 | 28.79 | 181.27 | 47.47 | 15.96 | 22.87 | 35.16 | 16.95 | 9.39 | 37.71 | 38.47 | Glucosan | C6H10O5 | Organoheterocyclic compounds | Oxepanes | Oxepanes |
| 2-Aminobutyrate | 18.73 | 24.29 | 12.18 | 172.43 | 15.64 | 18.36 | 8.67 | 4.18 | 7.54 | 19.49 | 15.18 | 12.68 | 6.82 | 10.38 | 2.12 | 2.56 | 15.18 | 15.49 | 13.46 | 8.94 | 5.26 | 23.57 | 7.92 | 3.90 | 43.38 | 12.18 | 31.50 | 27.11 | 5.00 | 16.28 | 8.25 | 26.84 | 2.92 | 15.80 | 40.85 | 29.08 | 40.04 | 55.15 | 20.29 | 5.21 | 32.46 | 55.15 | 17.99 | 6.55 | 2.53 | 167.34 | 5.58 | 3.29 | 8.58 | 5.87 | 4.71 | 11.36 | 22.65 | 8.33 | 3.78 | 3.78 | 7.39 | 18.17 | 20.70 | 6.30 | 28.79 | 2.97 | 4.66 | 7.85 | 3.10 | 7.77 | 13.46 | 5.53 | 8.94 | 4.06 | 1.93 | 1.28 | 13.87 | 10.49 | 5.16 | 26.05 | 15.03 | 2-Aminobutyric acid | C4H9NO2 | Fatty Acyls | Fatty acids | Amino FA |
| 2-Hydroxyisobutyrate | 26.05 | 41.68 | 65.37 | 74.44 | 83.93 | 80.64 | 42.52 | 12.94 | 34.81 | 72.24 | 28.79 | 15.03 | 46.06 | 32.14 | 7.85 | 7.85 | 46.06 | 47.94 | 31.19 | 64.07 | 47.94 | 68.03 | 54.60 | 11.02 | 30.88 | 25.03 | 33.78 | 40.45 | 8.25 | 63.43 | 16.61 | 32.46 | 40.85 | 64.72 | 12.81 | 24.53 | 61.56 | 70.81 | 14.30 | 30.27 | 85.63 | 51.42 | 37.34 | 65.37 | 77.48 | 82.27 | 18.73 | 10.49 | 66.02 | 15.18 | 15.80 | 8.17 | 60.95 | 59.15 | 8.33 | 4.85 | 36.23 | 46.53 | 38.47 | 27.94 | 18.92 | 5.16 | 27.11 | 19.69 | 9.30 | 46.06 | 93.69 | 17.64 | 51.94 | 9.30 | 15.80 | 5.58 | 44.26 | 22.42 | 23.57 | 15.03 | 12.55 | 2-Hydroxyisobutyric acid | C4H8O3 | Fatty Acyls | Fatty acids | Hydroxy FA |
| 2-Oxoglutarate | 71.52 | 67.36 | 23.81 | 1199.91 | 33.12 | 47.94 | 223.63 | 25.03 | 80.64 | 73.70 | 357.81 | 68.03 | 111.05 | 32.46 | 8.33 | 6.89 | 32.79 | 28.79 | 47.94 | 20.49 | 212.72 | 287.15 | 20.49 | 170.72 | 104.58 | 28.22 | 88.23 | 70.81 | 11.70 | 221.41 | 55.15 | 62.80 | 46.99 | 88.23 | 26.05 | 64.07 | 174.16 | 92.76 | 97.51 | 7.39 | 25.03 | 74.44 | 21.33 | 1053.63 | 2465.13 | 468.72 | 5.53 | 9.68 | 38.09 | 16.78 | 7.24 | 5.64 | 230.44 | 88.23 | 14.30 | 8.08 | 75.94 | 81.45 | 164.02 | 24.05 | 85.63 | 8.08 | 22.42 | 38.47 | 10.59 | 55.15 | 230.44 | 14.44 | 982.40 | 65.37 | 25.28 | 8.50 | 99.48 | 62.80 | 46.99 | 23.34 | 22.20 | Oxoglutaric acid | C5H6O5 | Organic acids | TCA acids | TCA acids |
| 3-Aminoisobutyrate | 1480.30 | 116.75 | 14.30 | 555.57 | 29.67 | 17.46 | 56.26 | 8.67 | 17.99 | 57.97 | 93.69 | 105.64 | 8.08 | 43.38 | 2.97 | 6.36 | 31.82 | 16.12 | 79.04 | 18.73 | 50.40 | 104.58 | 63.43 | 2.97 | 54.05 | 72.97 | 64.07 | 126.47 | 8.41 | 15.49 | 3.39 | 29.67 | 22.42 | 11.70 | 21.76 | 13.07 | 53.52 | 561.16 | 8.41 | 8.41 | 184.93 | 354.25 | 26.84 | 14.15 | 19.49 | 53.52 | 2.61 | 26.84 | 66.69 | 11.25 | 3.13 | 5.99 | 53.52 | 22.65 | 24.29 | 22.87 | 9.87 | 44.70 | 206.44 | 14.88 | 31.82 | 5.99 | 27.11 | 9.30 | 13.20 | 7.03 | 10.80 | 15.49 | 198.34 | 50.40 | 13.46 | 13.74 | 208.51 | 10.91 | 13.33 | 33.45 | 21.33 | 3-Amino-isobutanoic acid | C4H9NO2 | Fatty Acyls | Fatty acids | Amino FA |

##### Get significant metabolite via same function significant\_met. You may need to change the parameters accordingly

```
stats_metabolites = significant_met_own(metabolomics_data=refmet_names,met_col="metabolite_name",metadata=metadata_own, factor1='cachexic', factor2='control', factor_col='Muscle loss',sample_col='local_sample_id', p_adjust='fdr',normalization="50percent")
```

```
sig_metabolites = stats_metabolites[which(stats_metabolites[,"pval"] <= 0.05&abs(stats_metabolites[,"log2Fold_change"])>0.5),]
```

```
plot_volcano(stats_metabolites, thres_pval= 0.05,thres_log2foldchange = 0.5, TRUE)
```

```
sig_metabolites_kegg_id= map_keggid(sig_metabolites)
```

```
count_changes = metcountplot(df_metclass=sig_metabolites_kegg_id, metclass='sub_class', plotting=TRUE, thres_logfC = 0.5)
count_changes$plotimg
```

```
metenrichment = metclassenrichment(df_metclass=sig_metabolites_kegg_id, refmet_names, metclass= 'sub_class',enrich_stats="HG",no=1)
```

```
plot_met_enrichment(metenrichment, metclass='sub_class',"HG", no=1)
```

The rest of the pipeline can be followed similarly.
