## Supplementary figures and images for "MetENP/MetENPWeb: An R package and web application for metabolomics enrichment and pathway analysis in Metabolomics Workbench"

### dotplot.pdf

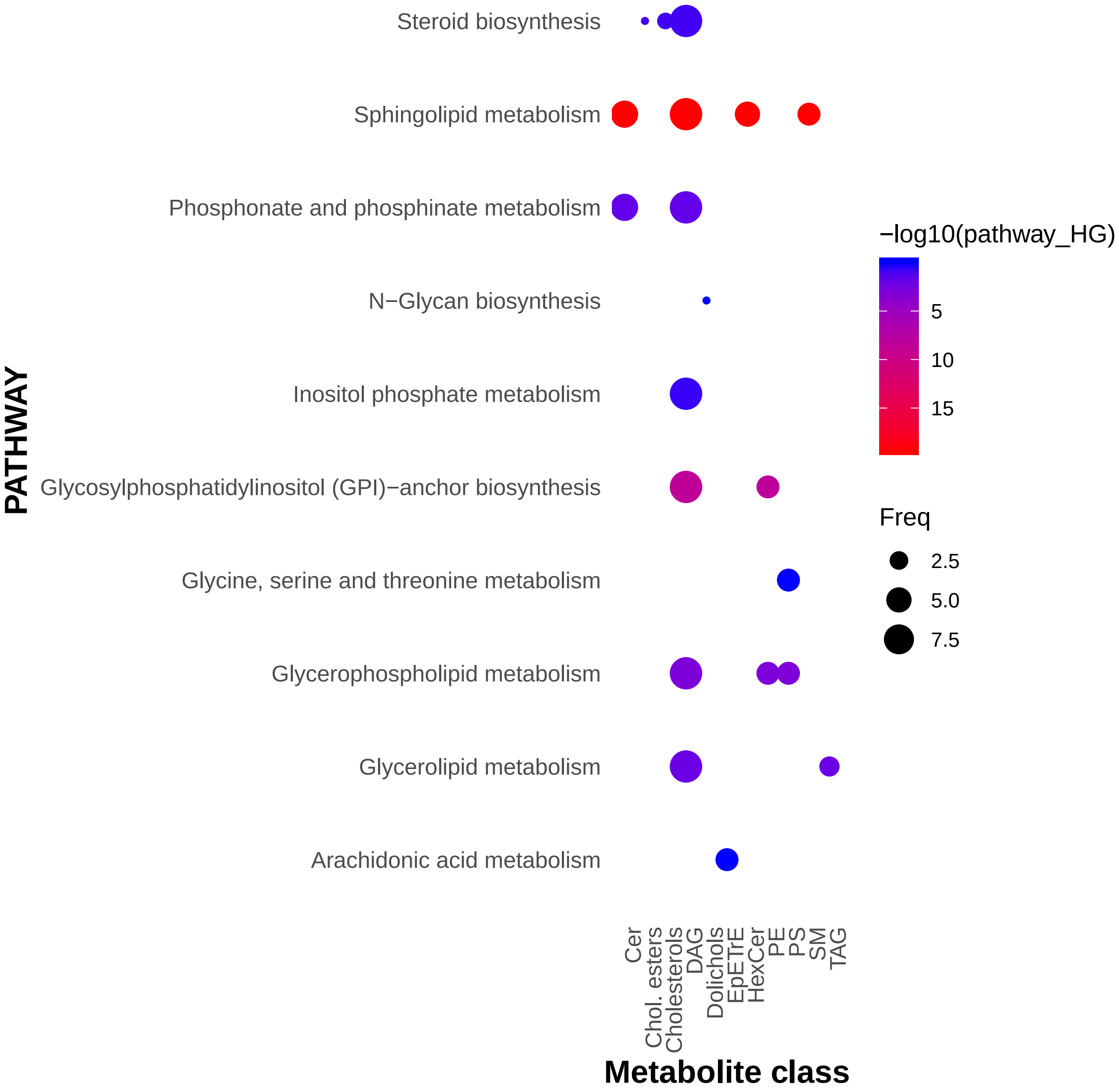

### dotplot.pdf

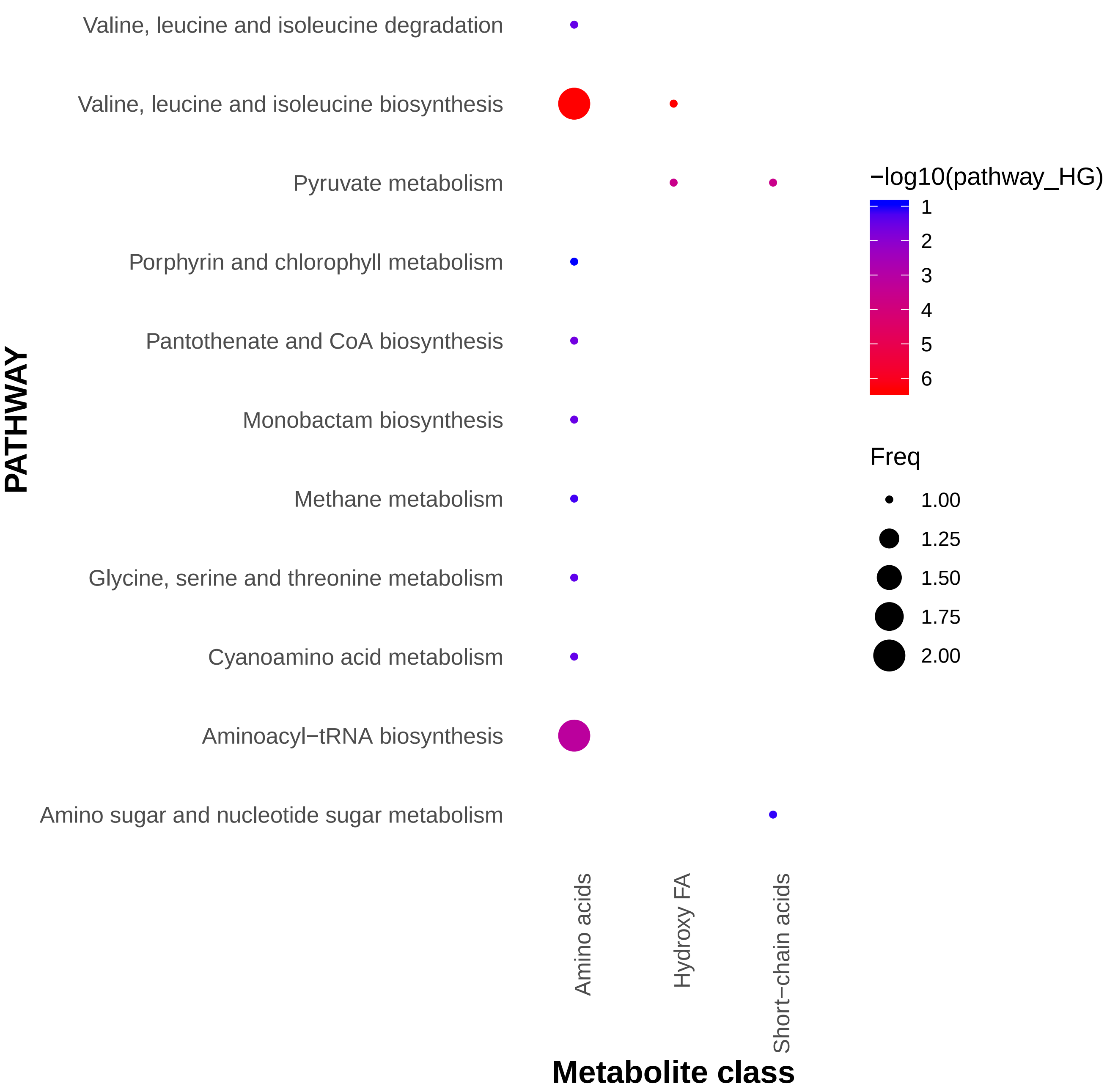

### dotplot.pdf

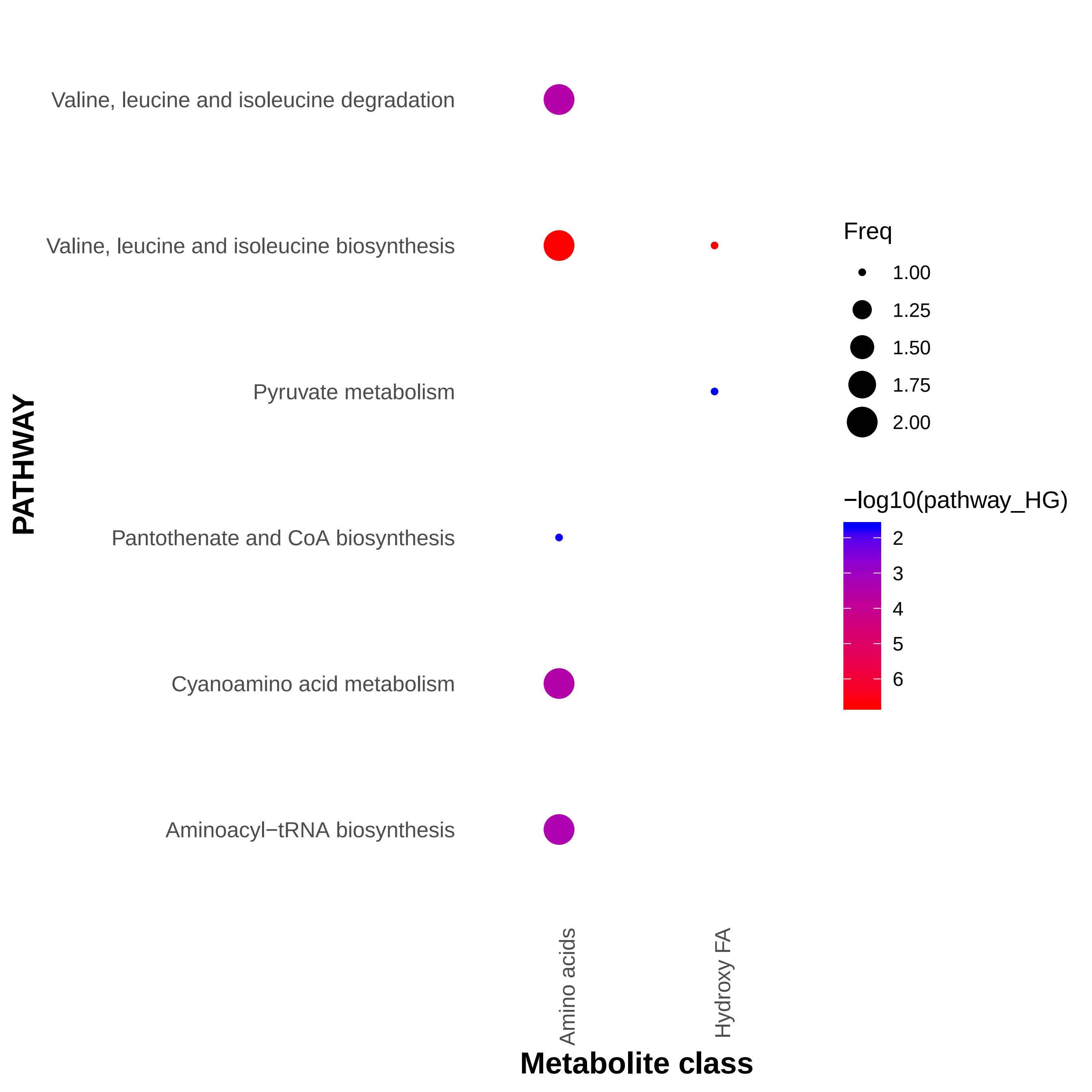

### dotplot.pdf

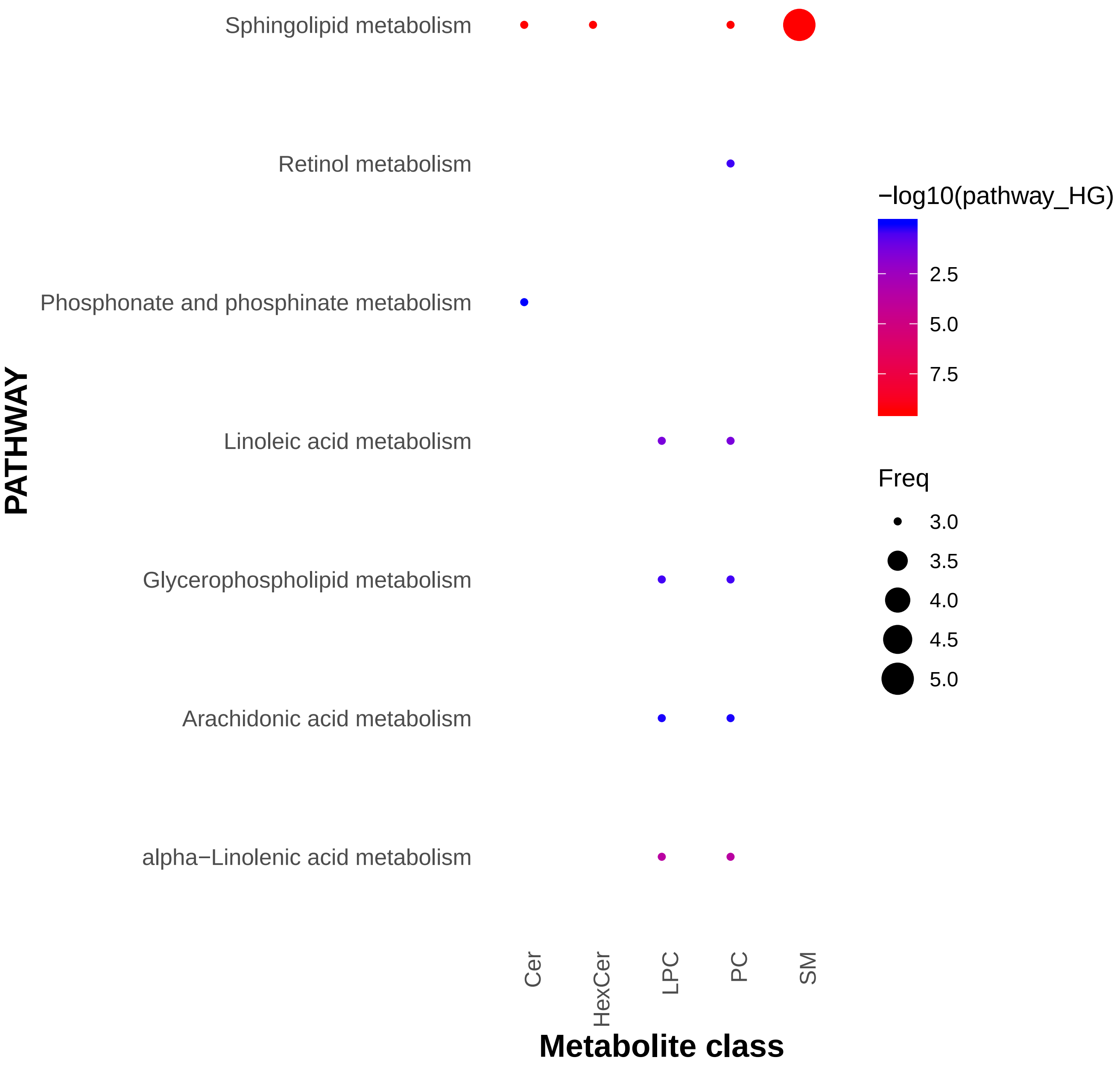

### heatmap.pdf

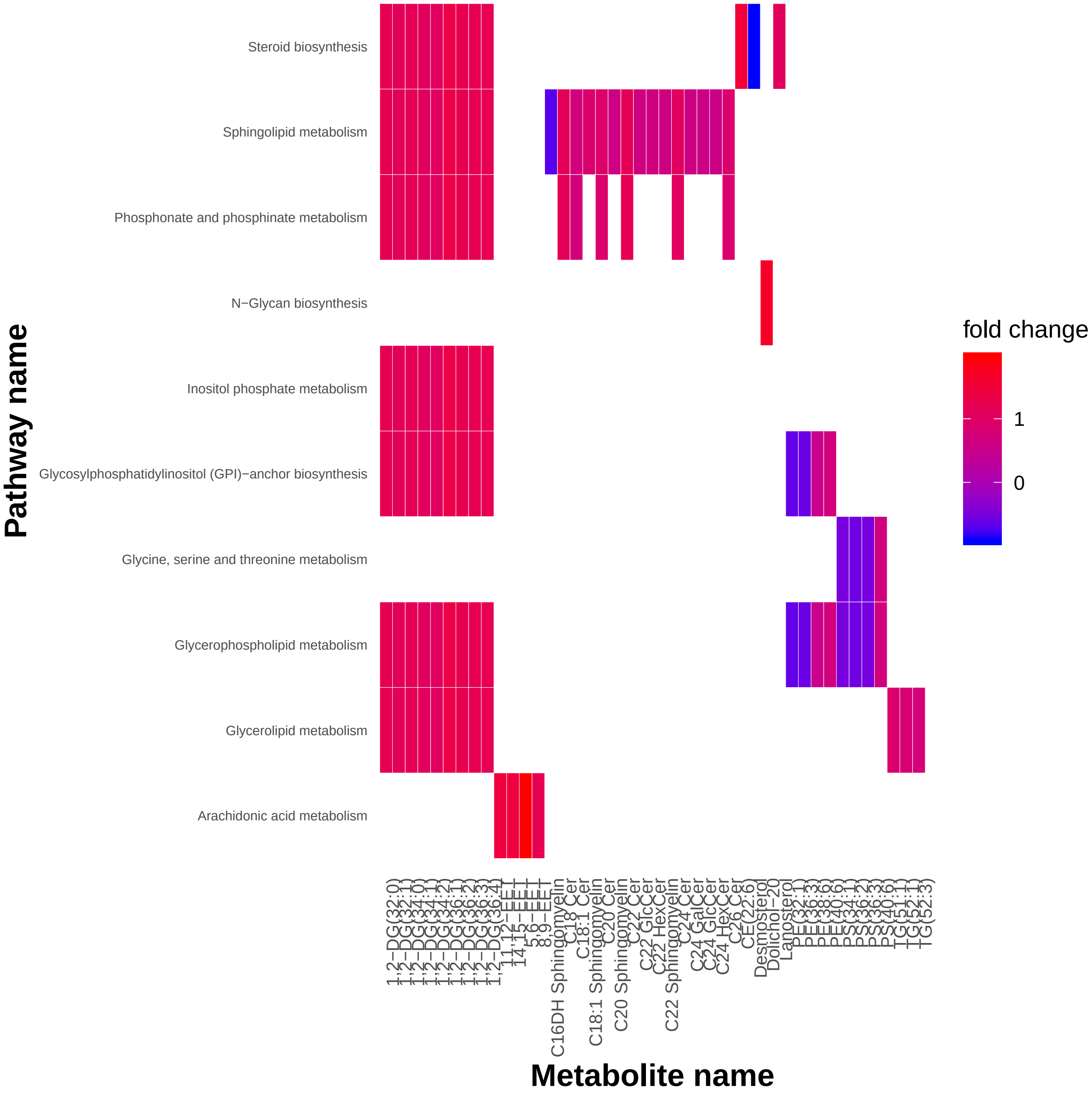

### heatmap.pdf

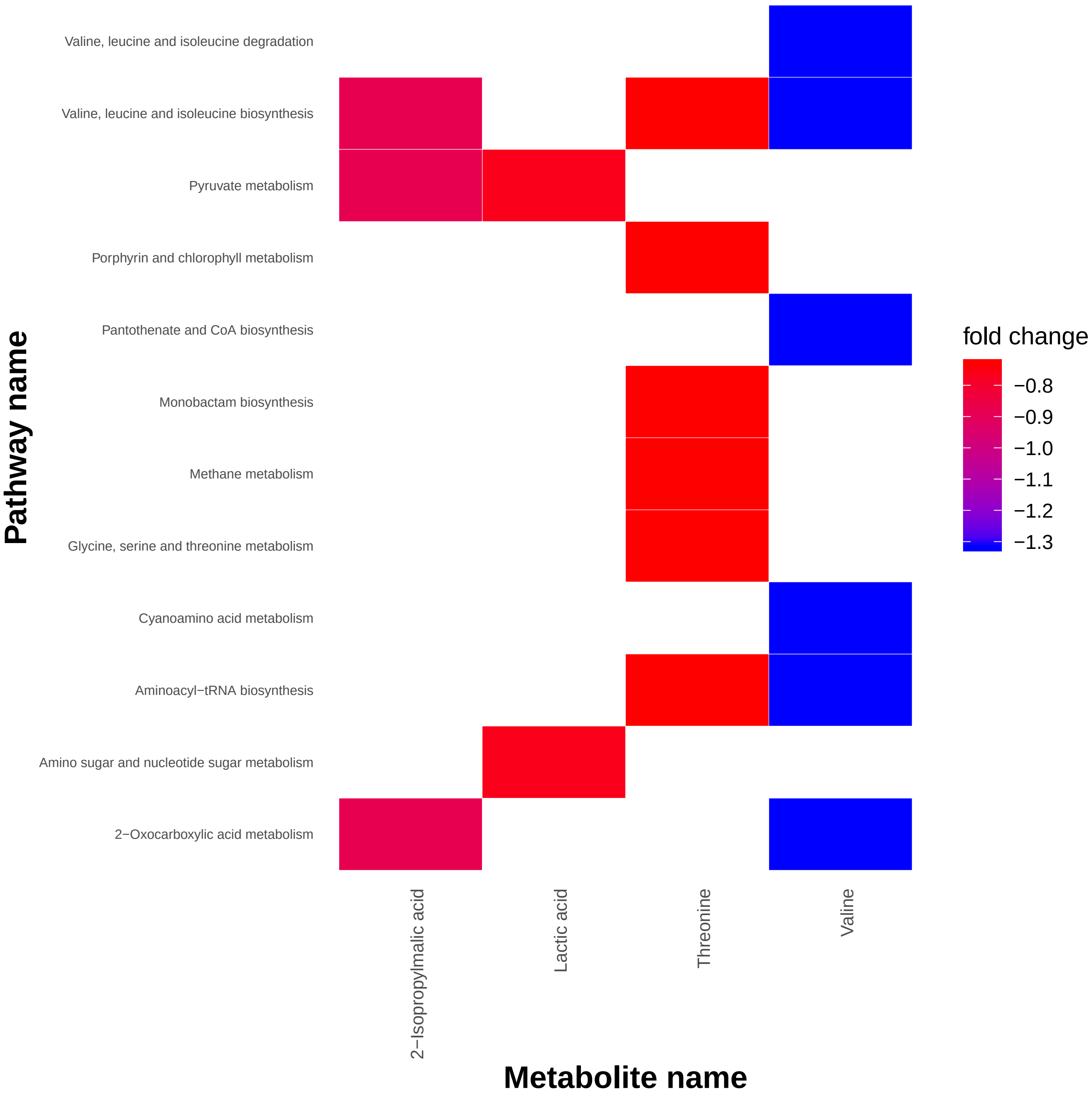

### heatmap.pdf

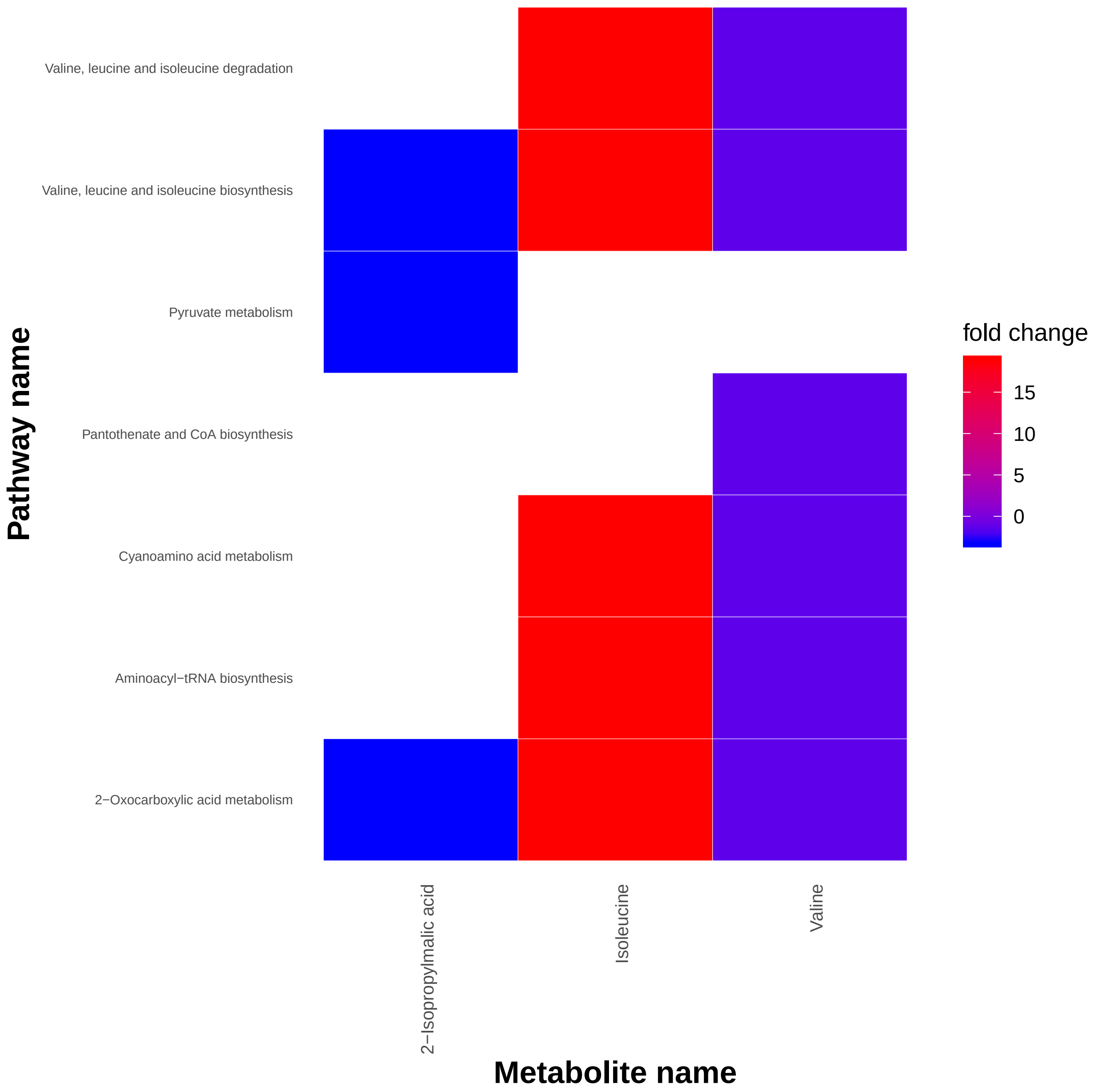

### heatmap.pdf

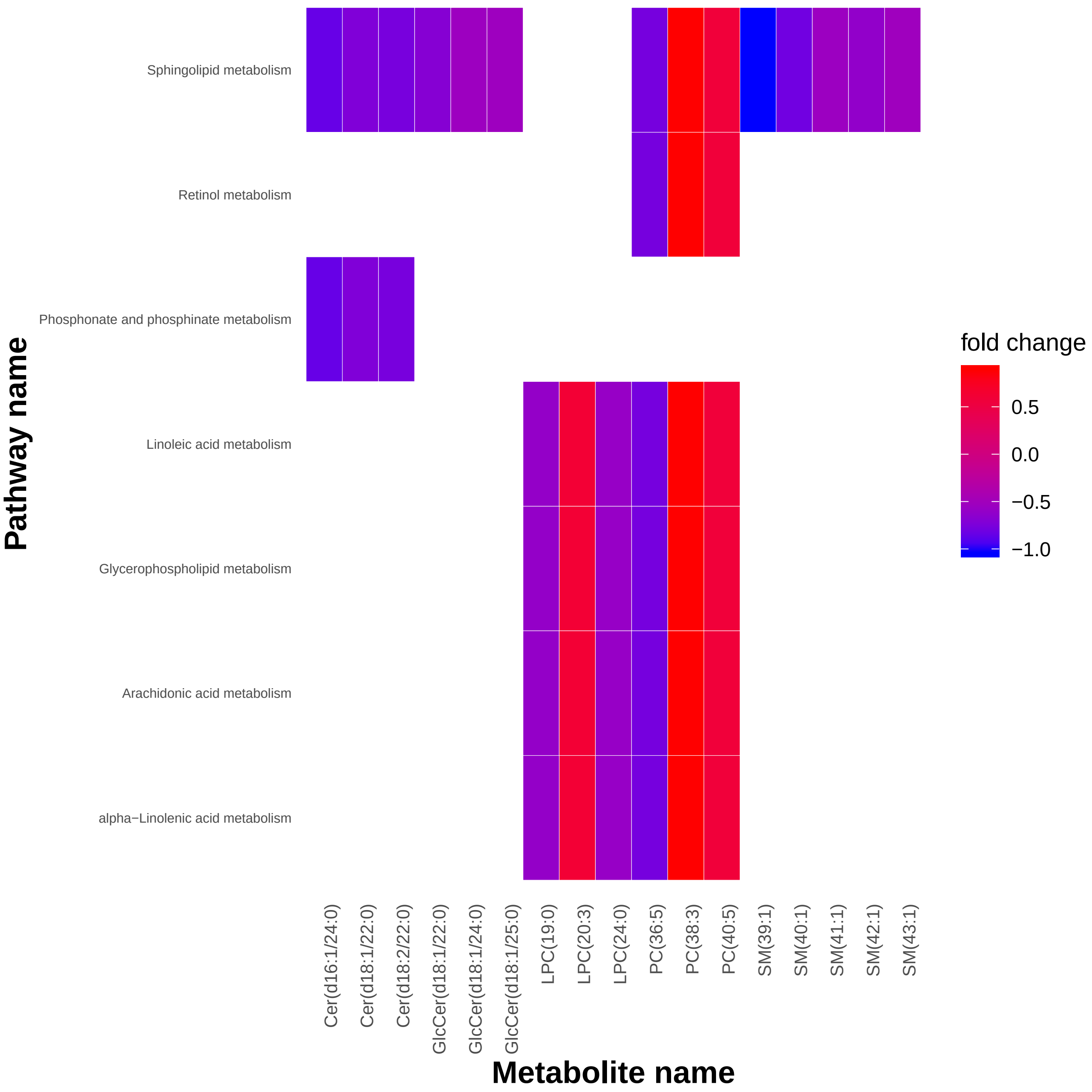

### metcountplot.pdf

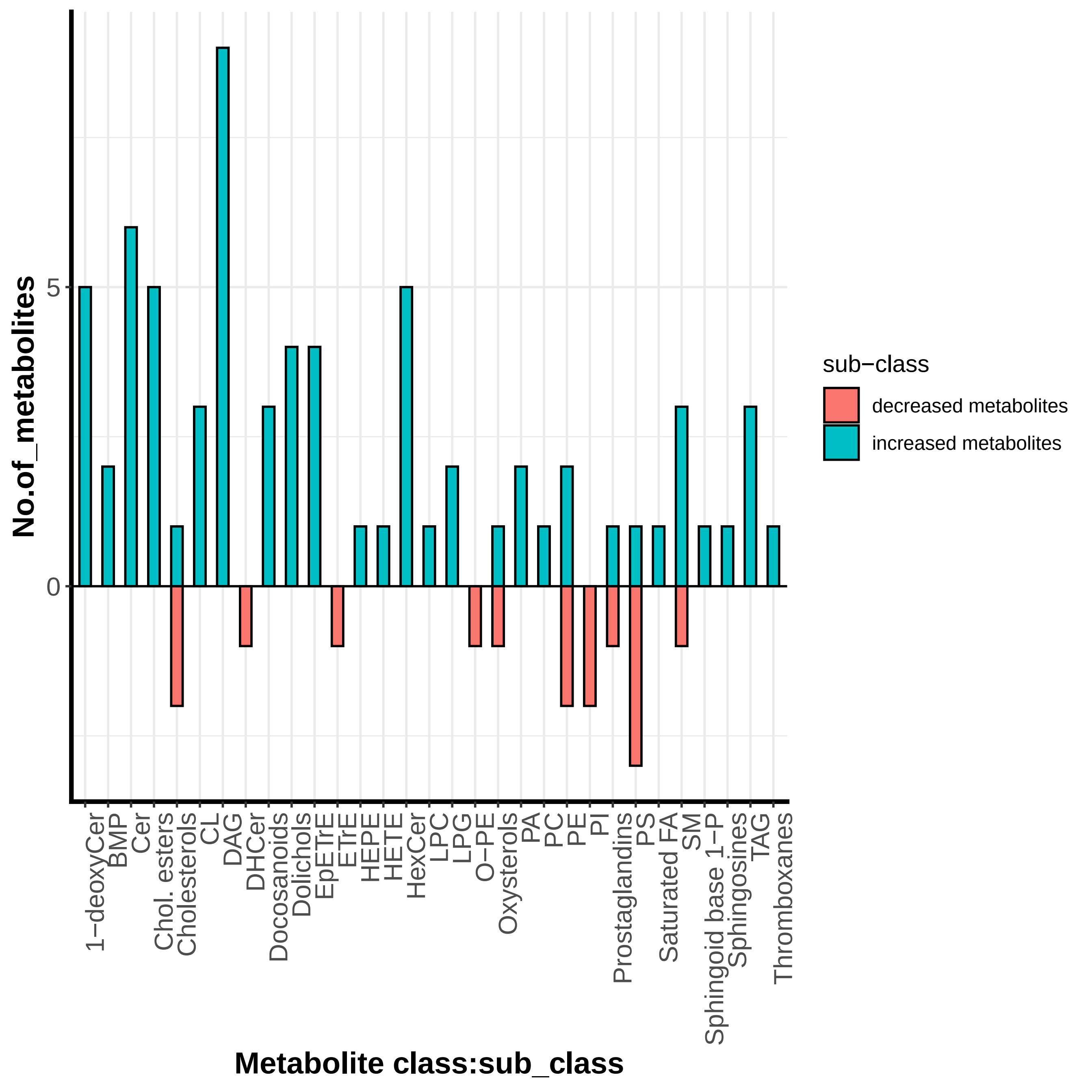

### metcountplot.pdf

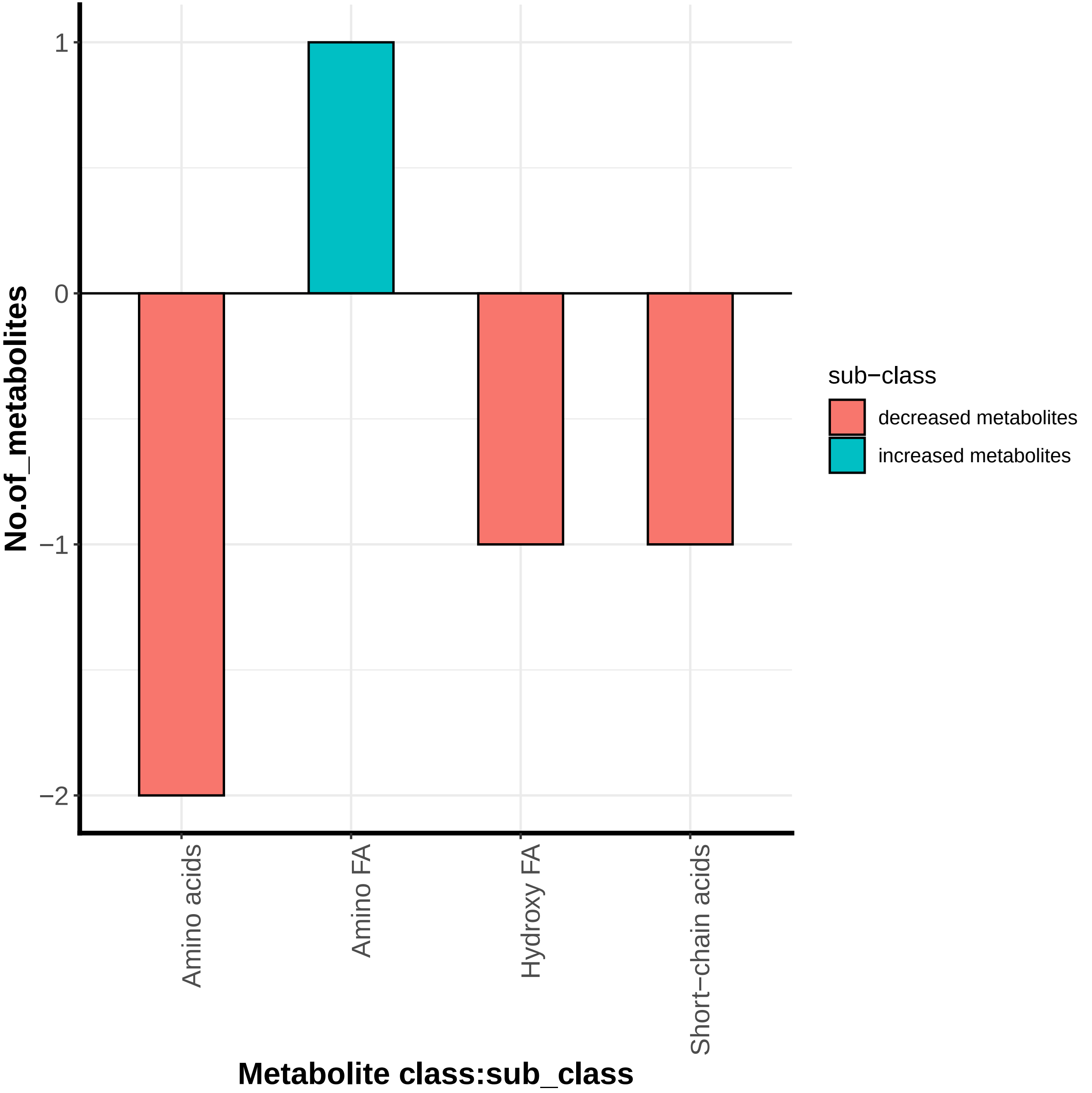

### metcountplot.pdf

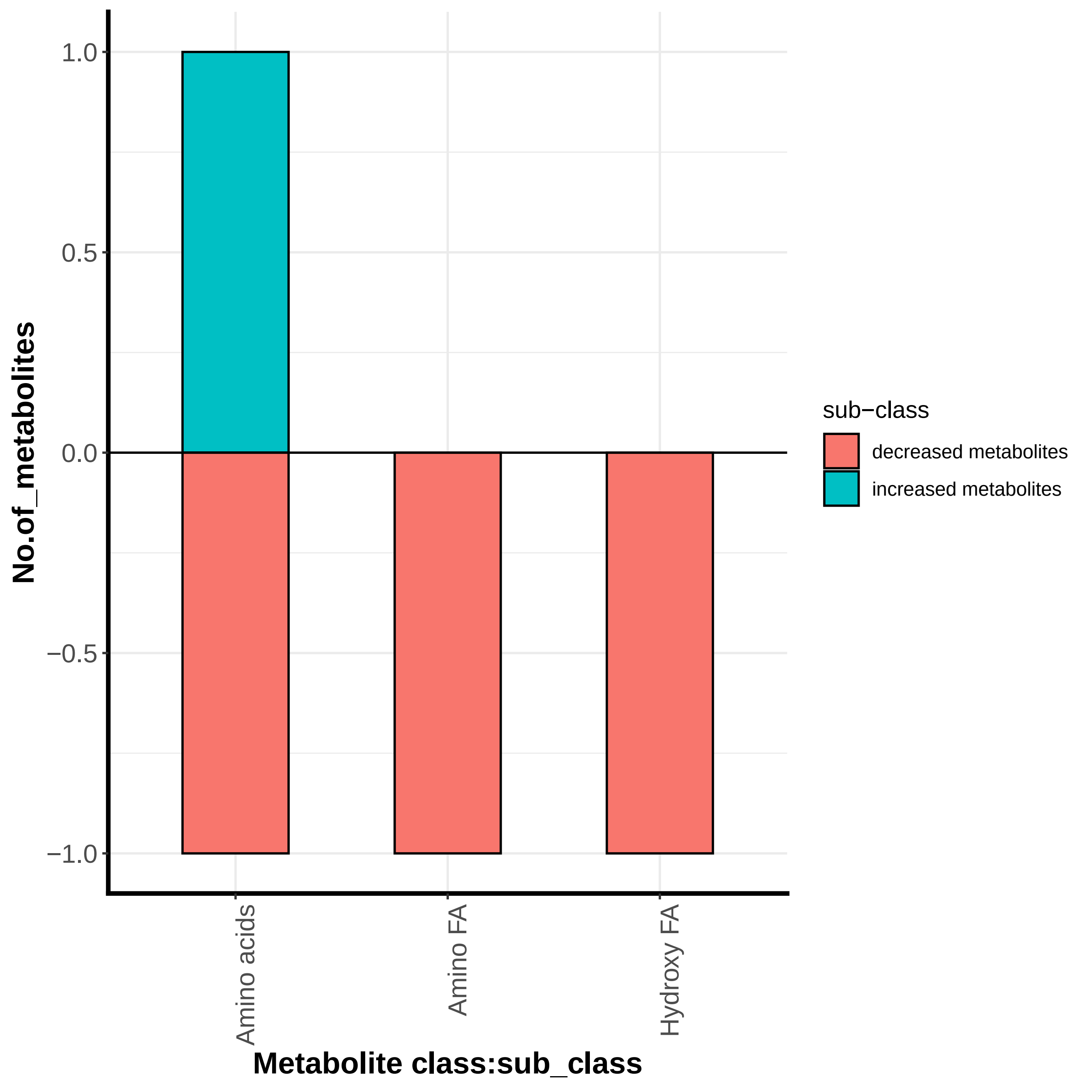

### metcountplot.pdf

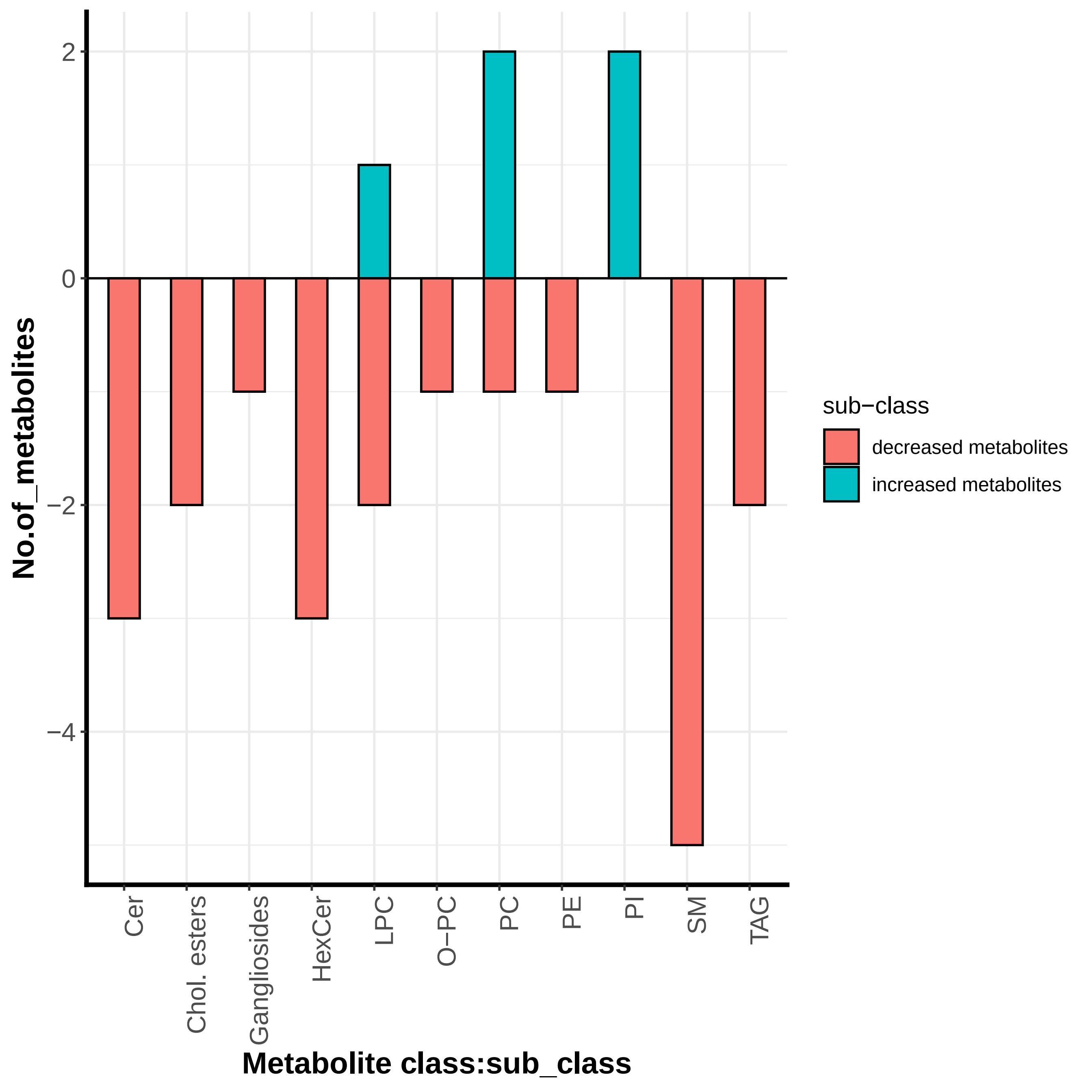

### metenrichment.pdf

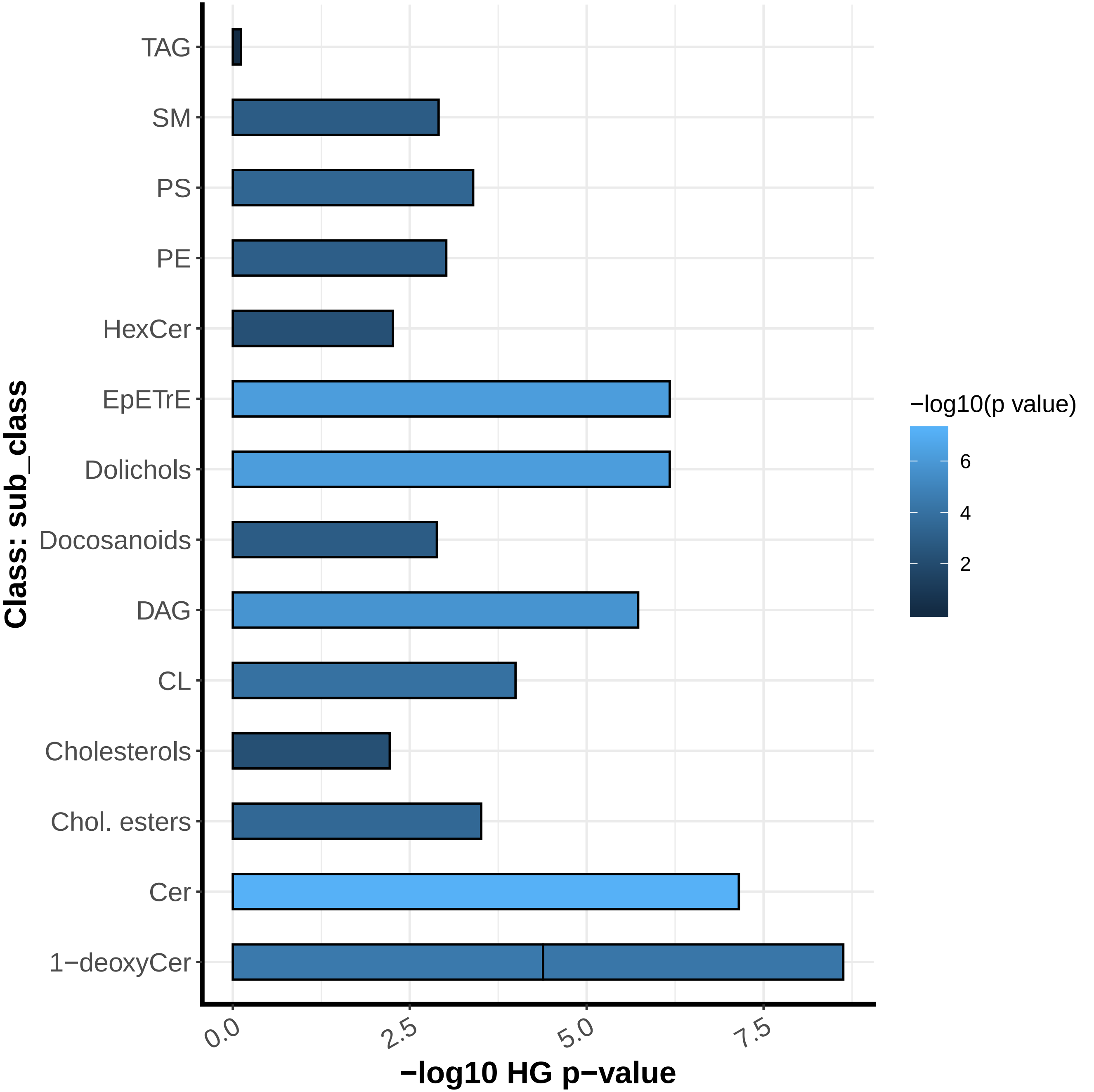

### metenrichment.pdf

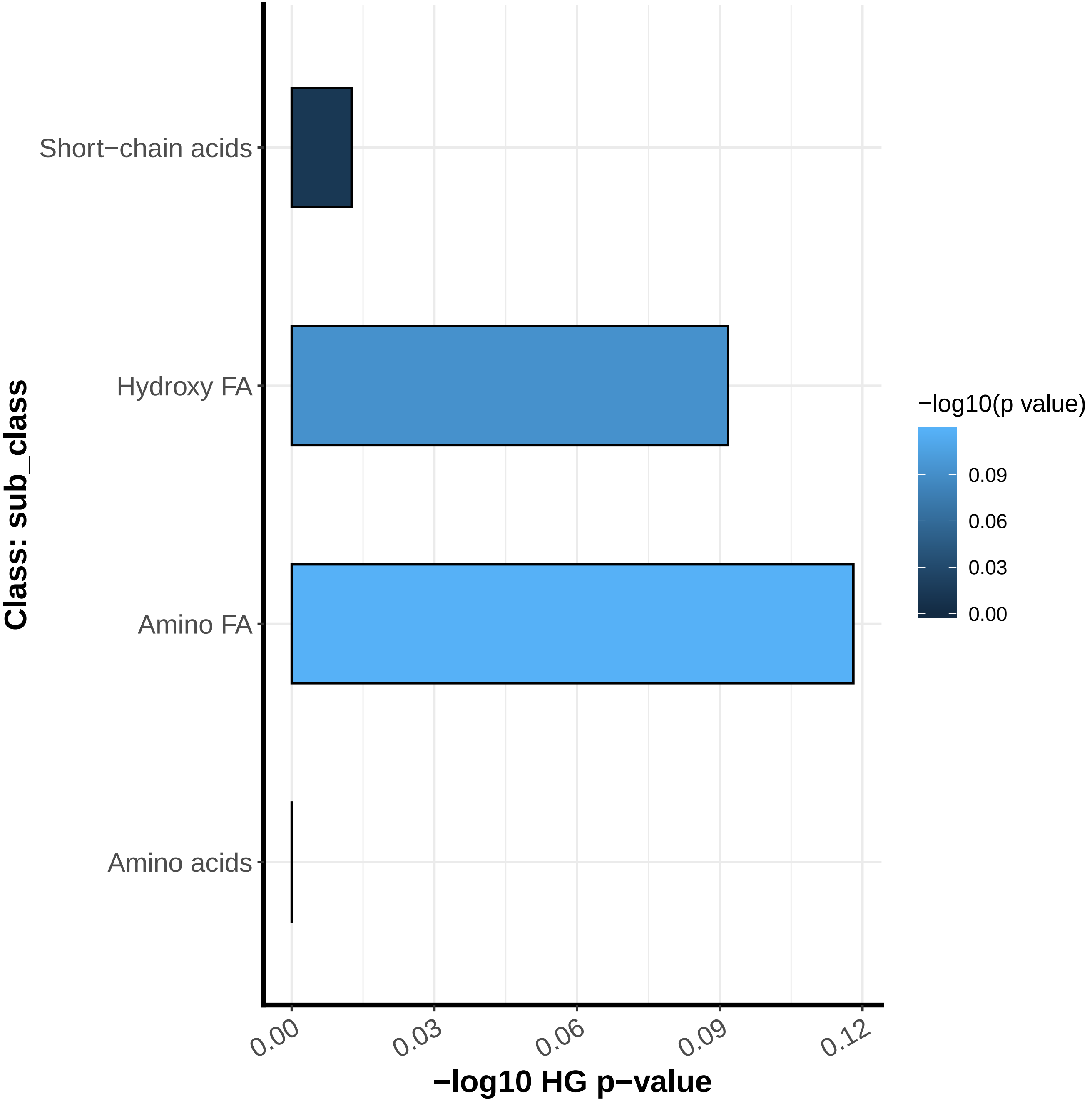

### metenrichment.pdf

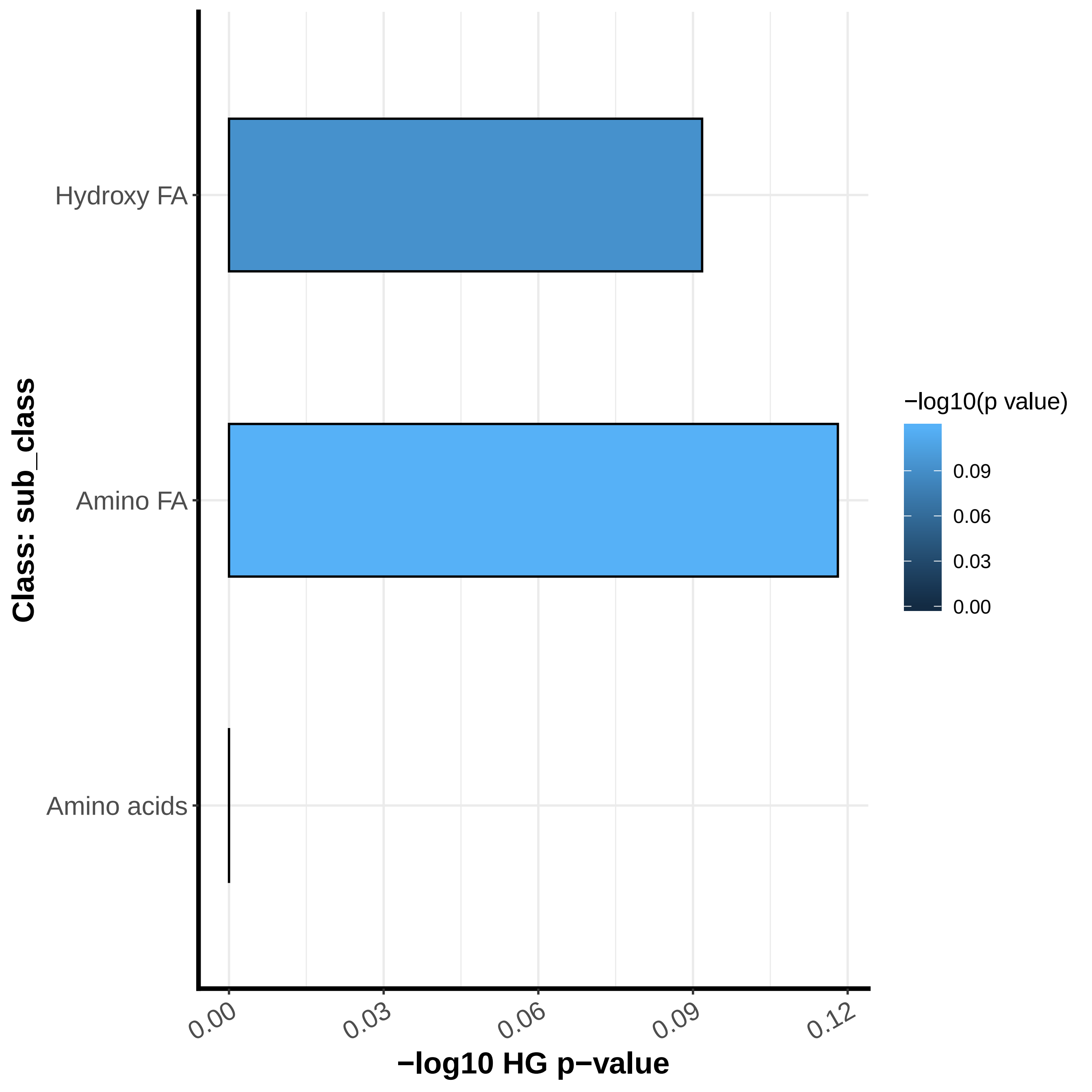

### metenrichment.pdf

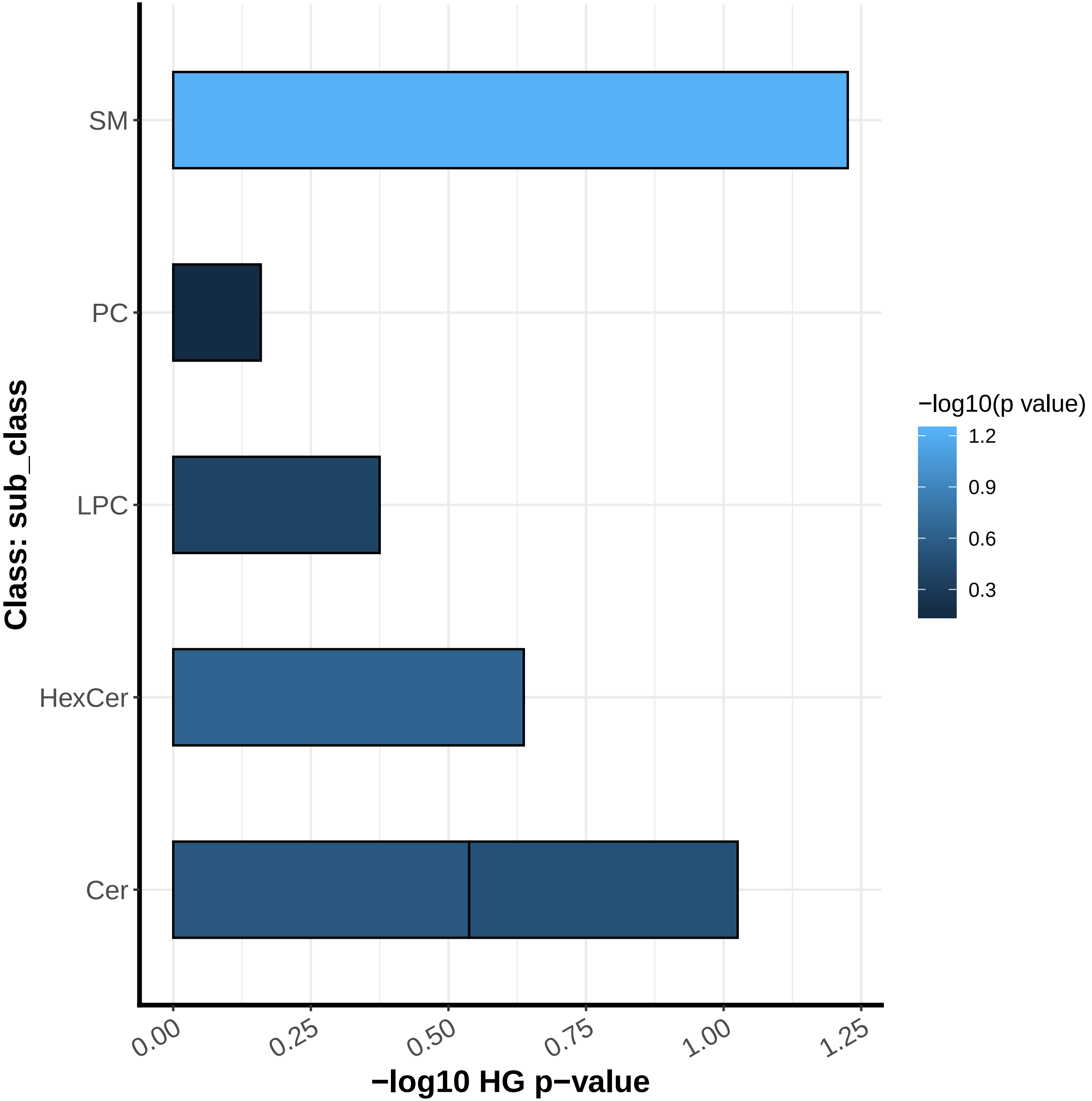

### pathway_network.pdf

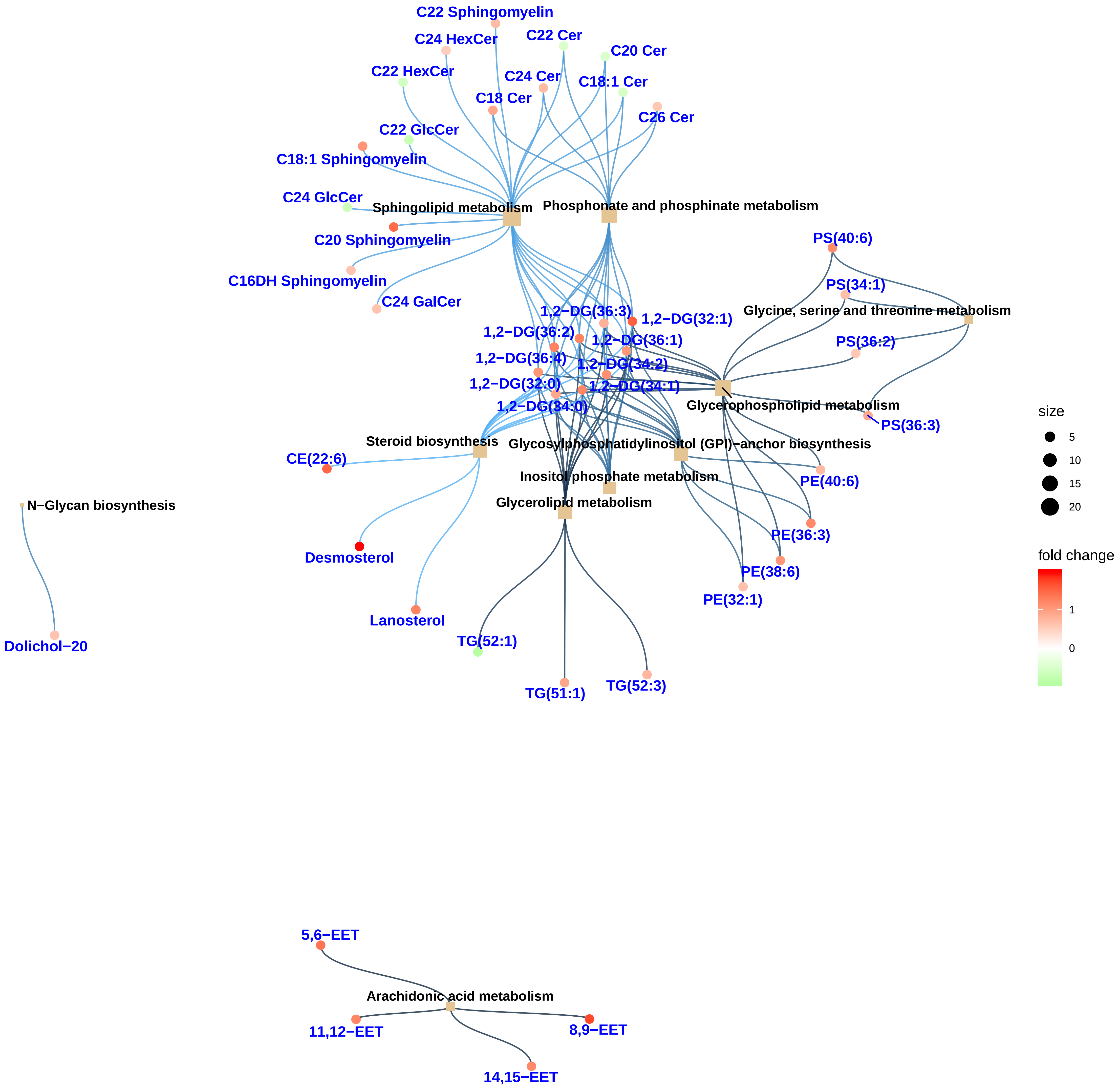

### pathway_network.pdf

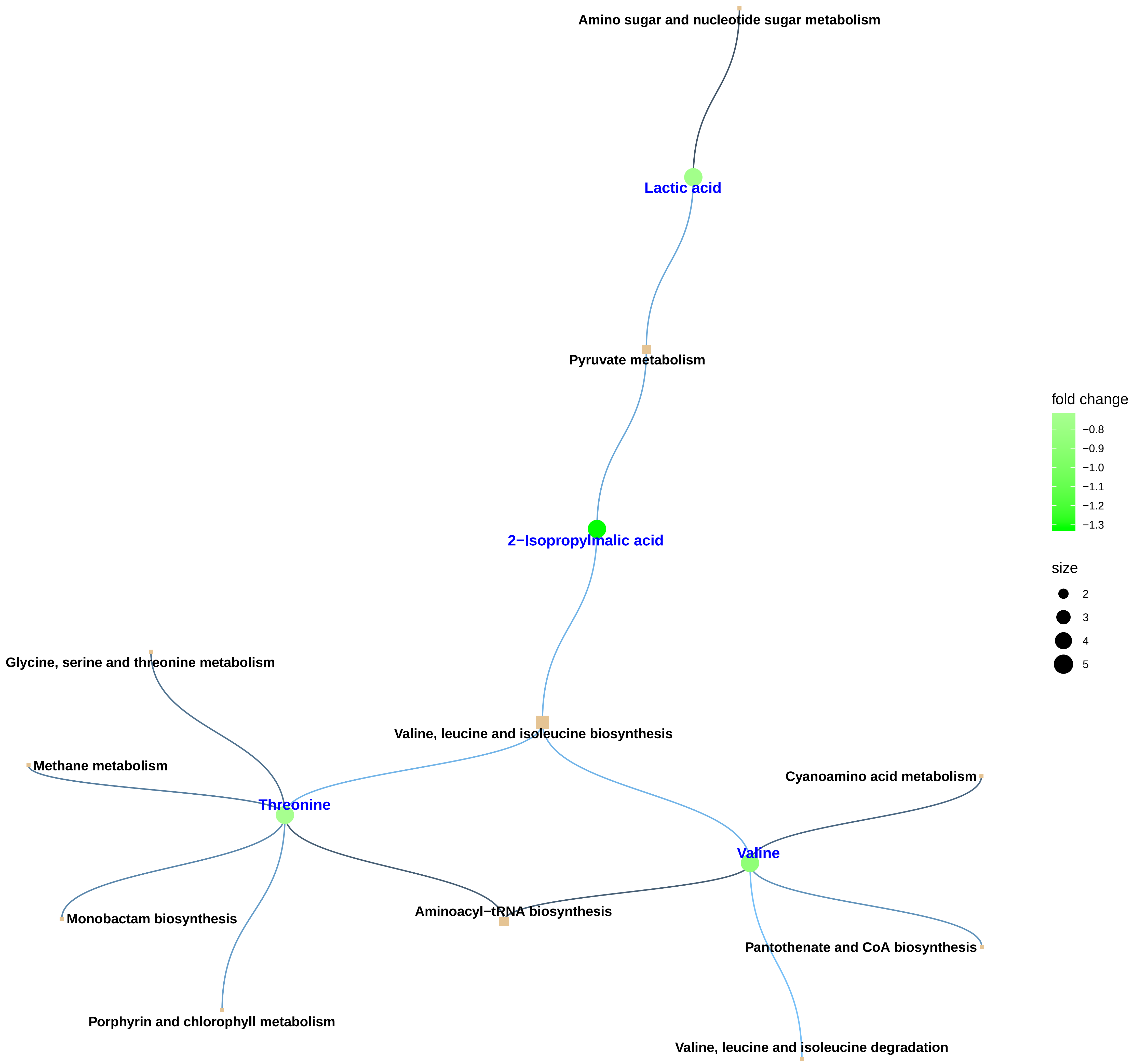

### pathway_network.pdf

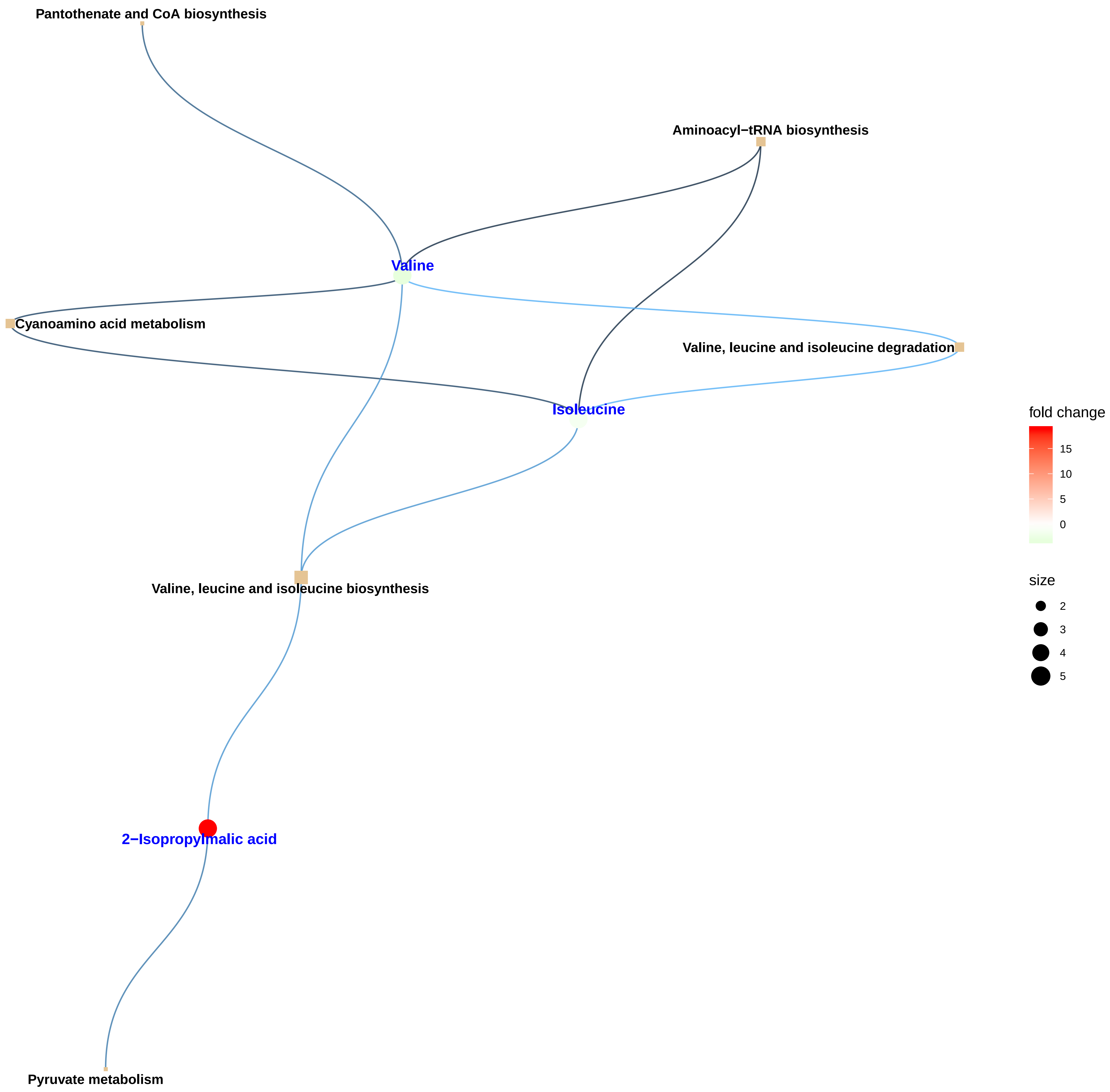

### pathway_network.pdf

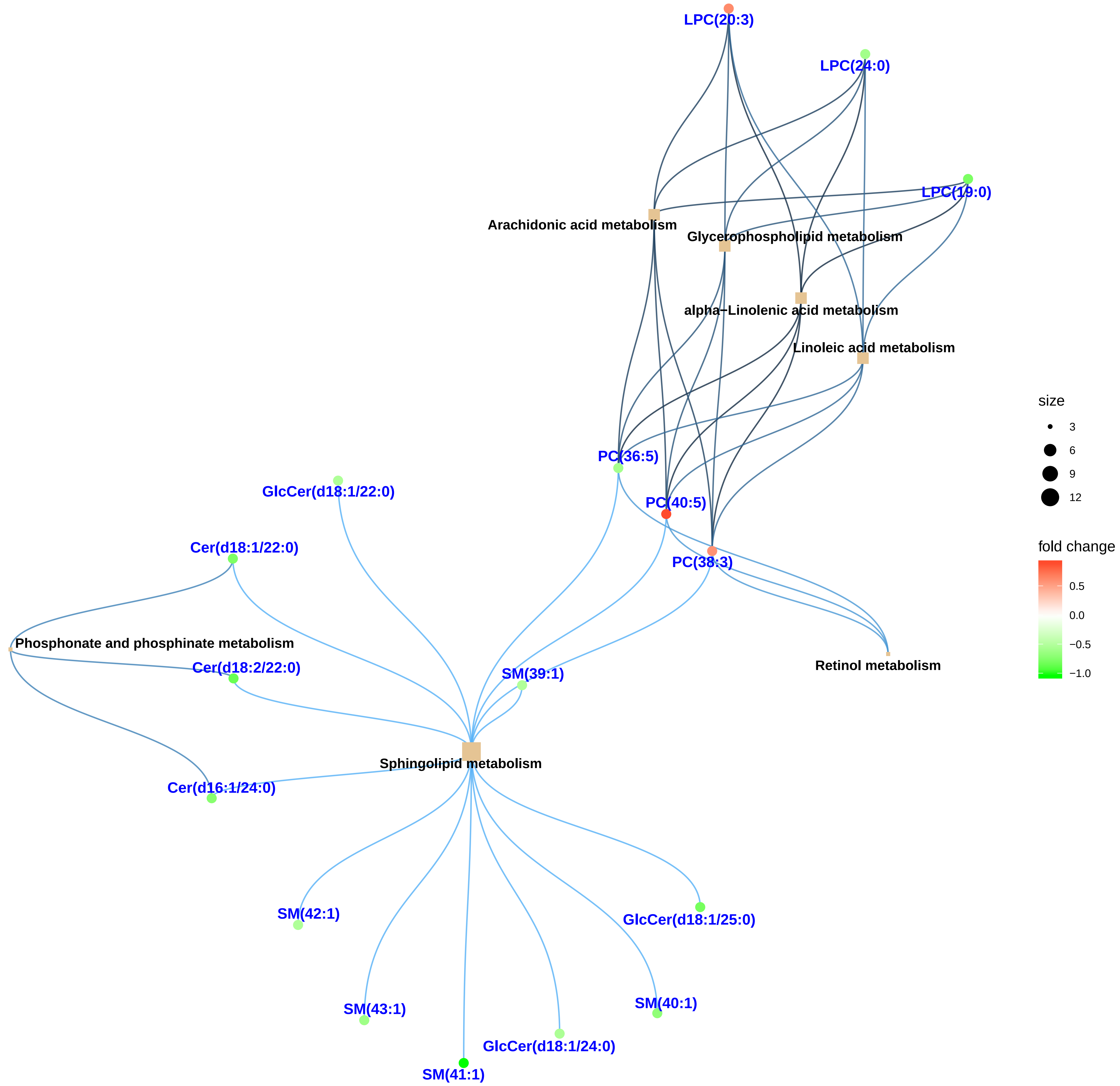

### volcano.pdf

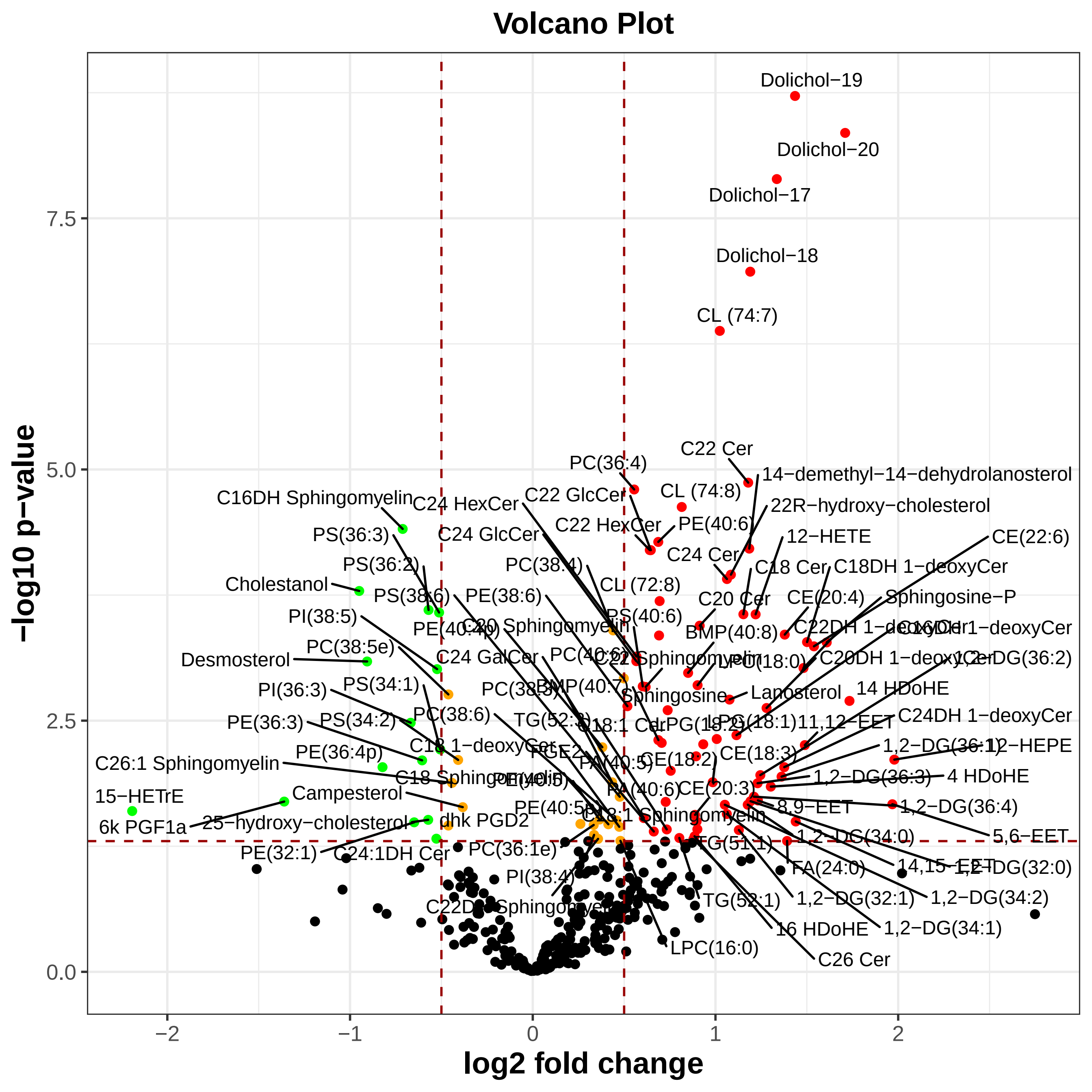

### volcano.pdf

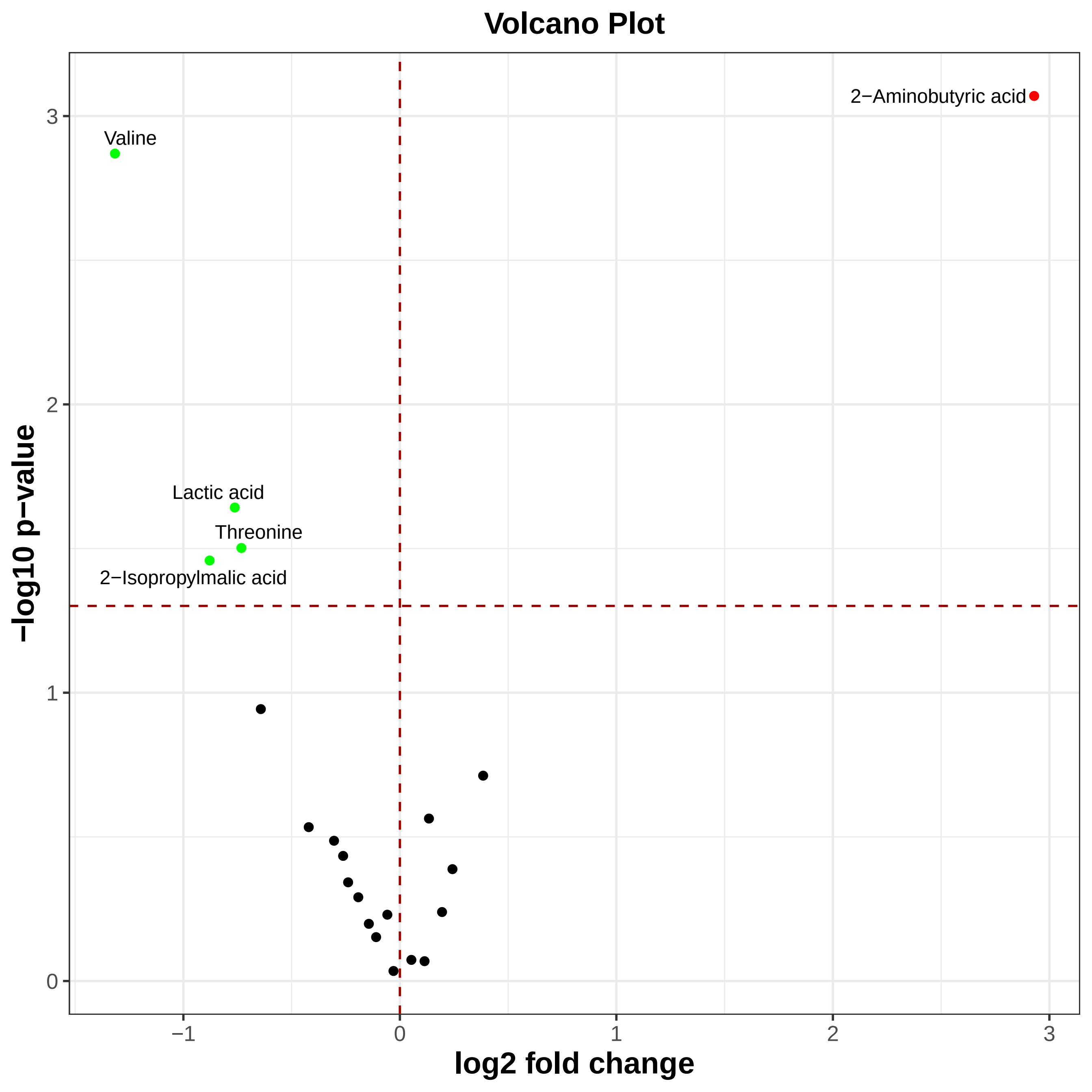

### volcano.pdf

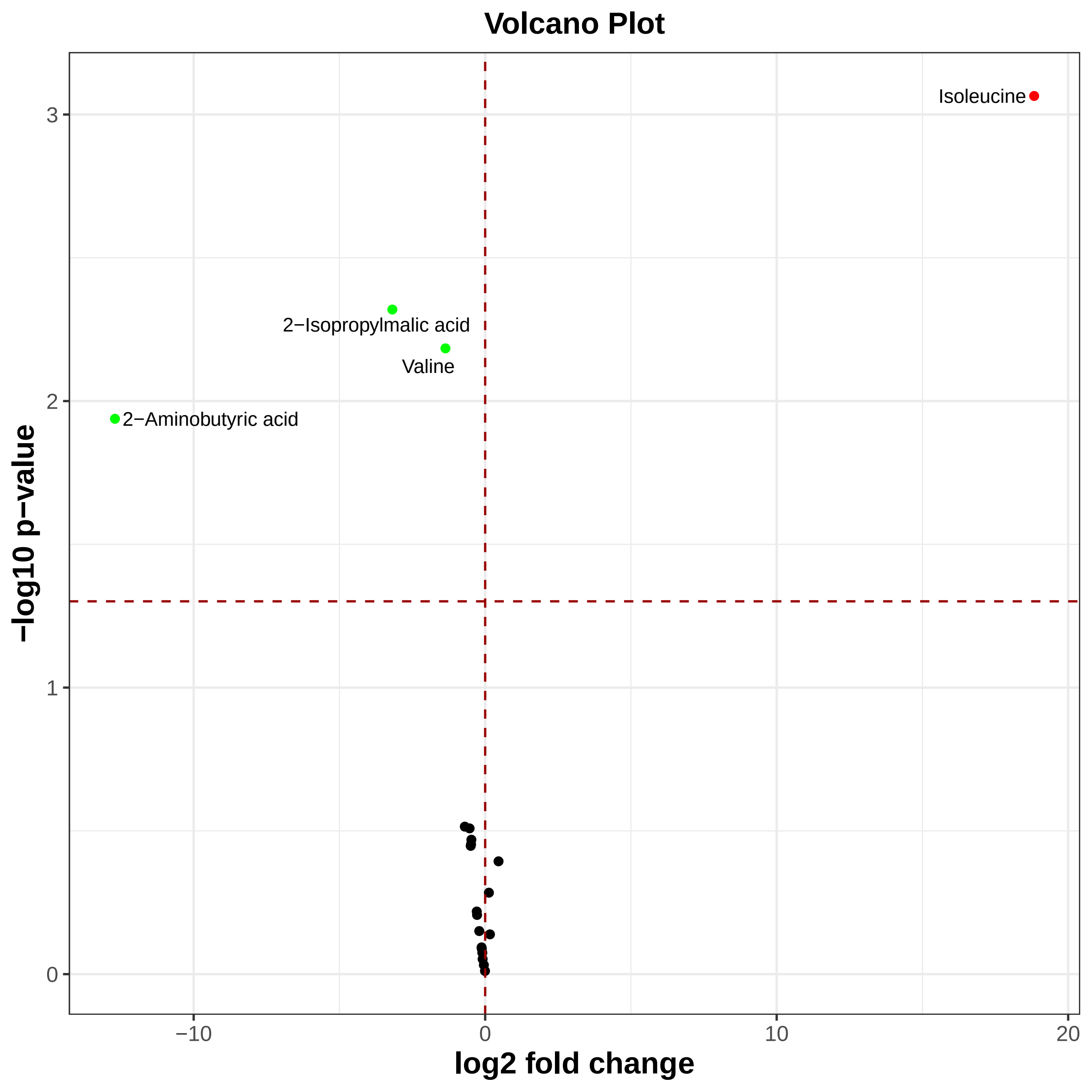

### volcano.pdf

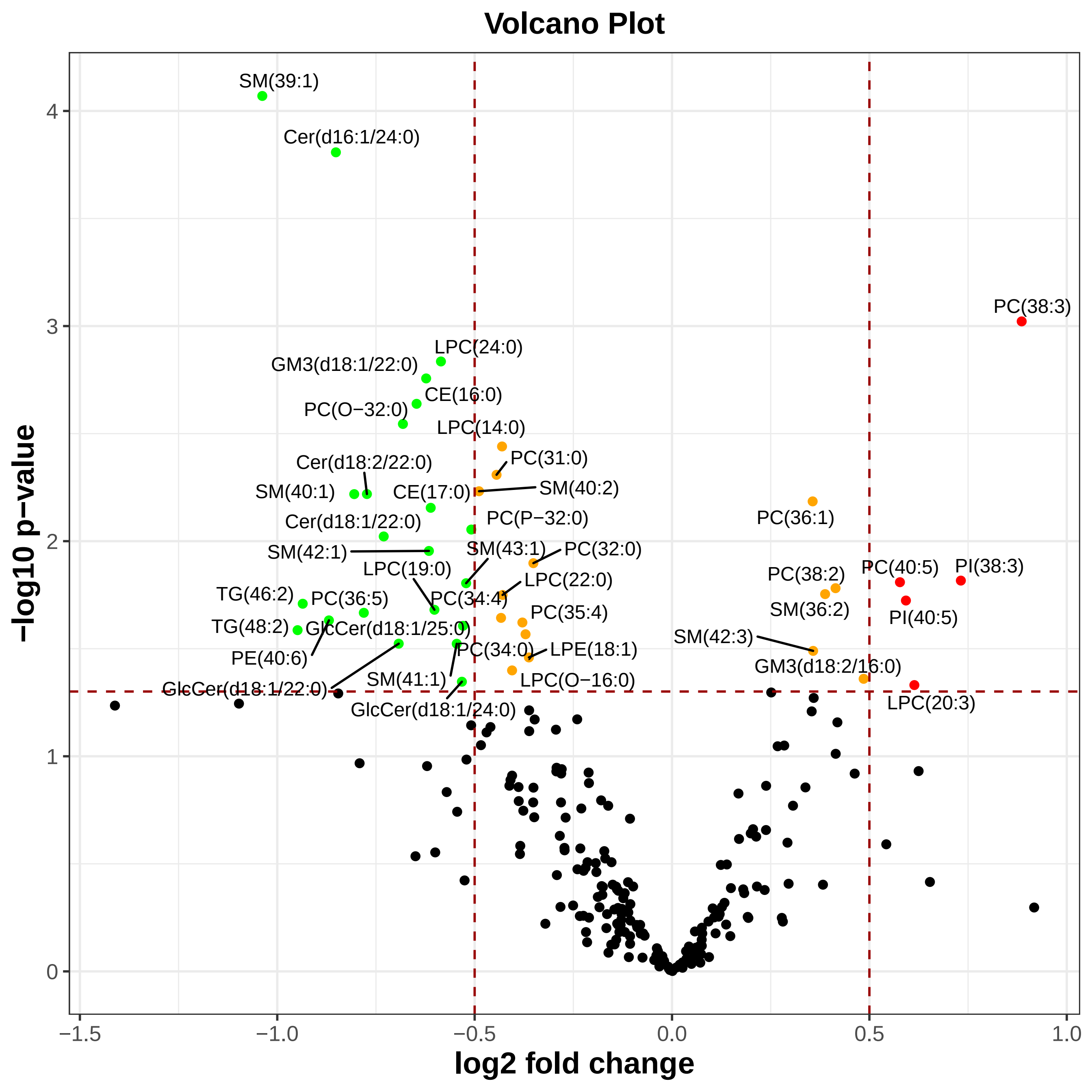
